# Layer-to-cell-class correspondence is gene-specific for pyroptosis-related transcription in human skin

**DOI:** 10.64898/2026.05.03.722547

**Authors:** Daisuke Oryoji, Goro Doi, Sho Fujimoto, Naoya Nishimura, Kyoko Otsuka, Ayako Kuwahara, Masahiro Ayano, Yasutaka Kimoto, Koichi Akashi, Hiroaki Niiro, Hiroki Mitoma

**Affiliations:** Department of Internal Medicine, Kyushu University Beppu Hospital, Beppu, Japan; Department of Medicine and Biosystemic Science, Graduate School of Medical Sciences, Kyushu University, Fukuoka, Japan; Department of Medical Education, Graduate School of Medical Sciences, Kyushu University, Fukuoka, Japan

**Author notes:** Corresponding author Daisuke Oryoji, Department of Internal Medicine, Kyushu University Beppu Hospital 30-10 Soen-cho, Beppu, Oita 874-0838, Japan, Fax: not available.

**Keywords:** spatial transcriptomics, tissue layer, cell class, systemic sclerosis, NLRP1, GSDMD

## Abstract

Anatomical-layer enrichment does not establish cellular attribution, so correspondence with major-cell-class expression must be tested gene by gene. Systemic sclerosis (SSc) skin is informative because inflammasome- and pyroptosis-related molecules have been reported across epidermal, fibroblast-associated and vascular contexts. Spatial transcriptomics of SSc and control skin, with same-donor single-nucleus profiles from 10 SSc donors, revealed gene-specific correspondence. Across 13 discovery sections, *NLRP1*, *PYCARD* and *CASP4* were epidermally biased, whereas *GSDMD* was dermally biased. An independent 10-section SSc cohort supported epidermal *NLRP1* (q = 0.0078) and *PYCARD* (q = 0.043), with directional, non-significant support for *CASP4* and *GSDMD*. Despite epidermal *NLRP1* bias, endothelial class-pseudobulk exceeded keratinocyte class-pseudobulk in all 10 donors. Restricted three-class reconstruction reproduced the *NLRP1* direction in 1 and 0 of 10 donors under the two fixed dermal weightings and in 0–8 of 10 across a post hoc sweep of the dermal endothelial share; *PYCARD* and *GSDMD* were concordant in most donors at every weighting, and *CASP4* in no more than 4 of 10. Thus, tissue layer and major cell class are gene-specifically non-equivalent axes of spatial interpretation.

## Introduction

Spatial transcriptomics preserves tissue architecture but does not make anatomical compartments synonymous with the cells that populate them. A compartment-level gene signal can reflect differences in cell-class abundance, cell-class-specific expression, omitted populations or spatial state variation within a class. Methods that map cell types, deconvolve composition or infer cell-type-specific spatial variation formalize these distinct quantities [1–4], yet biological studies may still move from compartment enrichment to cellular attribution without testing whether separately measured same-donor major-cell-class expression reproduces the observed direction. That correspondence need not be uniform across a pathway: the same tissue may support a composition-based interpretation for one transcript but not for another. We therefore treat layer-to-cell-class correspondence as a gene-level empirical question.

Systemic sclerosis (SSc) provides a structured human setting because fibrosis, vasculopathy and immune dysregulation reshape skin architecture [5], while bulk and single-cell studies either average across compartments or dissociate cells from their native positions [6–8]. A layer-associated signal may therefore reflect cell-class composition, class-specific expression, a combination of both or tissue contexts not represented by the major measured classes. Inflammasome-related transcription spans several skin contexts [9,10]. Human keratinocyte studies identify NLRP1 as a principal epidermal inflammasome sensor [11], whereas SSc work has emphasized NLRP3–caspase-1 activity in dermal fibroblast-associated fibrosis [10]. GSDMD has also been linked to endothelial injury in vascular settings [12–15]. These observations are compatible with a distributed architecture, but they do not establish that a tissue-layer signal can be substituted by the expression ordering of the major cell classes presumed to compose the layer. Spatial transcriptomics makes that attribution testable while retaining compartment geometry [16].

We used public Visium data from 13 human skin discovery sections (4 healthy control and 9 SSc) [17,18] and GSE312129, which contains Visium sections and single-nucleus multiome profiles from the same 10 SSc donors [19,20]. The discovery analysis identified opposing epidermal upstream inflammasome-related and dermal *GSDMD* directions. The paired replication modalities enabled direct gene-level layer testing and a restricted reconstruction of whether major cell-class composition and class-pseudobulk expression reproduced each observed layer direction.

We therefore asked three ordered questions: which pyroptosis-related transcripts have reproducible epidermal or dermal polarity; whether four direct genes retain those directions in GSE312129; and whether a same-donor restricted three-class composition proxy reproduces the observed direction gene by gene. This design separates the existence of a layer signal from the more demanding attribution of that signal to major cell-class context.

## Methods

### Sex as a biological variable

Both female and male participants were included in the source cohorts. In the discovery SSc source cohort, clinical characteristics were reported for 10 SSc participants, of whom 4 (40%) were female; the analytic discovery subset comprised 9 SSc sections because 9 of the 10 attempted SSc repository folders loaded successfully. The SSc5380 section was retained. The replication SSc cohort included 10 sections, of which 7 (70%) were from female participants. Healthy control donors in the discovery cohort were matched to SSc donors in the source study. Because of sample-size constraints, sex was recorded and reported but was not used as a primary stratification variable in the present spatial analyses.

### Study design, cohorts, and ethics

We reanalyzed 13 Visium skin sections from the discovery cohort (4 healthy control [HC], 9 systemic sclerosis [SSc]; Supplementary Table S2) [17–18] and reanalyzed 10 public Whitfield SSc sections (Gene Expression Omnibus (GEO) accession GSE312129) for replication [19–20]. SSc cases fulfilled the 2013 American College of Rheumatology/European League Against Rheumatism (ACR/EULAR) classification criteria [21]. Cohort characteristics are summarized in Table 1. We used the publicly available processed spatial count matrices and associated metadata and did not rerun FASTQ-to-count processing.

**Table 1.** Cohort characteristics.

|  | Discovery cohort |  | Replication cohort |
| --- | --- | --- | --- |
| Characteristic | HC (n = 4) | SSc (clinical n = 10; analytic n = 9*) | SSc (n = 10) |
| Source study | Li et al. ARD 2025 | Li et al. ARD 2025 | Jarnagin et al. JCI Insight 2026 |
| Data repository | Zenodo: 10.5281/zenodo.14577696 | Zenodo: 10.5281/zenodo.14577696 | GEO: GSE312129 |
| Platform | Visium FFPE | Visium FFPE | Visium FFPE |
| Disease subtype | — | Diffuse cutaneous (100%) | Diffuse cutaneous (100%) |
| Age, years | Matched‡ | Median 56 (27–73) | Mean 55.4 (SD 10.5) |
| Female sex, n (%) | Matched‡ | 4 (40%) | 7 (70%) |
| Disease duration, months | — | Median 8 (4–48) | Mean 14.4 (SD 7.7) |
| mRSS | — | Median 21 (8–41) | Mean 22.9 (SD 12.3) |
| ILD, n (%) | — | 4 (40%) | 4 (40%) |
| Biopsy site | Forearm | Forearm | Forearm |
| Sections before QC | 4 | 10 | 10 |
| Sections after QC | 4 | 9* | 10 |
| Total spots after QC | 5,130 | 7,080 | 10,217 |
| Spots/section, median (range) | 1,313.5 (645–1,858) | 672 (453–1,729) | 973.5 (581–1,850) |
| Genes in feature space | 18,061 | 18,061 | 37,082 |

\* Clinical characteristics for the discovery SSc source cohort are reported for the 10 SSc participants in the source study. Loading was attempted for 10 SSc repository folders; 9 loaded successfully, and all 9 loaded SSc sections were retained after the implemented spot- and feature-level preprocessing. The SSc-HL33 repository folder failed to load because spatial/tissue\_positions\_list.csv was unavailable; SSc5380 was retained. All analytic discovery SSc results therefore use 9 loaded sections. ‡ HC donors were age- and sex-matched to SSc donors in the source study.
SSc discovery section IDs: SSc-HL01, SSc-HL05, SSc-HL06, SSc-HL11, SSc-HL13, SSc-HL25, SSc-HL35, SSc4994, SSc5380. HC section IDs: HC01, HC02, HC03, HC05. SSc cases fulfilled the 2013 ACR/EULAR classification criteria [21]. Discovery biopsies were taken from the forearm approximately 15 cm proximal to the styloid process of the ulna.
mRSS, modified Rodnan Skin Score; ILD, interstitial lung disease.

### Spatial preprocessing and inferential framework

After implemented spot- and feature-level preprocessing, Discovery layer labels were generated computationally from eight marker-score panels and a hybrid expression-spatial graph. Standardized spatial coordinates, weighted by two, were combined with the first 10 PCs from 2,000 highly variable genes for neighborhood construction and Leiden clustering. Marker-derived pseudo-labels were refined by multinomial logistic regression using 30 PCs, followed by two spatial-smoothing iterations and one-ring boundary labeling. The resulting layer_v2 and layer_v2_pure fields were used as detailed in the Supplementary Methods. Histology images were displayed for spatial orientation but were not inputs to label generation. The section was the inferential unit, and sectionwise layer contrasts were aggregated across sections to avoid pseudoreplication. Signed effect sizes used Dermis − Epidermis (positive = dermal bias) unless otherwise noted. For direct gene-mean and triad analyses, epidermal and dermal spot means were calculated separately before forming each signed layer difference; the prespecified five-gene replication endpoint instead used compartment medians. No whole-section mean was used. Replication spots were assigned by the maximum of expression-derived marker scores for epidermis (*KRT14*, *KRT5*, *KRT1*, and *KRT10*), dermis (*COL1A1*, *COL1A2*, *COL3A1*, *VIM*, and *DCN*), and endothelium (*PECAM1*, *CDH5*, *VWF*, and *ERG*). The label other was assigned when the maximum score was nonpositive or its margin over the second-highest score was below 0.05. This three-class route did not use the Discovery HVG, PCA, Leiden, pseudo-label, logistic-regression, or spatial-smoothing pipeline and is not an independent histologic validation.

### Replication endpoints, deconvolution, and context analyses

Replication tested related but nonidentical endpoints in 10 SSc sections; discovery comparisons therefore used the 9 SSc sections. In Whitfield, the prespecified dermal endpoint was direct log1p-normalized *GSDMD* expression summarized by compartment mean, and the prespecified epidermal endpoint comprised *PYCARD*, *NLRP3*, *CASP1*, *IL1B*, and *IL18* and was summarized by compartment median. These constructs differed from the discovery *GSDMD*-only and *IL1B*– *IL18* module scores and from the discovery *NLRP1*–*PYCARD*–*CASP4* triad, providing endpoint-level directional support rather than exact construct replication. Cell2location [1], with reference-label initialization by CellTypist [22] and using a Whitfield single-nucleus multiome reference (4 control and 10 SSc participants; 65,066 nuclei) [19], estimated cell-type context at each spot.

The 10 SSc participants contributing the Whitfield reference also contributed matching biopsies for the replication Visium dataset [19]; the replication expression endpoints were calculated directly from the Visium data and did not use deconvolution estimates. Estimated proportions supported dermal *GSDMD* context and epidermal keratinocyte associations. Post hoc sensitivity analysis quantified the partial Spearman association between the implemented 13-gene interferon-gamma-response score and *GSDMD* after adjustment for Cell2location-estimated endothelial relative abundance in the 13 discovery sections (4 HC, 9 SSc). Section-level estimates and methods are reported in Supplementary Table S8 and Supplementary Methods Section 11; additional ancillary context analyses are presented in Supplementary Figs. S10 and S11. Full deconvolution definitions are provided in the Supplementary Methods and Supplementary Table S1. A separate targeted post hoc analysis evaluated the fixed triad in the same 10 sections. Mean log1p-normalized expression was calculated within epidermis and dermis; the signed contrast was Epidermis − Dermis, and the triad was the mean of all three gene-level contrasts. A two-sided one-sample Wilcoxon signed-rank test against zero was exploratory and secondary.

### Direct gene-level replication deepening

For the four fixed direct genes, the GSE312129 primary mask retained only stored epidermis and dermis labels; endothelium and other were excluded. Arithmetic layer means were compared within each section, with expected epidermal direction for *NLRP1*, *PYCARD* and *CASP4* and expected dermal direction for *GSDMD*. The exact two-sided sign test used the section as the inferential unit and BH adjustment across four genes. Descriptive median intervals used 10,000 section-level bootstrap resamples at seed 42. Detection fractions and a dermis-plus-endothelium mask were descriptive sensitivities without additional P values. The fixed donor/sample crosswalk is given in the ReplicationDonorCrosswalk worksheet of the Supporting Data, and all section values are given in Supplementary Table S3.

### Same-donor snMultiome pseudobulk

Raw layers[’counts’] from the GSE312129 snMultiome object were aggregated by donor for keratinocyte, fibroblast and endothelial_strict nuclei. Library size was the sum across 18,703 retained features, CPM was 10^6^ × gene count / library size, and class-pseudobulk expression was log2(CPM+1). The primary SSc analysis required at least 20 nuclei in each class; a floor of 50 was descriptive sensitivity. HC donors were described without formal P values. For each gene, the donor-level contrast compared keratinocyte log2(CPM+1) with the mean of fibroblast and endothelial_strict log2(CPM+1), using the opposite sign for *GSDMD* (Supplementary Table S4). Exact two-sided sign tests were adjusted as a separate BH m=4 family. These are class-pseudobulk values, not per-cell expression.

### Composition adjustment and restricted mixture reconstruction

A discovery sensitivity analysis modeled normalized gene expression as a function of layer and cell2location-estimated keratinocyte, fibroblast and endothelial_strict abundance within each section. The abundance covariates were their posterior medians (q50). The simultaneous route required all predictor VIFs to be at most 10 and the condition number of the standardized predictor matrix to be at most 30; a two-stage route regressed expression on abundance before section-level residual layer summaries. Non-estimability was reported without changing thresholds or models. For reconstruction, observed direction was O_dg=epidermis_mean_log1p−dermis_mean_log1p. Class CPMs were mixed on the linear scale and then converted to log2(CPM+1). M1 defined epidermis as keratinocyte and dermis by the same donor’s fibroblast:endothelial nucleus ratio; M2 used equal 0.5/0.5 dermal weights. The reconstruction was a targeted post hoc analysis of existing outputs; M1 and M2 were fixed in its analysis plan before it was run, and exact sign concordance was primary. A post hoc descriptive sensitivity analysis applied a common fixed endothelial share w of the dermal proxy (0 to 1) to every donor, together with a descriptive reference share (0.0965) computed as endothelial_strict/(fibroblast + endothelial_strict) from discovery dermis-layer mean Cell2location q05 relative abundances, with all other steps unchanged (Supplementary Methods; Supplementary Table S9). Raw cross-platform residuals, percent explained and R-squared were not calculated; leave-one-donor-out was descriptive and used no P value.

### Analysis provenance

Direct replication, composition adjustment and mixture reconstruction used fixed analysis definitions. Full source and row provenance are in the Supplementary Methods and Supporting Data. The snMultiome object was the same reference object previously used for discovery cell2location and is not independent biological validation.

### Generative AI disclosure

ChatGPT and Codex (OpenAI) and Claude (Anthropic) were used to assist with manuscript preparation, development and checking of analysis and plotting code, and review of results. The authors reviewed and verified the outputs and take full responsibility for the analyses, interpretation, and manuscript.

### Statistics and reproducibility

The tissue section was the inferential unit for spatial analyses and the donor for snMultiome analyses. Multiple testing used the Benjamini–Hochberg false discovery rate procedure [23]. The six-gene discovery screen was exploratory and post-selection (BH m=6). The 60.8% dilution was descriptive and post-selection and had no P value. The four-test deconvolution family retained its frozen tests. For the ancillary interferon-gamma–*GSDMD* analysis in Supplementary Fig. S11, the reporting-stage primary was an equal-weight two-sided one-sample Wilcoxon signed-rank test of 13 section-level partial rho values, with the planned Fisher-z pool retained secondarily [24]. Direct four-gene replication used exact two-sided sign tests with BH m=4. SSc pseudobulk used exact two-sided sign tests in a separate fixed four-gene family; HC and floor-50 results were descriptive. The simultaneous composition model was subject to its fixed collinearity criteria, and the two-stage route was reported without rescue. Mixture reconstruction used exact sign concordance without P values, confidence intervals, cross-platform residuals, percent explained or R-squared. The post hoc dermal-weight sensitivity analysis was descriptive and used no P values. Leave-one-donor-out results were descriptive. All formal tests were two-sided with nominal alpha 0.05. No a priori sample-size calculation was performed because public sections and donors were fixed. No spot-level P values were used.

### Ethics approval

Ethics approval was not required for this secondary analysis because it used only de-identified public data. The original discovery study by Li et al. was approved by the ethics committees of Huashan Hospital, Fudan University, was conducted in accordance with the Declaration of Helsinki, and obtained written informed consent from all participants [17]. The replication source study was approved by the University of Michigan Institutional Review Board; all participants gave written informed consent, and the Committee for the Protection of Human Subjects at Dartmouth College’s Geisel School of Medicine approved all human-participant analyses [19].

## Results

### Discovery identifies opposing tissue-layer polarity

To define the spatial organization of pyroptosis-related transcription across human skin, we analyzed 13 loaded Visium FFPE sections from the discovery cohort, comprising 4 healthy control (HC) and 9 systemic sclerosis (SSc) sections. All 13 loaded sections were retained after the implemented spot- and feature-level preprocessing. The analysis used sectionwise epidermal-versus-dermal comparisons rather than whole-section averaging. A compact study schematic and representative maps from HC03, SSc4994, SSc-HL35, and SSc-HL06 revealed an immediate split between the adaptor *PYCARD* and the gasdermin effector *GSDMD*: *PYCARD* signal was concentrated toward the epidermal compartment, whereas *GSDMD* signal was concentrated toward the dermis. This divergence was visible in both HC and SSc sections, indicating that the pattern is not restricted to diseased skin, while many SSc sections exhibited stronger visual separation than HC sections (Fig. 1A,B; Supplementary Figs. S1 and S2).

**Figure 1.**
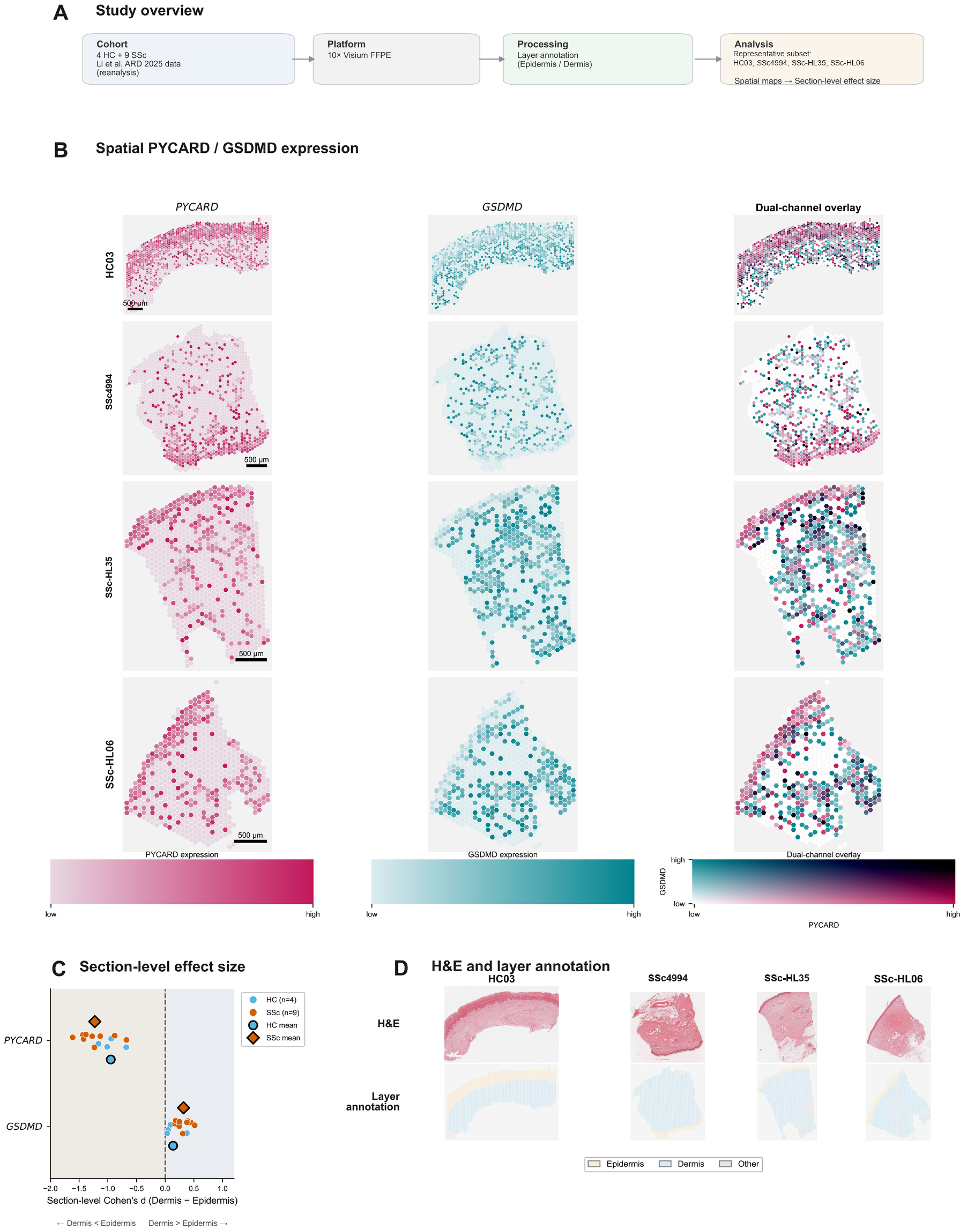
Spatial mapping displays divergent epidermal *PYCARD* and dermal *GSDMD* enrichment across the discovery cohort. (A) Compact study design schematic summarizing the discovery cohort (reanalysis of Li et al. [17]), Visium FFPE platform, computational layer-label workflow, and representative subset selection. (B) Representative spatial maps from HC03, SSc4994, SSc-HL35, and SSc-HL06 showing single-channel *PYCARD* and *GSDMD* expression together with a dual-channel composite view from the same sections. (C) Sectionwise Cohen’s d values for Dermis − Epidermis across all 13 discovery sections; negative values indicate epidermal enrichment and positive values indicate dermal enrichment, with cohort summaries shown for HC and SSc. (D) H&E thumbnails are shown for spatial orientation and were not used to assign layers; the paired computational label thumbnails show epidermis, dermis, and unassigned or other regions used for sectionwise comparisons. Discovery cohort: 13 loaded sections (4 HC and 9 SSc). Supplementary Figs. S1 and S2 extend the single-channel *PYCARD* and *GSDMD* maps to all discovery sections. Panel C uses the exact preserved Figure 1C plotting export derived from layer_v2_pure, which retains only Epidermis and Dermis spots and excludes Boundary, Adipose, Other, and missing labels. Figure 2A direct-gene analyses and later module-score analyses use the broader layer_v2 assignment, so their spot-count denominators are not interchangeable.

Sectionwise effect size summaries reproduced the same directional split across the full discovery cohort. Using Cohen’s d defined as Dermis − Epidermis, negative values indicate epidermal enrichment and positive values indicate dermal enrichment. Under that convention, *PYCARD* was epidermally biased in 13 of 13 discovery sections, whereas *GSDMD* was dermally biased in 13 of 13 sections. The cohortwise dumbbell summary therefore demonstrated complete directional concordance for both genes, despite variation in absolute magnitude from section to section. The plotted group pattern was primarily one of amplitude rather than direction: HC sections already exhibited the split, whereas SSc sections tended to extend farther toward epidermal *PYCARD* and dermal *GSDMD* values. These findings establish a reproducible organization across two compartments at the mRNA level, with upstream adaptor signal and downstream gasdermin signal occupying different spatial layers (Fig. 1C; Supplementary Figs. S1 and S2).

Layer annotations were assigned at spot level and are visualized in the boundary thumbnails (Fig. 1D). In sum, Fig. 1 establishes a cohortwide split, with *PYCARD* enriched in the epidermis and *GSDMD* in the dermis across HC and SSc sections, and the atlas supports representative selection (Supplementary Figs. S1 and S2).

We asked whether the epidermal signal seen for *PYCARD* reflected a broader inflammasome-related program or a more selective upstream module. *CARD8* and *CASP1* had been excluded during probe filtering and were absent from the filtered discovery FFPE feature space, leaving six evaluable genes. In this exploratory post-selection screen, signed epidermal consistency identified *PYCARD*, *NLRP1*, and *CASP4* as epidermally biased in 13 of 13 discovery sections, whereas the remaining surveyed genes were weaker, mixed, or oppositely directed. *NLRP3* followed the inverse pattern, with dermal predominance in 12 of 13 sections and HC01 as the lone epidermal exception. Representative spatial maps of *PYCARD*, *NLRP1*, and *CASP4* across the same four sections used in Fig. 1 showed coherent epidermal concentration, while sectionwise direction maps and concordance summaries extended that observation across the full cohort. We therefore refer descriptively to a direction-concordant epidermal *NLRP1*–*PYCARD*– *CASP4* triad rather than to a generic feature of all inflammasome-related genes (Fig. 2A,B; Supplementary Figs. S3, S4, and S5).

**Figure 2.**
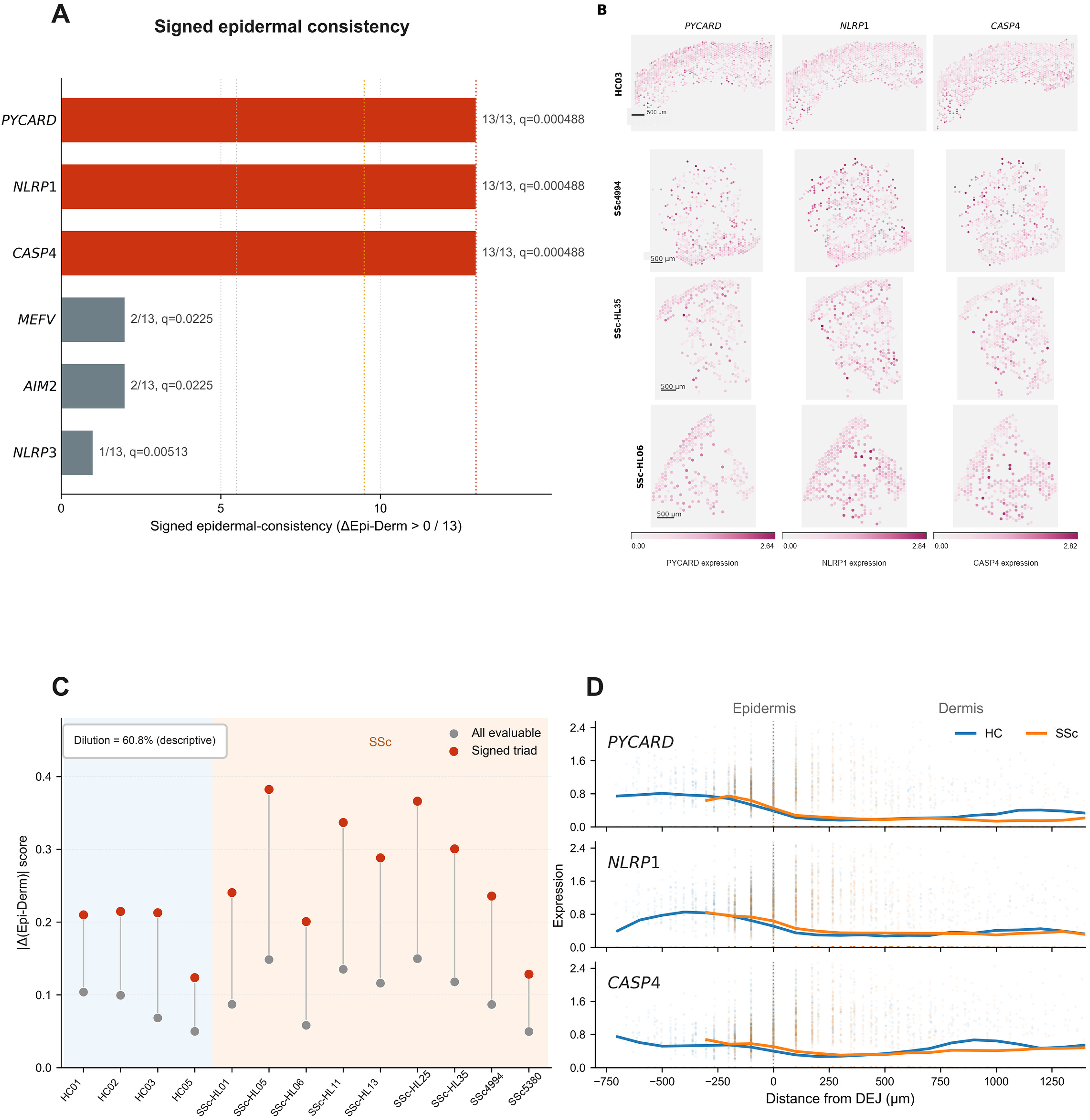
Upstream inflammasome components show consistent epidermal enrichment with descriptive signal near the computed layer interface. (A) Bars show the number of sections with Δ(Epidermis − Dermis) > 0 among the 13 discovery sections for the six evaluable inflammasome-related genes; annotations give the corresponding n/N and BH-adjusted q values. The six-gene direct-expression screen used the broader layer_v2 assignment, with sectionwise arithmetic means calculated separately for Epidermis and Dermis before forming Δ(Epidermis − Dermis). *PYCARD*, *NLRP1*, and *CASP4* were directionally concordant across all sections, whereas comparator genes were weaker or oppositely directed. The BH-adjusted q values shown in panel A are exploratory post-selection summaries across the six evaluable genes; the underlying exact sign-test P values are provided in the Supporting Data workbook. (B) Representative spatial maps from HC03, SSc4994, SSc-HL35, and SSc-HL06 showing epidermal concentration of *PYCARD*, *NLRP1*, and *CASP4*. (C) Sectionwise comparison of a broader inflammasome-related composite versus the signed *NLRP1*–*PYCARD*–*CASP4* triad, illustrating dilution of the layer-specific signal in the broader score. Dilution was calculated as 1 – mean(|delta_broader|) / mean(|delta_signed_triad|). Because the triad was defined in these same discovery sections, the 60.8% dilution was treated as a descriptive post-selection metric and no formal P value was assigned. (D) Descriptive signed-distance profiles for *PYCARD*, *NLRP1*, and *CASP4* individually, using 100-μm bins relative to computational boundary-anchor spots. Faint points are individual spots; cohort lines show smoothed pooled-spot bin means, as defined in Supplementary Methods Section 12. These are individual-gene profiles, not a triad-composite curve. Signed distance is derived from computed boundary labels with pitch scaling for each section and is not an anatomical contour segmented from histology. Panel C uses the frozen section-level contrasts from the Figure 2C source table; this source-specific contrast is distinct from the separate direct gene-mean triad metric in Supplementary Fig. S7.

Separate discovery endpoint summaries sharpened that selectivity without representing the Whitfield constructs. The *IL1B*–*IL18* cytokine-maturation module score, summarized by compartment median, was epidermally directed in all 13 discovery sections; the *GSDMD*-only module score, summarized by compartment mean, was dermally directed in all 13. The exploratory six-gene heatmap likewise showed that not every screened gene tracked with the direction-concordant epidermal triad, with *MEFV* and *AIM2* predominantly dermal rather than epidermal. These discovery summaries support treating the pathway-related transcription as spatially structured rather than as a single spatially undifferentiated score (Supplementary Figs. S3, S5, and S6).

### Pathway selectivity and composite dilution

The descriptive dilution metric was calculated in the panel comparing composites. When the broader composite using all available genes was compared with the direction-concordant epidermal triad, the broader score yielded weaker layer separation, consistent with dilution by genes that were not directionally concordant with the triad. Using the signed *NLRP1*–*PYCARD*– *CASP4* triad, the descriptive post-selection dilution metric relative to the broader inflammasome-related composite was 60.8%. Because the triad was defined from directional consistency in these same discovery sections, no formal inferential P value was assigned to this comparison. In practical terms, the most spatially coherent upstream signal was recovered when the analysis was restricted to the direction-concordant triad rather than expanded to a broader gene set with mixed directions (Fig. 2C).

To assess where this epidermal triad was positioned relative to the computed layer interface, we added a descriptive signed-distance analysis using computationally generated boundary-anchor spots and pitch scaling for each section. Across HC and SSc groups, *PYCARD*, *NLRP1*, and *CASP4* all peaked on the epidermal side of the anchor set and declined across the approximate layer interface into the dermis. Because this distance metric is derived from computed boundary labels rather than a continuous contour segmented from histology, we interpret the curves descriptively rather than as a precise anatomical reconstruction. Even with that caution, the panel supports the same directional message as the sectionwise analyses. In a descriptive compact-set comparison, the pyroptosis-related upstream-effector split was larger than those of apoptosis and necroptosis; this provides pathway context but does not establish pathway specificity (Fig. 2D; Supplementary Fig. S7).

### Direct four-gene replication establishes gene-specific layer directions

We evaluated four direct genes in the 10 GSE312129 Visium sections. The primary contrast retained stored epidermis and dermis labels only; 270 endothelium-labelled and 909 other-labelled spots were excluded, a definition that is not symmetric with discovery layer_v2_pure because the discovery route has no corresponding endothelial class. In the planned descriptive sensitivity that added the endothelium-labelled spots to dermis, the number of sections in the expected direction was unchanged for all four genes (NLRP1 10, PYCARD 9, CASP4 7 and GSDMD 8 of 10; Supporting Data). With section as the inferential unit and BH adjustment across four fixed genes, *NLRP1* was epidermally directed in 10 of 10 sections (q=0.0078), *PYCARD* in 9 of 10 (q=0.043), *CASP4* in 7 of 10 (q=0.34), and *GSDMD* was dermally directed in 8 of 10 (q=0.15). Direction counts are the primary interpretive context; the *CASP4* and *GSDMD* q values were non-significant (Fig. 3; Supplementary Table S3). Earlier related but nonidentical endpoints remain in Supplementary Figs. S8 and S12 and Supplementary Results.

**Figure 3.**
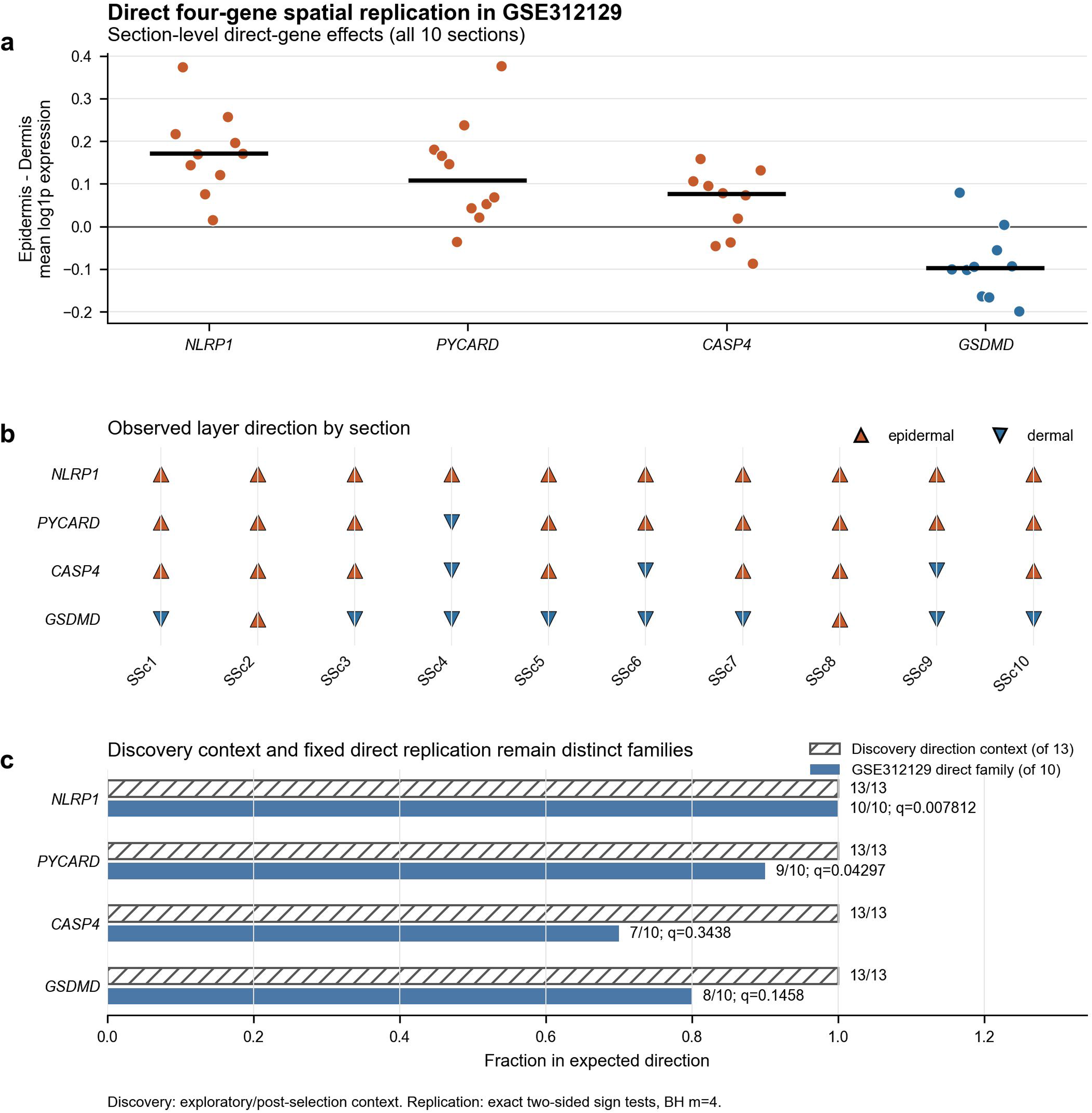
Direct four-gene spatial replication in GSE312129. (a) Epidermis-minus-dermis mean log1p expression for all 10 sections and four direct genes. Points show sections and black bars medians; positive values are epidermal. (b) Direction matrix showing every section. (c) Discovery direction context and the fixed direct replication family are displayed separately. Discovery counts are exploratory/post-selection context. Replication used four fixed direct endpoints, exact two-sided sign tests and BH m=4. *NLRP1* was epidermal in 10 of 10 (q=0.0078), *PYCARD* in 9 of 10 (q=0.043), *CASP4* in 7 of 10 (q=0.34), and *GSDMD* was dermal in 8 of 10 (q=0.15). The *CASP4* and *GSDMD* q values were non-significant. The primary mask included stored epidermis and dermis only and excluded 270 endothelium-labelled and 909 other-labelled spots.

### Same-donor major-cell-class context is concordant for some genes but not others

We summarized raw-count snMultiome expression as donor-level pseudobulk for keratinocyte, fibroblast and endothelial_strict nuclei, using the 18,703 retained-feature library denominator and a primary floor of at least 20 nuclei in each class. Among the 10 SSc donors, *PYCARD* was higher in keratinocyte than fibroblast in 10 of 10 donors (median K−F=+1.738 log2[CPM+1]) and higher than endothelial_strict in 9 of 10 (median K−E=+1.597). *GSDMD* was higher in fibroblast and endothelial_strict than keratinocyte in all 10 donors (median K−F=−1.925; K−E=−3.184). These genes showed class-pseudobulk ordering concordant with their observed layer directions (Fig. 4; Supplementary Table S4).

**Figure 4.**
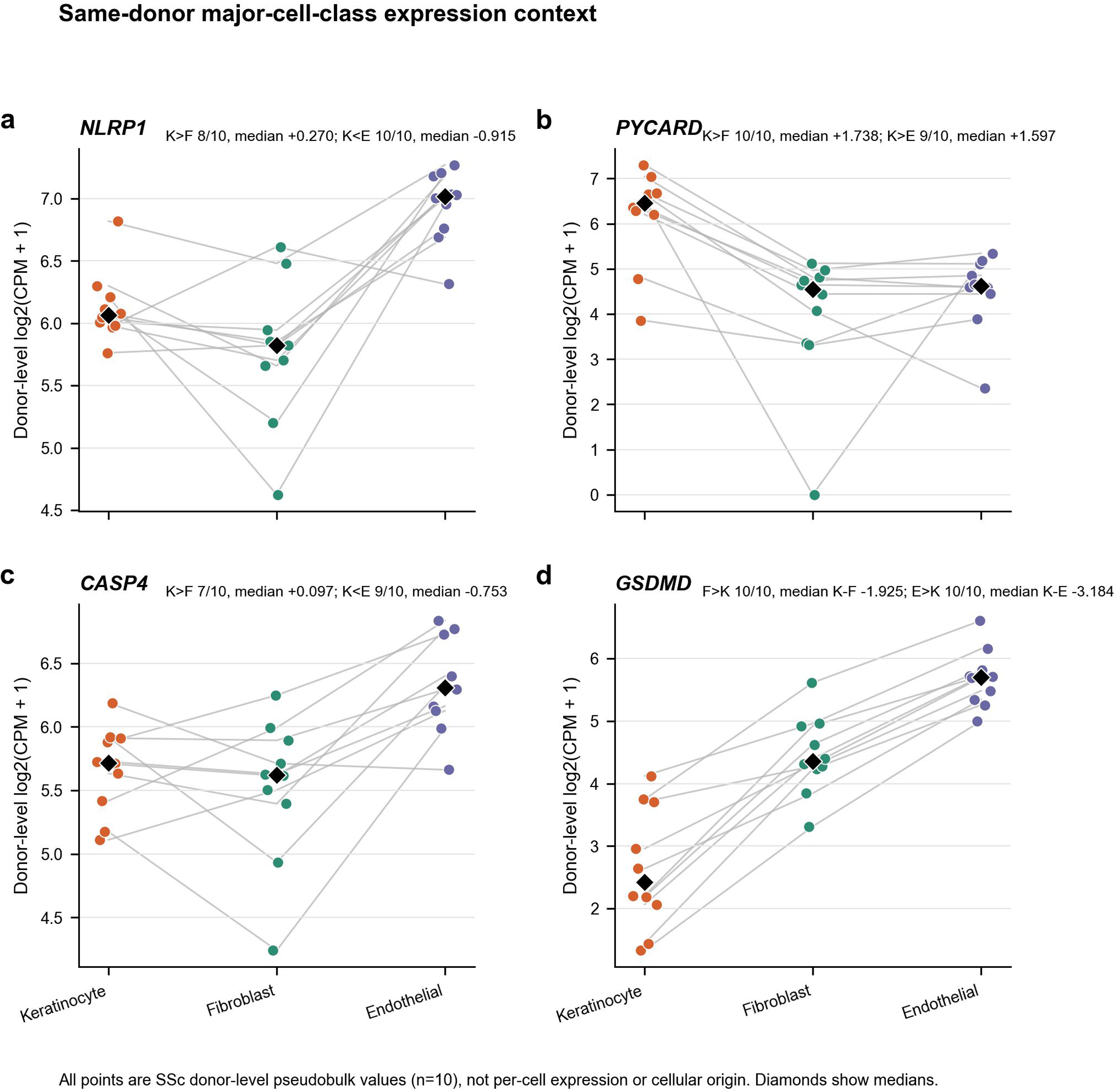
Same-donor major-cell-class expression context. Donor-level log2(CPM+1) pseudobulk for keratinocyte, fibroblast and endothelial_strict in 10 SSc donors is shown for (a) *NLRP1*, (b) *PYCARD*, (c) *CASP4* and (d) *GSDMD*. Grey lines connect classes within donors; colored points show every donor and black diamonds show medians. Contrast annotations give frozen directional counts and median K−F/K−E values. Raw counts used the 18,703-retained-feature library denominator and the primary floor of at least 20 nuclei in each class. These values describe donor-level class context, not per-cell expression or cellular origin.

The other genes did not support a simple epidermis-equals-keratinocyte attribution. *NLRP1* was higher in keratinocyte than fibroblast in 8 of 10 donors (median K−F=+0.270) but lower than endothelial_strict in all 10 (median K−E=−0.915). *CASP4* was higher in keratinocyte than fibroblast in 7 of 10 (median K−F=+0.097) but lower than endothelial_strict in 9 of 10 (median K−E=−0.753). In the fixed, targeted post hoc same-donor contrasts on the log2(CPM+1) scale (keratinocyte minus the equal-weight fibroblast–endothelial_strict mean for NLRP1, PYCARD and CASP4; the reverse sign for GSDMD), PYCARD and GSDMD followed their layer directions in 10 of 10 donors (q = 0.0039 each) and CASP4 in 3 of 10 (q = 0.34), whereas NLRP1 ran opposite to its epidermal layer direction in 9 of 10 (q = 0.029). Descriptive pooled K/F/E CPM across the broader 10-SSc-plus-4-HC scope was 70.3/61.4/130.4 for *NLRP1*, 76.3/26.1/26.2 for *PYCARD*, 47.6/49.9/82.9 for *CASP4*, and 6.4/23.1/54.6 for *GSDMD*. These are class-pseudobulk contexts, not per-cell expression or cellular-origin assignments (Fig. 4; Supplementary Table S4).

### Collinearity motivates forward direction reconstruction

A frozen discovery sensitivity attempted simultaneous adjustment for q50 keratinocyte, fibroblast and endothelial abundance. The maximum variance inflation factor was 15.5, and 5 of 13 sections failed the fixed collinearity gate; the simultaneous route was therefore NOT_ESTIMABLE_COLLINEARITY. A two-stage residual route yielded BH q=1.0 for all four genes, but this estimator is attenuation-prone and does not test the absence of layer polarity.

At Visium resolution, layer and keratinocyte abundance were not statistically separable by this regression design. We therefore used forward restricted reconstruction to ask a different directional question, conditional on a stated dermal weighting, without changing the frozen thresholds or models (Supplementary Table S5).

### Restricted same-donor reconstruction reveals gene-specific correspondence

For each gene and donor, the observed sign was the Visium epidermis-minus-dermis direction. The restricted three-class proxy treated epidermis as 100% keratinocyte; M1 mixed fibroblast and endothelial_strict CPM for dermis according to the same donor’s nucleus ratio, whereas M2 used equal fibroblast:endothelial weights. Class CPMs were mixed on the linear scale before conversion to log2(CPM+1). The primary comparison was exact sign concordance, not cross-platform magnitude or a raw residual. The proxy is not a measurement of Visium spot composition (Fig. 5a,b; Supplementary Tables S6 and S7).

**Figure 5.**
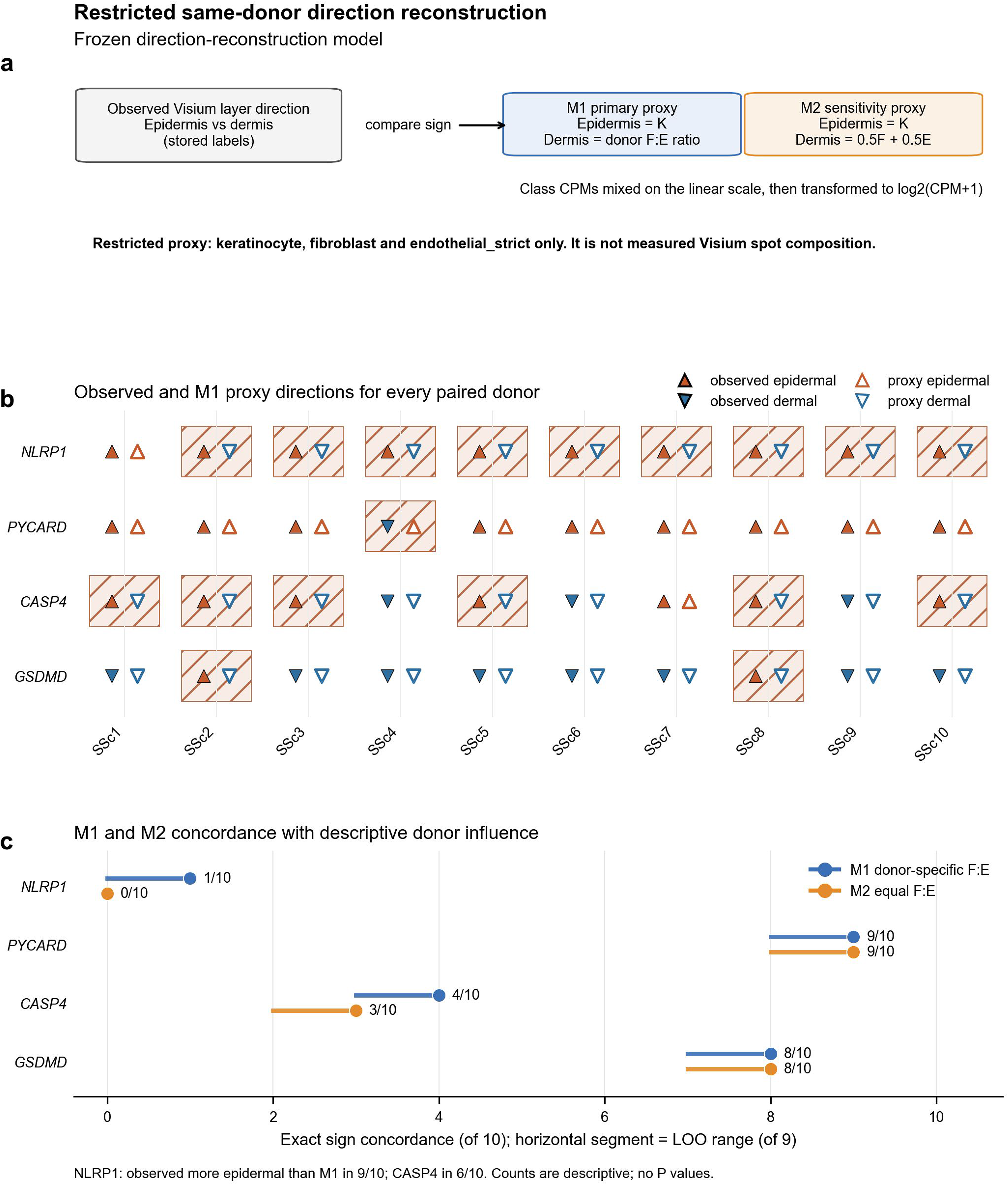
Restricted same-donor direction reconstruction. (a) Frozen model schematic. Observed Visium epidermis-versus-dermis direction was compared with a restricted proxy in which epidermis was keratinocyte; dermis used the same donor’s fibroblast:endothelial nucleus ratio in M1 and equal fibroblast:endothelial weights in M2. Class CPMs were mixed on the linear scale before log2(CPM+1). The proxy is restricted to three classes and is not measured Visium spot composition. (b) Complete observed-versus-M1 direction matrix for four genes and all 10 paired donors. Filled markers are observed directions, open markers proxy directions, and hatched cells denote non-reproduction. (c) Exact sign concordance under M1 and M2 with descriptive leave-one-donor-out ranges. Concordance was M1/M2 1/0 of 10 for *NLRP1*, 9/9 for *PYCARD*, 4/3 for *CASP4* and 8/8 for *GSDMD*. *NLRP1* was OBSERVED_MORE_EPIDERMAL_THAN_PROXY in 9 of 10 donors under M1 and *CASP4* in 6 of 10. Post hoc concordance across dermal endothelial shares from 0 to 1 is given in Supplementary Table S9. Counts are descriptive; no P values, raw cross-platform residuals, percent explained or R-squared were calculated.

Directional correspondence was gene-specific. For *NLRP1*, the proxy reproduced the observed sign in 1 of 10 donors under M1 and 0 of 10 under M2; M1 classified 9 of 10 as OBSERVED_MORE_EPIDERMAL_THAN_PROXY. For *PYCARD*, concordance was 9 of 10 under both models. For *CASP4*, it was 4 of 10 under M1 and 3 of 10 under M2, with OBSERVED_MORE_EPIDERMAL_THAN_PROXY in 6 of 10 under M1. For *GSDMD*, concordance was 8 of 10 under both models. Leave-one-donor-out ranges were 0–1 of 9 and 0 of 9 for *NLRP1* (M1/M2), 8–9 of 9 for *PYCARD* under both models, 3–4 of 9 and 2–3 of 9 for *CASP4*, and 7–8 of 9 for *GSDMD* under both models (Fig. 5c; Supplementary Tables S6 and S7).

Under both fixed weightings, the restricted proxy therefore reproduced most *PYCARD* and *GSDMD* directions, was intermediate for *CASP4*, and reproduced the observed epidermal *NLRP1* direction in at most 1 of 10 donors. Because neither weighting is calibrated to dermal tissue composition, a post hoc sensitivity analysis varied the endothelial share of the dermal proxy from 0 to 1 (Supplementary Table S9). Across this range, concordance was 9–10 of 10 for *PYCARD* and 8 of 10 for *GSDMD*, consistent with keratinocyte class-pseudobulk being the highest of the three modelled classes for *PYCARD* in 9 of 10 donors and the lowest for *GSDMD* in all 10; *CASP4* remained at 2–4 of 10, although different donors were concordant at low and high endothelial shares; and *NLRP1*, whose endothelial class-pseudobulk exceeded keratinocyte in every donor, was concordant in 8, 7 and 7 of 10 at shares of 0, 0.05 and 0.10, in 4 of 10 at 0.20 and in none at 0.5 or more. At the share implied by discovery-cohort Cell2location dermal abundances (0.0965; a cross-cohort descriptive reference, not a composition estimate for the replication sections), concordance was 7, 9, 2 and 8 of 10 for *NLRP1*, *PYCARD*, *CASP4* and *GSDMD*. Because the observed replication dermis excluded endothelium-labelled spots whereas the proxy dermis includes endothelial expression at any nonzero share, the sweep describes sensitivity to this assumption rather than resolving it. *PYCARD* and *GSDMD* are descriptive comparators rather than inferential controls. Directional non-reproduction does not prove a layer-intrinsic program, keratinocyte upregulation or cellular origin.

### Endothelial and interferon-gamma context

The frozen four-test deconvolution family remained a formal main-text result although its display moved to Supplementary Fig. S13. P1 linked dermal *GSDMD* with estimated endothelial context (meta rho=0.181; positive in 13 of 13 sections; q=2.9×10^−7^). P2 linked *GSDMD* inversely with estimated fibroblast context (meta rho=−0.166; negative in 12 of 13; q=9.5×10^−5^). P3 linked the epidermal inflammasome device score with estimated keratinocyte context (meta rho=0.098; positive in 10 of 13; q=0.018). P4 showed lower estimated endothelial abundance in the high-fibrosis-score half in 12 of 13 sections (high minus low negative; exact two-sided sign-test P=0.0034; q=0.0046). The q values belong to the frozen section-level test family, whereas the meta rho values are secondary Fisher-z summaries. These are inferred tissue-context associations, not direct cell counts or cellular-origin assignments (Supplementary Figs. S9 and S13).

In a post hoc sensitivity analysis, the dermal IFN-γ-response-score–*GSDMD* association remained small and positive after adjustment for Cell2location-estimated endothelial relative abundance (median sectionwise partial rho=0.065; 95% section-bootstrap CI 0.034–0.114; positive in 12 of 13 sections; two-sided Wilcoxon P=0.00049; Supplementary Table S8).

### Layer and major cell class are gene-specific interpretive axes

The discovery, direct replication, class-pseudobulk and reconstruction results distinguish two analytical steps. Gene-set aggregation can dilute opposing layer directions, while assignment of a layer signal to the expression of its presumed cell-class composition can succeed for some genes, fail for others, or depend on a composition that the data do not identify. Layer-to-cell-class correspondence is gene-specific, and therefore cellular attribution of a spatial compartment signal must be tested gene by gene rather than assumed. This conclusion is directional and mRNA-level and does not assign cellular origin or mechanism (Figs. 1–5; Supplementary Fig. S14; Supplementary Tables S3–S7 and S9).

## Discussion

The principal advance is that pyroptosis-related transcription in human skin is organized along two related but non-equivalent axes: anatomical layer and major cell class. At the tissue level, *NLRP1*, *PYCARD* and *CASP4* formed a recurrent epidermal pole, opposed by dermal *GSDMD*, across all 13 discovery sections. Direct testing in GSE312129 supported epidermal *NLRP1* and *PYCARD* after adjustment across four fixed genes, while *CASP4* and *GSDMD* retained the expected directions without significant adjusted support. Despite its recurrent epidermal tissue direction, *NLRP1* class-pseudobulk was higher in endothelial_strict than in keratinocyte in all ten SSc donors. In the same ten SSc donors and under the fixed weightings, the restricted three-class proxy reproduced most *PYCARD* and *GSDMD* directions, was intermediate for *CASP4* (4 of 10 under M1 and 3 under M2), and reproduced the epidermal *NLRP1* direction in only one donor under M1 and none under M2; in the post hoc sweep, *NLRP1* concordance ranged from 8 of 10 with a fibroblast-only dermis to 0 of 10 at endothelial shares of 0.5 or more (Supplementary Table S9). The tissue-level architecture was therefore directionally recurrent, but its translation into major-cell-class expression was gene-specific. This gene specificity is the central biological result. The class ordering beneath it did not depend on the dermal weighting: among the three modelled classes, keratinocyte class-pseudobulk was the highest for *PYCARD* in 9 of 10 donors and the lowest for *GSDMD* in all 10, whereas for *NLRP1* it was never the highest and lay between fibroblast and endothelial values in 8 of 10 donors. Proxy concordance for *NLRP1*, by contrast, depended on the endothelial share assumed for the dermis, whereas the separation between PYCARD and GSDMD (at least 8 of 10 at every share) and CASP4 (at most 4 of 10 at every share, although its replication direction was itself weaker) did not (Supplementary Table S9). Under the fixed weightings, the four genes formed a spectrum from frequent correspondence, through intermediate correspondence, to non-reproduction.

Two distinct information-loss steps follow. First, pathway aggregation can combine genes with different or opposing layer polarities: relative to the direction-concordant *NLRP1–PYCARD– CASP4* triad, the broader inflammasome-related composite showed 60.8% descriptive post-selection dilution, while the separate *GSDMD* endpoint occupied the opposing dermal pole. Second, replacing an anatomical layer with the expression program of its presumed cell-class composition can preserve some gene directions while failing to reproduce others or making their reproduction contingent on a composition that the data do not identify. These are not redundant limitations. The first removes gene-level direction within a pathway-related set; the second removes anatomical information when a compartment is represented only by the expression of its presumed constituent cell classes. In SSc skin, where epidermal stress, vasculopathy, inflammation and fibrotic remodeling coexist, the consequence is biological as well as analytical: a single pyroptosis-related score or a single compartment-to-cell-class assignment can collapse a distributed epidermal–dermal architecture into an apparently coherent program. The compact apoptosis and necroptosis comparisons provide contextual support but, because their component sets differ, do not establish pathway specificity. *NLRP1* provides the clearest gene-level example of the second information-loss step: its layer direction was the most consistently replicated of the four genes (10 of 10 sections; q = 0.0078), yet among the three modelled classes keratinocyte class-pseudobulk was never its highest, and whether class-level expression reproduced that direction depended on the dermal composition assumed (Supplementary Table S9).

The *NLRP1* result is biologically compatible with, but does not prove, known epidermal inflammasome biology. Human keratinocyte studies identify NLRP1 as an epidermal sensor [11], whereas SSc work has emphasized NLRP3 in dermal fibroblast-associated fibrosis [10]. Consistent with those contexts, *NLRP1* was epidermal in 13 of 13 discovery sections and *NLRP3* was dermal in 12 of 13. The central outcome genes *NLRP1*, *PYCARD*, *CASP4* and *GSDMD* were outside the 2,000 highly variable gene set used for discovery PCA. By contrast, *NLRP3* and *IL1B* were highly variable genes, so expression-derived dependence cannot be excluded for analyses involving them, including the *NLRP1*–*NLRP3* comparison. In these paired measurements, *NLRP1* therefore provides a concrete counterexample to reading layer enrichment as cell-class ranking: its epidermal direction, recurrent in all 10 replication sections, coexisted with higher endothelial than keratinocyte class-pseudobulk in all 10 donors. Whether the restricted proxy recovered that direction depended on the endothelial share assumed for the dermis: at most 1 of 10 donors under either fixed weighting, but 8, 7 and 7 of 10 at shares of 0, 0.05 and 0.10 in the post hoc sweep (Supplementary Table S9). Neither observation establishes keratinocyte-intrinsic regulation or cellular origin.

The external cohort strengthened the architecture at the direct-gene level while retaining uncertainty. *NLRP1* and *PYCARD* passed the four-gene BH family, whereas the *CASP4* and *GSDMD* q values were non-significant and their direction counts remain the appropriate context. The earlier post hoc triad was epidermal in 10 of 10 sections, but *CASP4* component agreement was 7 of 10. The GSE312129 pseudobulk and reconstruction used paired modalities from the same donors, yet the modalities have distinct capture properties. Replication labels were generated by a separate expression-derived score route; verification of the discovery labels does not test or validate the replication labels. Detection of the layer split in HC discovery skin further indicates that SSc need not create the ordering de novo, although disease effects on magnitude were not formally tested.

Regression non-identifiability explains why forward reconstruction was needed. Epidermal location and keratinocyte abundance were tightly coupled, producing maximum VIF 15.5 and making the simultaneous route non-identifying. The attenuation-prone two-stage q values of 1.0 should not be read as evidence that layer polarity is absent. Reconstruction instead asked whether class-pseudobulk ordering combined with a fixed restricted proxy reproduced the observed direction. For *NLRP1*, the observed direction was more epidermal than the M1 proxy direction in 9 of 10 donors; this count depends on the assumed dermal weighting (Supplementary Table S9). This is a scale-invariant directional category, not a quantitative residual, percentage explained or cross-platform magnitude comparison.

The dermal *GSDMD* result retains a vascular and inflammatory context without assigning cellular origin [25]. In discovery sections, *GSDMD* showed the strongest positive association among tested dermal contexts with estimated endothelial abundance, and its GSE312129 class-pseudobulk ordering was higher in fibroblast and endothelial_strict than keratinocyte in all 10 donors. The modest interferon-gamma–*GSDMD* association remained positive after adjustment for Cell2location-estimated endothelial relative abundance, but this is secondary and hypothesis-generating. Endothelial adjustment applies only to the dermal interferon-gamma–*GSDMD* association and cannot be generalized to layer polarity or reconstruction. *PYCARD* and *GSDMD* agreement with the restricted proxy provides descriptive comparison, not an inferential control or proof of cellular origin.

Several limitations define the claim boundary. Visium FFPE and snMultiome are mRNA-level, platform-distinct measurements; nucleus recovery, dissociation and gene-specific capture can affect class-pseudobulk ordering. The proxy contains only keratinocyte, fibroblast and endothelial_strict and does not measure Visium composition; using the same-donor nucleus ratio (M1) or a fixed share (M2) as the fibroblast:endothelial contribution to the unmeasured Visium dermis is an assumption, and *NLRP1* concordance depended on it (Supplementary Table S9); omitted immune, perivascular, appendage and other populations can generate discordance. Layer labels and abundance estimates are expression-derived, and the snMultiome object was also the discovery cell2location reference, so this is not independent biological validation. Compartment proportions were not quantified directly. Protein localization, GSDMD cleavage, pore formation, executed pyroptosis and cellular origin were not measured, and causality is not established. The data cannot distinguish a coordinated distributed program from independent compartment-restricted programs that yield the same architecture. Broader convergence across diseases remains untested.

These boundaries define a validation program rather than diminish the finding. Target-independent layer annotation, expanded composition models and spatial single-cell assays can test whether class-level expression reproduces the *NLRP1* layer direction once dermal cell-class contributions are measured rather than assumed and omitted populations and platform capture are addressed; protein-resolved studies can then examine NLRP1, ASC and CASP4 in epidermal compartments and GSDMD localization and cleavage in dermal compartments alongside vascular markers. The present data establish neither cellular origin nor pathway execution. They show instead that anatomical organization is not a lower-resolution synonym for cell identity.

For SSc, this reframes pyroptosis-related transcription from a unitary pathway score to a distributed tissue architecture spanning epidermal and dermal contexts, with the dermal *GSDMD* pole associated with vascular and interferon-related tissue context. More generally, spatial pathway interpretation should proceed in two ordered steps: resolve gene-level anatomical polarity before aggregation, then test cellular attribution against matched cellular data rather than assuming it.

## Data Availability

Data are available in public, open access repositories. Discovery spatial transcriptomics data are available at Zenodo (doi: 10.5281/zenodo.14577696) [18], and replication data are available in GEO (GSE312129) [20]. Values for all graph data points are in the Supporting Data Values file.

## Code availability

Code, selected intermediate values and environment snapshots for direct replication, composition diagnostics and restricted reconstruction are available at Zenodo (version 1.1.0; doi: 10.5281/zenodo.22215741). Additional plotting code, frozen display inputs and figure-to-code provenance for Figure 5 and Supplementary Figures S8D, S10A and S14 are provided in Supplementary Software 1. Code, frozen input values and provenance for the separate post hoc endothelial-relative-abundance sensitivity analysis are also provided in Supplementary Software 1. Code and values for the post hoc dermal-weight sensitivity analysis (Supplementary Table S9) are also provided in Supplementary Software 1.

## Supporting information

Supporting Data Values

Supplementary Software 1

## Acknowledgements

We thank the staff at the Department of Internal Medicine, Kyushu University Beppu Hospital, for their support.

## Author contributions

D.O. conceived the study, designed the analyses, curated the public datasets, performed the computational analyses, generated the figures, interpreted the data, and drafted the manuscript. G.D. contributed to study conceptualization, interpretation of the spatial transcriptomic findings, and critical revision of the manuscript. S.F., N.N., K.O., A.K., M.A., Y.K., K.A., and H.N. contributed to clinical, immunologic, and systemic sclerosis–related interpretation of the findings and critically revised the manuscript for important intellectual content. H.M. supervised the study, contributed to study conceptualization and interpretation, and critically revised the manuscript. All authors approved the final version of the manuscript and agree to be accountable for all aspects of the work. Generative AI tools were used for the limited assistive purposes described in Methods; the authors reviewed and verified all outputs and retain full responsibility.

