## Supplementary Information for "Layer-to-cell-class correspondence is gene-specific for pyroptosis-related transcription in human skin"

Supplementary Methods and Results are followed by Tables S1–S9, Figures S1–S14 and Supplementary References; full-precision source values and provenance are provided in the Supporting Data workbook.

#### Supplementary Methods

#### 1. Spatial data acquisition and preprocessing

The discovery dataset comprised Visium formalin-fixed paraffin-embedded (FFPE) skin sections generated from 4 healthy control (HC) donors and 10 systemic sclerosis (SSc) donors, as reported by Li et al. [1]. Loading was attempted for 14 repository folders. Thirteen loaded successfully; the SSc-HL33 repository folder could not be loaded because spatial/tissue_positions_list.csv was unavailable. The final analytic discovery cohort therefore comprised 13 loaded sections (4 HC and 9 SSc): HC01, HC02, HC03, HC05, SSc-HL01, SSc-HL05, SSc-HL06, SSc-HL11, SSc-HL13, SSc-HL25, SSc-HL35, SSc4994, and SSc5380.

Brightfield histology images paired to the Visium FFPE workflow were displayed for spatial orientation but were not inputs to layer-label generation.

The executed concatenation bookkeeping shifted the last three display identifiers after the failed middle load. Supplementary Table S2 gives the authenticated repository-to-report crosswalk.

Correcting report identifiers changes neither the analytic matrices nor any numerical result.

Exact retained-barcode identity, array-coordinate agreement, reconstructed log1p(10,000) matrix equivalence, and the Figure 1D image-to-section authentication manifest were used to bind report labels to raw repository folders.

For replication, we reanalyzed the publicly available Whitfield/Jarnagin Visium cohort (GSE312129), which contains 10 SSc skin sections from an independent study of early untreated disease [2]. We used publicly available processed spatial count matrices and associated metadata for both cohorts and did not rerun FASTQ-to-count processing. The replication source study reports initial alignment and quality control with Space Ranger v2.0.0 [2]. Subsequent layer assignment, endpoint calculation, and section-level aggregation followed the same inferential framework in the two cohorts, with the exception that the replication cohort was SSc-only by design.

#### 2. Quality control filtering

Quality control was applied at spot and section levels before any pyroptosis-focused analysis. At the spot level, locations with fewer than 200 detected genes or mitochondrial-gene fractions greater than 20% were excluded. Layer-bias and endpoint analyses were then performed only on QC-passing in-tissue spots.

No section-level low-library-complexity exclusion was implemented in the present discovery analysis. All 13 successfully loaded sections, including SSc5380, were retained after the implemented spot-and feature-level preprocessing. Within retained sections, gene-detectability checks were performed before analysis-family summaries were computed so that missing feature-space genes were carried forward as explicit missingness rather than silently converted to zero values.

#### 3. Layer annotation and compartment assignment

Discovery layer labels were generated computationally from eight marker-score panels and a hybrid expression-spatial graph. Standardized spatial coordinates, weighted by two, were combined with the first 10 PCs from 2,000 highly variable genes to construct a 10-neighbor graph, followed by Leiden clustering at resolution 0.8. Marker-derived pseudo-labels were refined by multinomial logistic regression using 30 PCs, two spatial-smoothing iterations, and one-ring boundary labeling. This produced layer_v2 and layer_v2_pure. For Figure 1C, sectionwise Cohen's d was calculated from layer_v2_pure, retaining only Epidermis and Dermis spots and excluding Boundary, Adipose, Other, and missing assignments. The six-gene direct-expression screen in Figure 2A and later module-score analyses used the broader layer_v2 assignment; Figure 2A used sectionwise arithmetic means for Epidermis and Dermis before forming Δ(Epidermis − Dermis). The spot-count denominators of these estimands are not interchangeable. Histology images were displayed for orientation but were not label inputs.

Replication spots were assigned by the maximum of expression-derived marker scores for epidermis (*KRT14*, *KRT5*, *KRT1*, and *KRT10*), dermis (*COL1A1*, *COL1A2*, *COL3A1*, *VIM*, and *DCN*), and endothelium (*PECAM1*, *CDH5*, *VWF*, and *ERG*). The label other was assigned when the maximum score was nonpositive or its margin over the second-highest score was below 0.05.

This expression-derived three-class route did not use the Discovery HVG, PCA, Leiden, pseudo-label, logistic-regression, or spatial-smoothing pipeline and is not an independent histologic validation.

For the signed-distance analysis, a computational boundary label was retained for spots nearest the estimated layer interface. The same computationally generated layer and boundary labels were reused across related analyses within each section so that representative maps, all-section summaries, and boundary-centered descriptive curves referenced the same stored compartment calls. Boundary thumbnails and section-level annotation panels are provided for visual inspection of the resulting epidermis, dermis, boundary, and other assignments.

In a verification exercise conducted before submission, the layer-assignment pipeline was re-executed from the authenticated source under a captured runtime. Two independent clean re-executions produced identical labels, and all 11 marker and composite score arrays reproduced the stored values exactly (maximum absolute difference 0), as did the 2,000-gene highly variable set. Small numerical differences at the principal-component stage propagated to the neighbourhood graph and the Leiden partition (26 stored versus 25 regenerated clusters), so the regenerated labels were not identical to the stored labels: 532 of 12,210 spots differed for layer_v2 and 987 of 12,210 for layer_v2_pure. Restricted to spots labelled Epidermis or Dermis in the stored assignment, layer_v2 concordance was 98.4%, and direct Epidermis–Dermis exchanges in layer_v2_pure affected 7 spots. The remaining differences were concentrated in auxiliary tissue classes and in the width of the boundary ring. The reported analyses use the stored labels. The unique cause of the divergence was not established.

The central outcome genes did not overlap the marker panels used to generate Discovery pseudo-labels. *PYCARD*, *NLRP1*, *CASP4*, and *GSDMD* were not among the 2,000 HVGs used for PCA, whereas *NLRP3* and *IL1B* were HVGs. Direct marker-list circularity is absent, but expression-derived dependence for analyses involving those HVGs cannot be excluded. This verification applies only to the Discovery pipeline and does not test or validate the distinct Replication marker-score labels.

#### 4. Inference unit and pseudoreplication avoidance

The tissue section was the inferential unit throughout. Individual Visium spots within a section were not treated as statistically independent because they share tissue context, section-specific processing, and spatial autocorrelation. Accordingly, spot-level maps and within-section distributions were interpreted descriptively, whereas formal cross-section summaries were built from one statistic per section per endpoint.

Within-section summaries included compartment medians or means, signed layer deltas, effect sizes, and section-level correlation coefficients. Cross-section aggregation was then performed on those section-level values rather than on the pooled spot matrix. Any pooled Fisher z result was reported only as a secondary sensitivity summary of section-level coefficients and did not alter the section as the inferential unit.

#### 5. Signed effect-size conventions

Unless otherwise noted, the default sign convention was Dermis minus Epidermis, so that positive values indicated dermal enrichment and negative values indicated epidermal enrichment.

This convention was used for section-level gene summaries, cohort-wide dumbbell plots, and dermal-bias endpoints.

Two exceptions were retained for consistency with the final figure displays. First, the direction-concordant-triad composite comparison used Delta(Epidermis − Dermis) so that larger positive values represented stronger epidermal separation. Second, replication-endpoint scores preserved the native direction of each endpoint; the manuscript therefore reports endpoint direction explicitly rather than relying only on sign. Figure 2A reports signed epidermal consistency, whereas Supplementary Fig. S5 reports majority-direction concordance and names that direction; these are related but distinct summaries and are not numerically interchangeable.

#### 6. Gene-set definitions used in the study

Three distinct gene-set families were used.

Exploratory discovery screen. Eight genes were considered: *PYCARD*, *NLRP1*, *CASP4*, *NLRP3*, *AIM2*, *MEFV*, *CARD8*, and *CASP1*. *CARD8* and *CASP1* had probe rows in the relevant probe-set file but were excluded during probe filtering and were absent from the filtered discovery FFPE feature space (var_names); they were therefore carried as explicit missing data rather than treated as zero expression. The evaluable exploratory screen consisted of *PYCARD*, *NLRP1*, *CASP4*, *NLRP3*, *AIM2*, and *MEFV*.

Gasdermin-family context set. Direct normalized *GSDMD* expression was used for gene-level maps, context correlations, and the Whitfield replication endpoint. The discovery cross-cohort endpoint panels instead used the precomputed Scanpy *GSDMD*-only module score pyro_gasdermin_score, summarized by compartment mean. Context panels used *GSDMD*, *GSDMA*, *GSDMC*, *GSDMB*, and *GSDME* where evaluable; *PJVK* was unavailable in the gasdermin-family output. These direct-expression and module-score quantities are distinct and are identified separately throughout this supplement.

Replication endpoint support set. The prespecified epidermal inflammasome-related endpoint carried into the replication framework used a 5-gene support set comprising *PYCARD*, *NLRP3*, *CASP1*, *IL1B*, and *IL18*. This support set was used to contextualize the epidermal endpoint score in the anchor-gene heatmap and related replication summaries. It was prespecified as a separate endpoint-level construct and was not intended as a rediscovery of the discovery *NLRP1–PYCARD–CASP4* triad.

Implemented score objects. Supplementary Table S1 distinguishes the six precomputed score columns from the exploratory screen and the direction-concordant triad. The P3 device score was not a flat 12-gene score: it was the equal-weight mean of three module scores containing 7, 2, and 3 evaluable genes, respectively, so the module scores, not individual genes, had equal weight.

#### 7. Exploratory discovery direction-concordance screen

The exploratory classification was based on sectionwise directional consistency in the 13-section discovery cohort. The formal screen specification was written after the direction-concordant triad had been observed; the screen, its exact sign tests, and its BH-adjusted q values are therefore reported as exploratory and post-selection rather than prespecified. For Figure 2A, bars show the number of evaluable sections with Δ(Epidermis − Dermis) > 0; annotations give n/N and BH-adjusted q values. The corresponding fraction is the number with epidermal direction divided by the number evaluable. Supplementary Fig. S5 instead reports majority-direction concordance as max(n_epidermal, n_dermal) / n and explicitly names the majority direction. *NLRP1*, *PYCARD*, and *CASP4* had epidermal direction in 13 of 13 sections and are described as a direction-concordant epidermal triad; no formal biological class is claimed.

A gene was considered evaluable only if present in the filtered QC-passing feature space.

*CARD8* and *CASP1* were excluded during probe filtering and absent from the filtered discovery feature space; they were rendered as explicit missingness and excluded from concordance denominators. This wording distinguishes filtered-feature absence from absence of probe rows in the original probe-set file.

#### 8. Composite score construction

Two upstream composite summaries were compared to describe whether a direction-concordant triad preserved the epidermal signal more strongly than a broader mixed-direction panel.

Figure 2C uses the frozen section-level contrasts in fig2_signed_partial_metrics.xlsx, sheet S3_SIGNED_DILUTION. The source table contains one paired legacy-field delta_signed_t1 and delta_all value per discovery section; delta_signed_t1 denotes the direction-concordant *NLRP1–PYCARD–CASP4* triad in this manuscript. This source-specific estimand is distinct from the separate direct gene-mean triad used in Supplementary Fig. S7 and the targeted replication analysis. The broader set comprised *PYCARD*, *NLRP1*, *CASP4*, *NLRP3*, *AIM2*, and *MEFV*; the triad comprised *PYCARD*, *NLRP1*, and *CASP4*. Each spot-level score was the mean expression of its target genes minus the mean expression of a sampled background-gene set, using the stored log1p-normalized matrix X without further normalization at this step. The implemented helper used all retained features as the gene pool, expression-ranked bins with n_bins=25, up to ctrl_size=50 background genes from each represented bin, and random_state=0. Its ctrl_as_ref=True setting did not remove target genes from the background candidates. Within each section, arithmetic mean scores over layer_v2 Epidermis and Dermis spots were subtracted as Epidermis − Dermis; layer_v2_pure was not used for this comparison. These background-adjusted scores are not direct target-gene means.

The dilution statistic was calculated from those frozen section-level contrasts as:

dilution = 1 − mean(|delta_all|) / mean(|delta_direction_concordant_triad|)
Applied to the 13 discovery sections, this calculation yielded 60.8%. Because the contrast was selected from directional consistency in the same discovery data, the value is descriptive and post-selection; no formal P value was assigned.

#### 9. Replication endpoint design

Replication was framed as endpoint-level directional support using related but nonidentical cohort-specific endpoints rather than as a second feature-selection exercise. The discovery and Whitfield constructs are defined separately below and were not treated as the same score.

For the discovery comparison, the dermal endpoint was the precomputed Scanpy *GSDMD*-only module score pyro_gasdermin_score and section-level values were compartment means. For Whitfield, the prespecified dermal endpoint was direct log1p-normalized *GSDMD* expression and section-level values were also compartment means. These related endpoints share direction and aggregation but are not numerically identical constructs.

For the discovery comparison, the epidermal endpoint was the precomputed Scanpy *IL1B–IL18* cytokine-maturation module score pyro_B_maturation_score and section-level values were compartment medians. For Whitfield, the prespecified endpoint used *PYCARD*, *NLRP3*, *CASP1*, *IL1B*, and *IL18* and section-level values were compartment medians. Both differed from the direct gene-mean *NLRP1–PYCARD–CASP4* triad. Because Whitfield contained SSc sections only, the discovery endpoint comparison was restricted to the 9 SSc discovery sections.

Anchor-gene heatmaps were scaled within cohort rather than across cohorts. This preserved directional patterning while avoiding the implication that absolute score ranges were directly comparable across studies processed in different source pipelines.

Separately, after the locked endpoint analysis had been defined, a targeted post hoc analysis evaluated the discovery-fixed *NLRP1*, *PYCARD*, and *CASP4* triad directly in the same 10 replication sections. For each gene and section, the epidermal and dermal summaries were the means of log1p-normalized expression across spots in the corresponding compartment, and the signed gene contrast was Epidermis − Dermis. The signed triad contrast was the sectionwise mean of the three fixed gene contrasts. A section was evaluable only when all three genes were present and both layers were available; partial-gene composites and zero substitution for missing values were prohibited.

All 10 sections met these criteria. The section was the inferential unit. Direction counts and the distribution of section-level contrasts were the primary descriptive outputs. A single two-sided one-sample Wilcoxon signed-rank test of the 10 section-level signed-triad contrasts against zero was reported as exploratory and secondary; no spot-level pooled inference was performed. This analysis was kept analytically separate from the prespecified five-gene endpoint because the gene sets, score construction, and compartment summaries differed.

#### 10. Cell2location deconvolution parameters

Spot-level cell-type abundance was estimated with cell2location v0.1.5 [3]. The reference atlas was taken from the Whitfield/Jarnagin single-nucleus multiome study and contained 65,066 nuclei from 4 control and 10 SSc skin samples [2]. Initial reference labels were assigned with CellTypist 1.7.1 using Immune_All_Low.pkl and majority voting [4], followed by the implemented manual consolidation map, marker fallback, and an endothelial_strict override requiring both *PECAM1* and *CDH5*. The CellTypist model source was recorded; model-file hash unavailable. The reference atlas and replication Visium sections came from paired biopsies from the same 10 participants and therefore provide linked orthogonal modalities rather than statistically independent participant sets.

Reference signatures were estimated once from the processed source atlas, and the spatial mapping step was then run per discovery section rather than jointly across all sections. The standard negative-binomial reference-signature and spatial-mapping workflow from cell2location v0.1.5 was used, with the implemented label consolidation, marker fallback, and endothelial_strict override described above. Model training was run on a T4 GPU in Google Colab Pro.

The deconvolution output was interpreted as an estimated abundance matrix rather than as a direct cell-count matrix or evidence of cellular origin. This interpretation applies to the cell2location-derived analyses; the separate precomputed endothelial_score used in ancillary analyses is addressed in Section 11. All compartment-specific context analyses used the relevant computationally generated layer labels.

Within each section, Spearman correlations were calculated within the anatomically restricted compartment. The tissue section remained the inferential unit. For Supplementary Fig. S13C, fixed-effect Fisher-z pooling weighted by the number of contributing spots minus 3 was used only as a secondary display summary, and no pairwise test of superiority among contexts was performed. For Supplementary Fig. S13D, each plotted point and stem represents one section-level Spearman coefficient. The two-sided one-sample t-test evaluates the mean of the 13 section-level coefficients against zero, and its P value was adjusted by the Benjamini-Hochberg procedure within the four-test deconvolution family; this test is distinct from the secondary Fisher-z pooled display summaries. In Supplementary Fig. S13C, horizontal segments span the 5th to 95th percentiles of the section-level coefficients for each context; they are descriptive between-section intervals, not confidence intervals for the pooled estimate.

The four tests in the deconvolution family were: (1) within-dermis section-level Spearman correlation of direct normalized *GSDMD* expression with estimated endothelial abundance; (2) within-dermis section-level Spearman correlation of direct normalized *GSDMD* expression with estimated fibroblast abundance; (3) within-epidermis section-level Spearman correlation of the precomputed pyro_inflammasome_device_score with estimated keratinocyte abundance; and (4) the direction across sections of the high-minus-low difference in estimated endothelial relative abundance after splitting QC-passing dermal spots at the within-section median of the precomputed fibrosis_score. The first three tests used two-sided one-sample t-tests of the 13 section-level Spearman coefficients against zero, and the fourth used a two-sided exact sign test across sections. P4 was negative in 12 of 13 sections (high minus low), with exact P = 0.00341796875 and BH q = 0.004557291666666667 within the fixed four-test family. Both precomputed score columns were used as generated in the source analysis objects and were not recalculated for manuscript reporting.

#### 11. Partial correlation and sensitivity analyses

The implemented IFNγ-response score was computed at spot level using 13 genes (*IRF1*, *IRF2*, *STAT1*, *GBP1*, *GBP2*, *GBP4*, *GBP5*, *IFNGR1*, *IFNGR2*, *CIITA*, *CXCL9*, *CXCL10*, and *CXCL11*). The frozen analysis plan listed 12 genes and omitted *IRF2*; the plan remains unchanged, and this report uses the implemented 13-gene score without relabelling that exact composition as prespecified. The spot-level score was calculated from log1p-normalized expression in the discovery-cohort analysis AnnData using sc.tl.score_genes() with ctrl_size = 50 and random_state = 42. IFNG ligand itself was intentionally excluded. Dermal-only QC-passing spots were subset within section before rank transformation, with no cross-section rescaling beyond that rank transform.

The archived ancillary analysis used the precomputed endothelial_score column as a covariate in partial Spearman correlations within the dermal compartment of each discovery section. Only QC-passing dermal spots from that section contributed to the calculation. Executed records show reuse of the existing score, not execution of the fallback branch that could generate a score from *PECAM1*, *VWF*, and *CDH5*. The original construction of the precomputed endothelial_score could not be verified; analyses using this covariate are therefore not interpreted as adjustment for endothelial cell abundance. No marker set or equivalence to the separate cell2location c2l_rel_* columns is inferred from the column name. The same provenance limitation applies to the endothelial_score map in Supplementary Fig. S13A.

Operationally, the IFNγ score, *GSDMD* expression, and precomputed endothelial_score were rank-transformed within section. The ranks of the IFNγ score and *GSDMD* were then residualized on the ranked endothelial_score vector, and the section-level partial Spearman rho was computed as the Pearson correlation between the two residual vectors. The corresponding zero-order Spearman rho was computed on the unadjusted ranked variables. The attenuation metric was defined as:

Delta rho = rho_zero-order - rho_partial
The frozen partial-correlation analysis implemented the planned P1/P2 Fisher-z family. At manuscript reporting, P1 section-level coefficients were summarized primarily by their distribution, and an equal-weight two-sided one-sample Wilcoxon signed-rank test of the 13 section-level P1 values was designated the primary formal summary. The planned Fisher-z pooling was retained as a secondary summary [5], with weights proportional to $n - 3$. This reporting-stage change was conservative and did not alter any section-level estimate. The analysis plan's secondary macrophage-abundance covariate was implemented as the precomputed 12-gene mk_immune score; it is therefore described as an additional immune-score sensitivity covariate rather than as the prespecified macrophage covariate. Sensitivity analyses also examined leave-one-section-out stability and alternative adjustment variables, including fibroblast abundance.

In a separate post hoc sensitivity analysis, the saved IFNG_score and GSDMD_norm values were retained for all 9,224 QC-passing dermal spots from the same 13 discovery sections (4 HC, 9 SSc). Spots were matched by identifier between the archived partial-correlation input and the Cell2location v3 output, retaining the stored layer_v2 Dermis labels without imputation or additional exclusions. The new covariate was the saved c2l_rel_q05cell_abundance_w_sf_endothelial_strict column: the endothelial_strict q05 abundance divided by the sum of q05 abundances across all 13 classes, including unknown and stromal_reclassified. This is an estimated relative abundance, not a measured cell fraction or the posterior fifth percentile of a fraction. The saved column was used without recomputing the ratio at a different numerical precision.

Within each section, IFNG_score, GSDMD_norm and the new covariate were converted to average ranks. Ranked IFNG_score and ranked GSDMD_norm were separately residualized on an intercept and the ranked covariate, and the Pearson correlation of the residuals gave the partial Spearman coefficient. All 13 coefficients contributed with equal section weight. The sole new test was a two-sided one-sample Wilcoxon signed-rank test using exhaustive sign permutation, zero_method=wilcox and no continuity correction; all 13 coefficients were nonzero and had no tied absolute ranks. The 95% confidence interval for the median was a percentile bootstrap over sections with 2,000 resamples and seed 42, not a spot-level bootstrap or Fisher-z pooling interval.

This analysis used the same discovery sections rather than independent replication data. It did not recalculate the IFNγ-response score, normalize expression again, refit Cell2location, or select an alternative covariate or subset. The covariate is expression-derived, and posterior uncertainty was not propagated. The results therefore concern a post hoc association conditional on estimated endothelial relative abundance, not independence from true cell abundance, all cell composition or causal effects. The historical endothelial_score analysis and its unresolved source provenance are retained separately. Supplementary Table S8 lists all section-level results, and the Supporting Data workbook links legacy source labels to the established report identifiers.

#### 12. DEJ signed-distance analysis

The signed-distance analysis was descriptive and was designed to place the layer-biased signals into a common boundary-centered coordinate system. Computed boundary-anchor spots nearest the estimated layer interface were used as distance anchors. For every spot in a section, the distance to the nearest boundary anchor was calculated in image-coordinate space and scaled by the section-specific Visium pitch so that the output could be expressed in approximate 100-μm bins.

Signed distance was negative on the epidermal side, positive on the dermal side, and zero at the computed boundary-anchor set. Because this approach depends on discrete computational labels and pitch scaling rather than on a histology-segmented continuous contour, the resulting curves were interpreted as descriptive spatial profiles rather than as exact anatomic reconstructions. The same signed-distance framework was used for the direction-concordant triad and control-pathway comparisons so that modality contrasts were made in a matched coordinate system. For Figure 2D, faint points are individual spots within the displayed distance window; when necessary, only the point display is subsampled to at most 3,800 spots per cohort per gene. Lines use the stored mean_expr_smooth3 values: arithmetic means of pooled spots within each cohort and 100-μm bin, followed by a centered three-bin rolling mean with at least one available bin. Sections are not given equal weights in these descriptive curves. Smoothing precedes the display filter, which retains bin centers from −800 to 1,400 μm with at least 10 spots in the corresponding unsmoothed bin. *PYCARD*, *NLRP1*, and *CASP4* are shown separately; the curves are not a triad-composite score.

#### 13. Negative-control pathways and compact comparator families

Negative-control analyses were designed as descriptive compact-set comparisons of the upstream-versus-effector split across pyroptosis-related, apoptosis, and necroptosis modules. The same section-level layer-bias framework was applied to each module, and a compact gasdermin-family comparator set was analyzed alongside *GSDMD*. Because the component sets differed in size and composition, these comparisons were used for pathway context and were not interpreted as formal proof of pathway specificity.

For the gasdermin-family comparison, the contextual comparator genes were *GSDMA*, *GSDMC*, *GSDMB*, and *GSDME* where evaluable, with *GSDMD* retained as the direct dermal endpoint; *PJVK* was not present/evaluable in the gasdermin-family analysis output. Figure S7 used compact upstream/effector component sets matched to the submitted figure source. The pyroptosis-related set used an upstream component (*NLRP1*, *PYCARD*, *CASP4*) and an effector component (*GSDMD*). The apoptosis control used upstream components *BAX*, *BAK1*, *CASP8*, and *CASP9* and effector components *CASP3* and *CASP7*. The necroptosis control used upstream components *RIPK1* and *RIPK3* and effector component *MLKL*. The same section-level layer-bias framework was applied across these component sets.

#### 14. Multiple testing correction

Multiple testing was handled within analysis-specific families rather than across the manuscript as one omnibus pool. Prespecification status is stated separately for each family. The Benjamini-Hochberg false discovery rate procedure [6] used $q < 0.05$ where formal or exploratory family-wise correction was applied.

The exploratory six-gene discovery screen used BH correction across its six evaluable genes but remained post-selection. The prespecified deconvolution family comprised four tests (P1–P4), all of which are now reported. The planned partial-correlation family comprised Fisher-z P1 and P2 with BH correction across $m = 2$; those pooled results remain secondary after the reporting-stage choice of the equal-weight section-level P1 Wilcoxon summary. Compact apoptosis and necroptosis comparisons were descriptive, introduced no additional formal test, and were not a BH family. The separate post hoc Cell2location-relative-abundance sensitivity analysis introduced one section-level Wilcoxon test; its P value is unadjusted, is not a BH q value, and does not alter the previously reported test families.

All formal tests were two-sided, with a nominal alpha level of 0.05. Wilcoxon signed-rank tests were used for small section-level samples because they do not require a normality assumption.

Exact P values, effect sizes, confidence intervals, and BH-adjusted q values are reported where applicable in the main Results and supplementary displays. No a priori sample-size calculation was performed because the analyses were based on fixed available sections from previously generated cohorts.

#### 15. Software and runtime environment

The main local analysis used Python 3.12.2, scanpy 1.11.5, and numpy 2.4.0. Other package versions and the complete environment specification are recorded in the public versioned code deposit and its environment manifests (version 1.1.0; doi: 10.5281/zenodo.22215741). The separate post hoc sensitivity analysis used Python 3.12.2, NumPy 1.26.4, pandas 2.3.3, SciPy 1.16.3 and h5py 3.15.1. Its code, frozen input values and environment record are provided in Supplementary Software 1, separately from the public Zenodo v1.1.0 deposit.

The cell2location/CellTypist stage used Python 3.12.12, scanpy 1.12, numpy 2.0.2, CellTypist 1.7.1, cell2location 0.1.5, scvi-tools 1.4.2, PyTorch 2.10.0, and pyro-ppl 1.9.1 on a T4 GPU in Google Colab Pro. This runtime was distinct from the main local analysis environment.

Use of generative AI is described in the main Methods.

#### Direct four-gene replication analysis

Ten GSE312129 Visium sections were evaluated with the stored replication labels. The primary mask retained only spots labelled epidermis or dermis; 270 endothelium-labelled and 909 other-labelled spots were excluded. This definition is not symmetric with the discovery layer_v2_pure field, which has no corresponding endothelial class. For *NLRP1*, *PYCARD* and *CASP4*, the expected-direction contrast was the arithmetic epidermis mean minus dermis mean; for *GSDMD*, the physical contrast was reversed for the expected dermal direction. The section was the inferential unit. Exact two-sided sign tests were adjusted by BH across the four fixed genes (m=4). Bootstrap intervals for descriptive medians used 10,000 section-level resamples at seed 42. A dermis-plus-endothelium mask was retained only as a descriptive sensitivity without additional P values.

#### Same-donor cell-class pseudobulk

Raw counts from the same-donor snMultiome object were aggregated separately for keratinocyte, fibroblast and endothelial_strict nuclei. The library denominator comprised 18,703 retained features. CPM was calculated as $10^6 \times$ summed counts / retained-feature library size and transformed as log2(CPM+1). The primary SSc analysis required at least 20 nuclei in each of the three classes; a floor of 50 was descriptive sensitivity. HC donors were descriptive only. With K, F and E denoting the donor-specific class log2(CPM+1) values, the expected-direction contrast was $K - (F + E)/2$ for *NLRP1*, *PYCARD* and *CASP4*, and $(F + E)/2 - K$ for *GSDMD* (Supplementary Table S4). Exact two-sided sign tests across donors excluded zero contrasts and used BH correction across the four genes (m=4). These contrasts average F and E after log transformation, unlike the linear-CPM reconstruction. The snMultiome object was the same reference object previously used for discovery cell2location and therefore does not constitute independent biological validation of that model.

#### Composition diagnostics and restricted mixture reconstruction

Within each discovery section, ordinary least squares modeled normalized gene expression as a function of layer (epidermis versus dermis) and the posterior-median (q50) cell2location abundances of keratinocyte, fibroblast and endothelial_strict classes. Each layer required at least 20 spots. For collinearity assessment, the layer indicator and abundance predictors were standardized. All predictor VIFs had to be finite and at most 10, and the condition number had to be finite and at most 30. The simultaneous route required all 13 sections to pass these criteria. A two-stage route first removed the linear abundance component and then summarized section-level residual layer contrasts; this estimator is structurally attenuation-prone. The active four-gene route used exact two-sided sign tests of section-level effects with BH correction (m=4); the non-estimable simultaneous route had no formal P values. Neither route was altered after inspection. For the forward reconstruction, observed Visium direction was $O_{dg}$ = epidermis_mean_log1p − dermis_mean_log1p. Cell-class CPMs were mixed on the linear CPM scale before conversion to log2(CPM+1). M1 treated epidermis as 100% keratinocyte and used the donor-specific fibroblast:endothelial nucleus ratio for dermis. M2 used equal fibroblast:endothelial weights. The primary readout was exact sign concordance. Raw cross-platform residuals, percent explained and R-squared were prohibited. Leave-one-donor-out values were descriptive influence summaries without refitting or formal inference.

#### Provenance

Execution-receipt identifiers and source provenance are provided in the Supporting Data workbook (AnalysisStatus and IFNG_C2L_Summary sheets).

### Supplementary Methods (continued)

#### Composition diagnostics and restricted mixture reconstruction (continued)

Post hoc dermal-weight sensitivity analysis. For each of the 10 paired SSc donors and four fixed genes, the dermal proxy CPM was calculated as (1 - w) x fibroblast CPM + w x endothelial_strict CPM, with keratinocyte CPM as the epidermal proxy. Class CPMs were mixed before conversion to log2(CPM+1). The predicted sign of the keratinocyte-minus-dermal-proxy contrast was compared with the observed Visium epidermis-minus-dermis mean-log1p sign; exact sign concordance was counted across donors.

The fixed endothelial share w was swept across 0, 0.05, 0.10, 0.15, 0.20, 0.25, 0.30, 0.40, 0.50, 0.75 and 1.00. As internal controls, the original donor-specific nucleus-ratio model M1 (w = nE/(nF+nE)) reproduced concordance counts of 1/9/4/8, and the equal-weight model M2 (w = 0.5) reproduced 0/9/3/8, in NLRP1/PYCARD/CASP4/GSDMD order. This post hoc analysis was descriptive and used no P values (Supplementary Table S9).

#### Supplementary Results

The previously reported related-endpoint replication is retained for continuity: the direct *GSDMD* endpoint was dermally directed in 8 of 10 SSc sections, the separate five-gene endpoint was epidermally directed in 10 of 10, and the post hoc discovery triad was epidermally directed in 10 of 10. Component agreement for the triad was *NLRP1* 10 of 10, *PYCARD* 9 of 10 and *CASP4* 7 of 10. These constructs are related but not identical to the direct four-gene estimands in Table S3.

The direct four-gene analysis showed epidermal *NLRP1* in 10 of 10 sections (q=0.0078125), epidermal *PYCARD* in 9 of 10 (q=0.04296875), epidermal *CASP4* in 7 of 10 (q=0.34375), and dermal *GSDMD* in 8 of 10 (q=0.14583333333333334). The *CASP4* and *GSDMD* q values were non-significant; direction counts provide the principal replication context.

In SSc donor pseudobulk, *PYCARD* was higher in keratinocyte than fibroblast in 10 of 10 donors and higher than endothelial_strict in 9 of 10. *GSDMD* was higher in fibroblast and endothelial_strict than keratinocyte in all 10 donors. *NLRP1* was higher in keratinocyte than fibroblast in 8 of 10 but lower than endothelial_strict in all 10; *CASP4* was higher in keratinocyte than fibroblast in 7 of 10 but lower than endothelial_strict in 9 of 10. These are class-pseudobulk contexts, not per-cell expression or assignments of cellular origin. HC values and floor-50 results are descriptive.

The simultaneous discovery model was non-estimable because the maximum VIF was 15.545483429882879 and 5 of 13 sections failed the fixed collinearity gate. All four two-stage BH q values were 1.0; because this route is structurally attenuation-prone, this does not test the absence of layer polarity. The replication endothelial-context analysis was NOT_ESTIMABLE_ENDOTHELIAL_CONTEXT because the only available score was part of the expression-derived label generator; no replacement score was created.

The restricted proxy reproduced the observed direction for *NLRP1* in 1 of 10 donors under M1 and 0 of 10 under M2; for *PYCARD* in 9 of 10 under both models; for *CASP4* in 4 of 10 under M1 and 3 of 10 under M2; and for *GSDMD* in 8 of 10 under both models. Under M1, *NLRP1* was OBSERVED_MORE_EPIDERMAL_THAN_PROXY in 9 of 10 and *CASP4* in 6 of 10. These directional differences do not demonstrate a layer-intrinsic program, keratinocyte upregulation or cellular origin; omitted cell classes, nucleus recovery and platform-capture differences remain alternative explanations.

Cross-modal effect-size coupling was descriptive only (n=10; approximate standard error about 0.33): *NLRP1* rho = -0.0667, *PYCARD* rho = -0.1879, *CASP4* rho = -0.4303 and *GSDMD* rho = 0.3212. All values were not distinguishable from zero and are not used for inference.

For completeness, the confounded discovery dermal endothelial-context sensitivity is shown for all four genes together: positive direction occurred in 13 of 13 sections for *NLRP1*, 12 of 13 for *PYCARD*, 13 of 13 for *CASP4* and 13 of 13 for *GSDMD*. Total-cell-abundance confounding was not controlled, so these values remain descriptive and are not used to privilege *GSDMD*.

#### Supplementary Table S1. Implemented score definitions used in the reported analyses.

| Score column | Calculation | Genes used | Actual n | Compartment use | Cohort |
|---|---|---|---|---|---|
| pyro_gasdermin_score | Scanpy sc.tl.score_genes(); single-gene module | *GSDMD* | 1 | Compartment mean | Discovery |
| pyro_B_maturation_score | Scanpy sc.tl.score_genes() | *IL1B*, *IL18* | 2 | Compartment median | Discovery |
| pyro_inflammasome_device_score | Equal-weight mean of three Scanpy module scores (A1/A2/auxiliary) | A1: *NLRP3*, *NLRP1*, *NLRC4*, *AIM2*, *MEFV*, *PYCARD*, *NEK7*; A2: *CASP4*, *CASP5*; auxiliary: *P2RX7*, *TLR4*, *PSTPIP1* | 12 (7/2/3) | Spot-level; within-epidermis sectionwise Spearman correlation | Discovery |
| fibrosis_score | Scanpy sc.tl.score_genes(); 14 of 15 planned genes implemented | *COL1A1*, *COL1A2*, *COL3A1*, *COL4A1*, *COL5A1*, *FN1*, *ACTA2*, *TGFB1*, *TGFB2*, *PDGFA*, *PDGFB*, *MMP2*, *MMP9*, *TIMP1* | 14 | Within-dermis median split | Discovery |
| mk_immune | Scanpy sc.tl.score_genes() | *PTPRC*, *CD3D*, *CD3E*, *CD8A*, *CD4*, *MS4A1*, *CD19*, *CD14*, *FCGR3A*, *ITGAX*, *CD1C*, *CLEC4C* | 12 | Dermal spot-level sensitivity covariate | Discovery |
| IFNG_score | Scanpy sc.tl.score_genes(ctrl_size=50, random_state=42) | *IRF1*, *IRF2*, *STAT1*, *GBP1*, *GBP2*, *GBP4*, *GBP5*, *IFNGR1*, *IFNGR2*, *CIITA*, *CXCL9*, *CXCL10*, *CXCL11* | 13 | Dermal spot-level score; sectionwise rank-based correlation | Discovery |

The frozen 15-gene fibrosis plan included *CTGF*; the implemented source-object fibrosis_score used the 14 listed genes and omitted *CTGF*. The frozen analysis plan was not changed, and the precomputed score was not recalculated for reporting. The inflammasome-related constructs were therefore kept distinct: the eight-gene exploratory discovery screen, the three-gene *NLRP1–PYCARD–CASP4* direction-concordant triad, and the 12-evaluable-gene P3 device score formed as a 7/2/3 module-level average.

#### Supplementary Table S2. Repository-to-report section identifier crosswalk.

| raw_repo_id | legacy_R5.2_label | corrected_report_id | inclusion_status | retained_spots | reason |
|---|---|---|---|---|---|
| HC01 | HC01 | HC01 | included | 1858 | unchanged |
| HC02 | HC02 | HC02 | included | 645 | unchanged |
| HC03 | HC03 | HC03 | included | 1805 | unchanged |
| HC05 | HC05 | HC05 | included | 822 | unchanged |
| SSc-HL01 | SSc-HL01 | SSc-HL01 | included | 453 | unchanged |
| SSc-HL05 | SSc-HL05 | SSc-HL05 | included | 508 | unchanged |
| SSc-HL06 | SSc-HL06 | SSc-HL06 | included | 608 | unchanged |
| SSc-HL11 | SSc-HL11 | SSc-HL11 | included | 672 | unchanged |
| SSc-HL13 | SSc-HL13 | SSc-HL13 | included | 598 | unchanged |
| SSc-HL25 | SSc-HL25 | SSc-HL25 | included | 748 | unchanged |
| SSc-HL33 | not assigned to loaded data | not applicable | load_failed | 0 | missing spatial/tissue_positions_list. csv |
| SSc-HL35 | SSc-HL33 | SSc-HL35 | included | 877 | legacy key shifted after failed middle load |
| SSc4994 | SSc-HL35 | SSc4994 | included | 1729 | legacy key shifted after failed middle load |
| SSc5380 | SSc4994 | SSc5380 | included | 887 | legacy key shifted after failed middle load; 919 spots before filtering |

#### Supplementary Table S3. Direct four-gene Visium replication

| Gene | n | Median expected-direction contrast | + | 0 | − | Sign-test P | BH q (m=4) | 95% bootstrap median interval |
|---|---|---|---|---|---|---|---|---|
| *NLRP1* | 10 | 0.1706 | 10 | 0 | 0 | 0.0019531 | 0.0078125 | 0.1103 to 0.2271 |
| *PYCARD* | 10 | 0.1079 | 9 | 0 | 1 | 0.0214844 | 0.04296875 | 0.0373 to 0.1922 |
| *CASP4* | 10 | 0.0763 | 7 | 0 | 3 | 0.34375 | 0.34375 | -0.0373 to 0.114 |
| *GSDMD* | 10 | 0.0976 | 8 | 0 | 2 | 0.109375 | 0.14583333 | 0.0442 to 0.1638 |

| Visium sample | Gene | n epi | n derm | Epi mean log1p | Derm mean log1p | Expected-dir score | Epi detect frac | Derm detect frac | Eligible |
|---|---|---|---|---|---|---|---|---|---|
| GSM9338171_SSC1_visium | *NLRP1* | 292 | 739 | 0.47857 | 0.26186 | 0.21671 | 0.3699 | 0.1461 | True |
| GSM9338171_SSC1_visium | *PYCARD* | 292 | 739 | 0.2728 | 0.09233 | 0.18047 | 0.2877 | 0.065 | True |
| GSM9338171_SSC1_visium | *CASP4* | 292 | 739 | 0.29412 | 0.18751 | 0.1066 | 0.2979 | 0.1177 | True |
| GSM9338171_SSC1_visium | *GSDMD* | 292 | 739 | 0.186 | 0.28655 | 0.10055 | 0.1986 | 0.1583 | True |
| GSM9338172_SSC2_visium | *NLRP1* | 269 | 666 | 0.59857 | 0.22463 | 0.37394 | 0.4647 | 0.1441 | True |
| GSM9338172_SSC2_visium | *PYCARD* | 269 | 666 | 0.252 | 0.0854 | 0.1666 | 0.3048 | 0.0631 | True |
| GSM9338172_SSC2_visium | *CASP4* | 269 | 666 | 0.35648 | 0.19709 | 0.15939 | 0.3717 | 0.1411 | True |
| GSM9338172_SSC2_visium | *GSDMD* | 269 | 666 | 0.29255 | 0.21334 | -0.07922 | 0.3309 | 0.1486 | True |
| GSM9338173_SSC3_visium | *NLRP1* | 317 | 247 | 0.48219 | 0.33758 | 0.14461 | 0.3628 | 0.2996 | True |
| GSM9338173_SSC3_visium | *PYCARD* | 317 | 247 | 0.29706 | 0.15013 | 0.14693 | 0.3312 | 0.1498 | True |
| GSM9338173_SSC3_visium | *CASP4* | 317 | 247 | 0.38202 | 0.28672 | 0.0953 | 0.3659 | 0.2429 | True |
| GSM9338173_SSC3_visium | *GSDMD* | 317 | 247 | 0.27177 | 0.37286 | 0.10109 | 0.3407 | 0.3117 | True |
| GSM9338174_SSC4_visium | *NLRP1* | 272 | 344 | 0.58876 | 0.41876 | 0.16999 | 0.4044 | 0.4186 | True |
| GSM9338174_SSC4_visium | *PYCARD* | 272 | 344 | 0.17294 | 0.20859 | -0.03564 | 0.2537 | 0.3198 | True |
| GSM9338174_SSC4_visium | *CASP4* | 272 | 344 | 0.29585 | 0.34147 | -0.04562 | 0.3199 | 0.4186 | True |
| GSM9338174_SSC4_visium | *GSDMD* | 272 | 344 | 0.13449 | 0.22907 | 0.09458 | 0.2574 | 0.3285 | True |
| GSM9338175_SSC5_visium | *NLRP1* | 301 | 808 | 0.41981 | 0.34382 | 0.07599 | 0.402 | 0.2958 | True |
| GSM9338175_SSC5_visium | *PYCARD* | 301 | 808 | 0.44441 | 0.20698 | 0.23742 | 0.4219 | 0.2153 | True |
| GSM9338175_SSC5_visium | *CASP4* | 301 | 808 | 0.41794 | 0.33962 | 0.07832 | 0.3953 | 0.2908 | True |
| GSM9338175_SSC5_visium | *GSDMD* | 301 | 808 | 0.20815 | 0.37194 | 0.16379 | 0.299 | 0.3156 | True |
| GSM9338176_SSC6_visium | *NLRP1* | 369 | 453 | 0.34795 | 0.333 | 0.01495 | 0.1951 | 0.2362 | True |

#### Supplementary Table S3. Direct four-gene Visium replication (continued)

| Visium sample | Gene | n epi | n derm | Epi mean log1p | Derm mean log1p | Expected-dir score | Epi detect frac | Derm detect frac | Eligible |
|---|---|---|---|---|---|---|---|---|---|
| GSM9338176_SSC6_visium | *PYCARD* | 369 | 453 | 0.14018 | 0.09632 | 0.04386 | 0.1463 | 0.0993 | True |
| GSM9338176_SSC6_visium | *CASP4* | 369 | 453 | 0.18032 | 0.21766 | -0.03735 | 0.1518 | 0.1832 | True |
| GSM9338176_SSC6_visium | *GSDMD* | 369 | 453 | 0.08728 | 0.25371 | 0.16643 | 0.1138 | 0.1987 | True |
| GSM9338177_SSC7_visium | *NLRP1* | 374 | 743 | 0.34363 | 0.22231 | 0.12133 | 0.2406 | 0.1548 | True |
| GSM9338177_SSC7_visium | *PYCARD* | 374 | 743 | 0.1208 | 0.09904 | 0.02176 | 0.1497 | 0.0834 | True |
| GSM9338177_SSC7_visium | *CASP4* | 374 | 743 | 0.2266 | 0.20716 | 0.01944 | 0.2139 | 0.1561 | True |
| GSM9338177_SSC7_visium | *GSDMD* | 374 | 743 | 0.0735 | 0.12881 | 0.05531 | 0.107 | 0.1131 | True |
| GSM9338178_SSC8_visium | *NLRP1* | 174 | 568 | 0.46249 | 0.20534 | 0.25715 | 0.2931 | 0.1056 | True |
| GSM9338178_SSC8_visium | *PYCARD* | 174 | 568 | 0.12578 | 0.07301 | 0.05277 | 0.1207 | 0.0405 | True |
| GSM9338178_SSC8_visium | *CASP4* | 174 | 568 | 0.21891 | 0.14472 | 0.07419 | 0.1667 | 0.081 | True |
| GSM9338178_SSC8_visium | *GSDMD* | 174 | 568 | 0.23527 | 0.23033 | -0.00494 | 0.1839 | 0.1215 | True |
| GSM9338179_SSC9_visium | *NLRP1* | 415 | 869 | 0.35547 | 0.15843 | 0.19704 | 0.2072 | 0.1024 | True |
| GSM9338179_SSC9_visium | *PYCARD* | 415 | 869 | 0.12503 | 0.0562 | 0.06883 | 0.0892 | 0.0483 | True |
| GSM9338179_SSC9_visium | *CASP4* | 415 | 869 | 0.09689 | 0.18418 | -0.08729 | 0.0964 | 0.1197 | True |
| GSM9338179_SSC9_visium | *GSDMD* | 415 | 869 | 0.08445 | 0.17781 | 0.09336 | 0.0795 | 0.1139 | True |
| GSM9338180_SSC10_visium | *NLRP1* | 220 | 598 | 0.60528 | 0.43401 | 0.17127 | 0.6136 | 0.3244 | True |
| GSM9338180_SSC10_visium | *PYCARD* | 220 | 598 | 0.51036 | 0.13341 | 0.37694 | 0.5773 | 0.1204 | True |
| GSM9338180_SSC10_visium | *CASP4* | 220 | 598 | 0.42913 | 0.29652 | 0.13262 | 0.5545 | 0.2492 | True |
| GSM9338180_SSC10_visium | *GSDMD* | 220 | 598 | 0.26631 | 0.46479 | 0.19848 | 0.4409 | 0.3361 | True |

Intervals are 95% percentile bootstrap intervals for the median expected-direction contrast, using 10,000 section-resampling draws (seed 42); the endpoints are the 2.5th and 97.5th percentiles with linear interpolation. The displayed interval endpoints are retained from the frozen analysis.

#### Supplementary Table S4. Donor cell-class pseudobulk and class contrasts

| Floor | Group | Gene | Role | n | Median expected-direction contrast | + | 0 | − | P | BH q |
|---|---|---|---|---|---|---|---|---|---|---|
| 20 | SSc | *NLRP1* | primary_formal | 10 | -0.2858 | 1 | 0 | 9 | 0.02148438 | 0.02864583 |
| 20 | SSc | *PYCARD* | primary_formal | 10 | 1.7482 | 10 | 0 | 0 | 0.00195312 | 0.00390625 |
| 20 | SSc | *CASP4* | primary_formal | 10 | -0.24 | 3 | 0 | 7 | 0.34375 | 0.34375 |
| 20 | SSc | *GSDMD* | primary_formal | 10 | 2.5943 | 10 | 0 | 0 | 0.00195312 | 0.00390625 |
| 20 | Control | *NLRP1* | HC_descriptive_only | 4 | -0.5873 | | | | | |
| 20 | Control | *PYCARD* | HC_descriptive_only | 4 | 1.21 | | | | | |
| 20 | Control | *CASP4* | HC_descriptive_only | 4 | -0.1327 | | | | | |
| 20 | Control | *GSDMD* | HC_descriptive_only | 4 | 1.8504 | | | | | |
| 50 | SSc | *NLRP1* | floor50_descriptive_sensitivity | 7 | -0.3458 | 0 | 0 | 7 | | |
| 50 | SSc | *PYCARD* | floor50_descriptive_sensitivity | 7 | 1.7345 | 7 | 0 | 0 | | |
| 50 | SSc | *CASP4* | floor50_descriptive_sensitivity | 7 | -0.4484 | 0 | 0 | 7 | | |
| 50 | SSc | *GSDMD* | floor50_descriptive_sensitivity | 7 | 2.0417 | 7 | 0 | 0 | | |

| Sample | Group | Class | Gene | Nuclei | Raw count | Library 18,703 | CPM | log2(CPM+1) | >=20 | >=50 |
|---|---|---|---|---|---|---|---|---|---|---|
| GSM9338143_SSC1_multiome | SSc | keratinocyte | *NLRP1* | 2299 | 1291 | 1.66e+07 | 77.7742 | 6.2997 | True | True |
| GSM9338143_SSC1_multiome | SSc | keratinocyte | *PYCARD* | 2299 | 1344 | 1.66e+07 | 80.9671 | 6.357 | True | True |
| GSM9338143_SSC1_multiome | SSc | keratinocyte | *CASP4* | 2299 | 862 | 1.66e+07 | 51.9298 | 5.726 | True | True |
| GSM9338143_SSC1_multiome | SSc | keratinocyte | *GSDMD* | 2299 | 60 | 1.66e+07 | 3.6146 | 2.2062 | True | True |
| GSM9338143_SSC1_multiome | SSc | endothelial_strict | *NLRP1* | 664 | 898 | 6.24e+06 | 143.8903 | 7.1788 | True | True |
| GSM9338143_SSC1_multiome | SSc | endothelial_strict | *PYCARD* | 664 | 145 | 6.24e+06 | 23.234 | 4.599 | True | True |
| GSM9338143_SSC1_multiome | SSc | endothelial_strict | *CASP4* | 664 | 441 | 6.24e+06 | 70.6633 | 6.1632 | True | True |
| GSM9338143_SSC1_multiome | SSc | endothelial_strict | *GSDMD* | 664 | 322 | 6.24e+06 | 51.5954 | 5.7169 | True | True |
| GSM9338143_SSC1_multiome | SSc | fibroblast | *NLRP1* | 2113 | 422 | 8.53e+06 | 49.4801 | 5.6576 | True | True |
| GSM9338143_SSC1_multiome | SSc | fibroblast | *PYCARD* | 2113 | 205 | 8.53e+06 | 24.0365 | 4.646 | True | True |
| GSM9338143_SSC1_multiome | SSc | fibroblast | *CASP4* | 2113 | 413 | 8.53e+06 | 48.4248 | 5.6272 | True | True |

#### Supplementary Table S4. Donor cell-class pseudobulk and class contrasts (continued)

| Sample | Group | Class | Gene | Nuclei | Raw count | Library 18,703 | CPM | log2(CPM+1) | >=20 | >=50 |
|---|---|---|---|---|---|---|---|---|---|---|
| GSM9338143_SSC1_multiome | SSc | fibroblast | *GSDMD* | 2113 | 250 | 8.53e+06 | 29.3128 | 4.9219 | True | True |
| GSM9338145_SSC2_multiome | SSc | endothelial_strict | *NLRP1* | 419 | 619 | 4.86e+06 | 127.2723 | 7.0031 | True | True |
| GSM9338145_SSC2_multiome | SSc | endothelial_strict | *PYCARD* | 419 | 136 | 4.86e+06 | 27.9629 | 4.8561 | True | True |
| GSM9338145_SSC2_multiome | SSc | endothelial_strict | *CASP4* | 419 | 335 | 4.86e+06 | 68.8792 | 6.1268 | True | True |
| GSM9338145_SSC2_multiome | SSc | endothelial_strict | *GSDMD* | 419 | 247 | 4.86e+06 | 50.7856 | 5.6945 | True | True |
| GSM9338145_SSC2_multiome | SSc | keratinocyte | *NLRP1* | 2065 | 729 | 1.15e+07 | 63.3761 | 6.0085 | True | True |
| GSM9338145_SSC2_multiome | SSc | keratinocyte | *PYCARD* | 2065 | 890 | 1.15e+07 | 77.3727 | 6.2923 | True | True |
| GSM9338145_SSC2_multiome | SSc | keratinocyte | *CASP4* | 2065 | 386 | 1.15e+07 | 33.5572 | 5.1109 | True | True |
| GSM9338145_SSC2_multiome | SSc | keratinocyte | *GSDMD* | 2065 | 78 | 1.15e+07 | 6.781 | 2.96 | True | True |
| GSM9338145_SSC2_multiome | SSc | fibroblast | *NLRP1* | 656 | 190 | 3.14e+06 | 60.6023 | 5.9449 | True | True |
| GSM9338145_SSC2_multiome | SSc | fibroblast | *PYCARD* | 656 | 81 | 3.14e+06 | 25.8357 | 4.7461 | True | True |
| GSM9338145_SSC2_multiome | SSc | fibroblast | *CASP4* | 656 | 139 | 3.14e+06 | 44.3354 | 5.5026 | True | True |
| GSM9338145_SSC2_multiome | SSc | fibroblast | *GSDMD* | 656 | 59 | 3.14e+06 | 18.8186 | 4.3088 | True | True |
| GSM9338147_SSC3_multiome | SSc | keratinocyte | *NLRP1* | 2074 | 659 | 1.01e+07 | 65.0829 | 6.0462 | True | True |
| GSM9338147_SSC3_multiome | SSc | keratinocyte | *PYCARD* | 2074 | 268 | 1.01e+07 | 26.4677 | 4.7797 | True | True |
| GSM9338147_SSC3_multiome | SSc | keratinocyte | *CASP4* | 2074 | 423 | 1.01e+07 | 41.7755 | 5.4187 | True | True |
| GSM9338147_SSC3_multiome | SSc | keratinocyte | *GSDMD* | 2074 | 53 | 1.01e+07 | 5.2343 | 2.6402 | True | True |
| GSM9338147_SSC3_multiome | SSc | fibroblast | *NLRP1* | 309 | 68 | 1.2e+06 | 56.8492 | 5.8542 | True | True |
| GSM9338147_SSC3_multiome | SSc | fibroblast | *PYCARD* | 309 | 11 | 1.2e+06 | 9.1962 | 3.35 | True | True |
| GSM9338147_SSC3_multiome | SSc | fibroblast | *CASP4* | 309 | 75 | 1.2e+06 | 62.7013 | 5.9933 | True | True |
| GSM9338147_SSC3_multiome | SSc | fibroblast | *GSDMD* | 309 | 16 | 1.2e+06 | 13.3763 | 3.8456 | True | True |
| GSM9338147_SSC3_multiome | SSc | endothelial_strict | *NLRP1* | 104 | 93 | 9.1e+05 | 102.24 | 6.6899 | True | True |
| GSM9338147_SSC3_multiome | SSc | endothelial_strict | *PYCARD* | 104 | 22 | 9.1e+05 | 24.1858 | 4.6545 | True | True |
| GSM9338147_SSC3_multiome | SSc | endothelial_strict | *CASP4* | 104 | 103 | 9.1e+05 | 113.2336 | 6.8358 | True | True |
| GSM9338147_SSC3_multiome | SSc | endothelial_strict | *GSDMD* | 104 | 36 | 9.1e+05 | 39.5768 | 5.3426 | True | True |
| GSM9338149_SSC4_multiome | SSc | keratinocyte | *NLRP1* | 3576 | 2879 | 4.22e+07 | 68.2077 | 6.1129 | True | True |
| GSM9338149_SSC4_multiome | SSc | keratinocyte | *PYCARD* | 3576 | 568 | 4.22e+07 | 13.4567 | 3.8537 | True | True |

#### Supplementary Table S4. Donor cell-class pseudobulk and class contrasts (continued)

| Sample | Group | Class | Gene | Nuclei | Raw count | Library 18,703 | CPM | log2(CPM+1) | >=20 | >=50 |
|---|---|---|---|---|---|---|---|---|---|---|
| GSM9338149_SSC4_multiome | SSc | keratinocyte | *CASP4* | 3576 | 1481 | 4.22e+07 | 35.087 | 5.1734 | True | True |
| GSM9338149_SSC4_multiome | SSc | keratinocyte | *GSDMD* | 3576 | 64 | 4.22e+07 | 1.5163 | 1.3313 | True | True |
| GSM9338149_SSC4_multiome | SSc | endothelial_strict | *NLRP1* | 140 | 265 | 1.81e+06 | 146.7473 | 7.207 | True | True |
| GSM9338149_SSC4_multiome | SSc | endothelial_strict | *PYCARD* | 140 | 25 | 1.81e+06 | 13.8441 | 3.8918 | True | True |
| GSM9338149_SSC4_multiome | SSc | endothelial_strict | *CASP4* | 140 | 113 | 1.81e+06 | 62.5752 | 5.9904 | True | True |
| GSM9338149_SSC4_multiome | SSc | endothelial_strict | *GSDMD* | 140 | 56 | 1.81e+06 | 31.0107 | 5.0005 | True | True |
| GSM9338149_SSC4_multiome | SSc | fibroblast | *NLRP1* | 22 | 4 | 1.12e+05 | 35.799 | 5.2016 | True | False |
| GSM9338149_SSC4_multiome | SSc | fibroblast | *PYCARD* | 22 | 1 | 1.12e+05 | 8.9497 | 3.3147 | True | False |
| GSM9338149_SSC4_multiome | SSc | fibroblast | *CASP4* | 22 | 2 | 1.12e+05 | 17.8995 | 4.2403 | True | False |
| GSM9338149_SSC4_multiome | SSc | fibroblast | *GSDMD* | 22 | 1 | 1.12e+05 | 8.9497 | 3.3147 | True | False |
| GSM9338151_SSC5_multiome | SSc | keratinocyte | *NLRP1* | 2752 | 836 | 1.57e+07 | 53.2819 | 5.7624 | True | True |
| GSM9338151_SSC5_multiome | SSc | keratinocyte | *PYCARD* | 2752 | 2463 | 1.57e+07 | 156.9778 | 7.3036 | True | True |
| GSM9338151_SSC5_multiome | SSc | keratinocyte | *CASP4* | 2752 | 908 | 1.57e+07 | 57.8708 | 5.8795 | True | True |
| GSM9338151_SSC5_multiome | SSc | keratinocyte | *GSDMD* | 2752 | 195 | 1.57e+07 | 12.4282 | 3.7472 | True | True |
| GSM9338151_SSC5_multiome | SSc | endothelial_strict | *NLRP1* | 458 | 575 | 5.36e+06 | 107.3666 | 6.7598 | True | True |
| GSM9338151_SSC5_multiome | SSc | endothelial_strict | *PYCARD* | 458 | 180 | 5.36e+06 | 33.6104 | 5.1131 | True | True |
| GSM9338151_SSC5_multiome | SSc | endothelial_strict | *CASP4* | 458 | 563 | 5.36e+06 | 105.1259 | 6.7296 | True | True |
| GSM9338151_SSC5_multiome | SSc | endothelial_strict | *GSDMD* | 458 | 516 | 5.36e+06 | 96.3499 | 6.6051 | True | True |
| GSM9338151_SSC5_multiome | SSc | fibroblast | *NLRP1* | 756 | 197 | 3.54e+06 | 55.5934 | 5.8226 | True | True |
| GSM9338151_SSC5_multiome | SSc | fibroblast | *PYCARD* | 756 | 120 | 3.54e+06 | 33.864 | 5.1237 | True | True |
| GSM9338151_SSC5_multiome | SSc | fibroblast | *CASP4* | 756 | 266 | 3.54e+06 | 75.0653 | 6.2492 | True | True |
| GSM9338151_SSC5_multiome | SSc | fibroblast | *GSDMD* | 756 | 170 | 3.54e+06 | 47.974 | 5.6139 | True | True |
| GSM9338153_SSC6_multiome | SSc | keratinocyte | *NLRP1* | 1663 | 1562 | 2.14e+07 | 73.0394 | 6.2102 | True | True |
| GSM9338153_SSC6_multiome | SSc | keratinocyte | *PYCARD* | 1663 | 1988 | 2.14e+07 | 92.9592 | 6.554 | True | True |
| GSM9338153_SSC6_multiome | SSc | keratinocyte | *CASP4* | 1663 | 1271 | 2.14e+07 | 59.4321 | 5.9172 | True | True |
| GSM9338153_SSC6_multiome | SSc | keratinocyte | *GSDMD* | 1663 | 76 | 2.14e+07 | 3.5538 | 2.1871 | True | True |
| GSM9338153_SSC6_multiome | SSc | fibroblast | *NLRP1* | 33 | 4 | 1.69e+05 | 23.657 | 4.6239 | True | False |

#### Supplementary Table S4. Donor cell-class pseudobulk and class contrasts (continued)

| Sample | Group | Class | Gene | Nuclei | Raw count | Library 18,703 | CPM | log2(CPM+1) | >=20 | >=50 |
|---|---|---|---|---|---|---|---|---|---|---|
| GSM9338153_SSC6_multiome | SSc | fibroblast | *PYCARD* | 33 | 0 | 1.69e+05 | 0 | 0 | True | False |
| GSM9338153_SSC6_multiome | SSc | fibroblast | *CASP4* | 33 | 5 | 1.69e+05 | 29.5713 | 4.9341 | True | False |
| GSM9338153_SSC6_multiome | SSc | fibroblast | *GSDMD* | 33 | 4 | 1.69e+05 | 23.657 | 4.6239 | True | False |
| GSM9338153_SSC6_multiome | SSc | endothelial_strict | *NLRP1* | 62 | 98 | 7.97e+05 | 122.9839 | 6.954 | True | True |
| GSM9338153_SSC6_multiome | SSc | endothelial_strict | *PYCARD* | 62 | 28 | 7.97e+05 | 35.1383 | 5.1755 | True | True |
| GSM9338153_SSC6_multiome | SSc | endothelial_strict | *CASP4* | 62 | 63 | 7.97e+05 | 79.0611 | 6.323 | True | True |
| GSM9338153_SSC6_multiome | SSc | endothelial_strict | *GSDMD* | 62 | 44 | 7.97e+05 | 55.2173 | 5.8129 | True | True |
| GSM9338155_SSC7_multiome | SSc | fibroblast | *NLRP1* | 126 | 49 | 5.06e+05 | 96.8718 | 6.6128 | True | True |
| GSM9338155_SSC7_multiome | SSc | fibroblast | *PYCARD* | 126 | 8 | 5.06e+05 | 15.8158 | 4.0717 | True | True |
| GSM9338155_SSC7_multiome | SSc | fibroblast | *CASP4* | 126 | 26 | 5.06e+05 | 51.4014 | 5.7115 | True | True |
| GSM9338155_SSC7_multiome | SSc | fibroblast | *GSDMD* | 126 | 9 | 5.06e+05 | 17.7928 | 4.2321 | True | True |
| GSM9338155_SSC7_multiome | SSc | keratinocyte | *NLRP1* | 56 | 36 | 5.84e+05 | 61.6736 | 5.9698 | True | True |
| GSM9338155_SSC7_multiome | SSc | keratinocyte | *PYCARD* | 56 | 58 | 5.84e+05 | 99.363 | 6.6491 | True | True |
| GSM9338155_SSC7_multiome | SSc | keratinocyte | *CASP4* | 56 | 42 | 5.84e+05 | 71.9526 | 6.1889 | True | True |
| GSM9338155_SSC7_multiome | SSc | keratinocyte | *GSDMD* | 56 | 1 | 5.84e+05 | 1.7132 | 1.44 | True | True |
| GSM9338155_SSC7_multiome | SSc | endothelial_strict | *NLRP1* | 20 | 19 | 2.42e+05 | 78.5829 | 6.3144 | True | False |
| GSM9338155_SSC7_multiome | SSc | endothelial_strict | *PYCARD* | 20 | 1 | 2.42e+05 | 4.1359 | 2.3606 | True | False |
| GSM9338155_SSC7_multiome | SSc | endothelial_strict | *CASP4* | 20 | 12 | 2.42e+05 | 49.6313 | 5.662 | True | False |
| GSM9338155_SSC7_multiome | SSc | endothelial_strict | *GSDMD* | 20 | 9 | 2.42e+05 | 37.2235 | 5.2564 | True | False |
| GSM9338157_SSC8_multiome | SSc | keratinocyte | *NLRP1* | 2137 | 1402 | 2.25e+07 | 62.2188 | 5.9823 | True | True |
| GSM9338157_SSC8_multiome | SSc | keratinocyte | *PYCARD* | 2137 | 2956 | 2.25e+07 | 131.1831 | 7.0464 | True | True |
| GSM9338157_SSC8_multiome | SSc | keratinocyte | *CASP4* | 2137 | 1157 | 2.25e+07 | 51.346 | 5.71 | True | True |
| GSM9338157_SSC8_multiome | SSc | keratinocyte | *GSDMD* | 2137 | 370 | 2.25e+07 | 16.4201 | 4.1227 | True | True |
| GSM9338157_SSC8_multiome | SSc | fibroblast | *NLRP1* | 311 | 83 | 1.63e+06 | 51.0689 | 5.7024 | True | True |
| GSM9338157_SSC8_multiome | SSc | fibroblast | *PYCARD* | 311 | 44 | 1.63e+06 | 27.0727 | 4.8111 | True | True |
| GSM9338157_SSC8_multiome | SSc | fibroblast | *CASP4* | 311 | 78 | 1.63e+06 | 47.9925 | 5.6145 | True | True |
| GSM9338157_SSC8_multiome | SSc | fibroblast | *GSDMD* | 311 | 49 | 1.63e+06 | 30.1491 | 4.9611 | True | True |

#### Supplementary Table S4. Donor cell-class pseudobulk and class contrasts (continued)

| Sample | Group | Class | Gene | Nuclei | Raw count | Library 18,703 | CPM | log2(CPM+1) | >=20 | >=50 |
|---|---|---|---|---|---|---|---|---|---|---|
| GSM9338157_SSC8_multiome | SSc | endothelial_strict | *NLRP1* | 224 | 374 | 2.88e+06 | 129.9805 | 7.0332 | True | True |
| GSM9338157_SSC8_multiome | SSc | endothelial_strict | *PYCARD* | 224 | 66 | 2.88e+06 | 22.9377 | 4.5812 | True | True |
| GSM9338157_SSC8_multiome | SSc | endothelial_strict | *CASP4* | 224 | 240 | 2.88e+06 | 83.41 | 6.3993 | True | True |
| GSM9338157_SSC8_multiome | SSc | endothelial_strict | *GSDMD* | 224 | 202 | 2.88e+06 | 70.2034 | 6.1539 | True | True |
| GSM9338159_SSC9_multiome | SSc | keratinocyte | *NLRP1* | 3892 | 3244 | 2.9e+07 | 111.912 | 6.8191 | True | True |
| GSM9338159_SSC9_multiome | SSc | keratinocyte | *PYCARD* | 3892 | 2112 | 2.9e+07 | 72.8601 | 6.2067 | True | True |
| GSM9338159_SSC9_multiome | SSc | keratinocyte | *CASP4* | 3892 | 1410 | 2.9e+07 | 48.6424 | 5.6335 | True | True |
| GSM9338159_SSC9_multiome | SSc | keratinocyte | *GSDMD* | 3892 | 350 | 2.9e+07 | 12.0744 | 3.7087 | True | True |
| GSM9338159_SSC9_multiome | SSc | endothelial_strict | *NLRP1* | 839 | 849 | 5.53e+06 | 153.4285 | 7.2708 | True | True |
| GSM9338159_SSC9_multiome | SSc | endothelial_strict | *PYCARD* | 839 | 115 | 5.53e+06 | 20.7824 | 4.4451 | True | True |
| GSM9338159_SSC9_multiome | SSc | endothelial_strict | *CASP4* | 839 | 598 | 5.53e+06 | 108.0686 | 6.7691 | True | True |
| GSM9338159_SSC9_multiome | SSc | endothelial_strict | *GSDMD* | 839 | 242 | 5.53e+06 | 43.7334 | 5.4833 | True | True |
| GSM9338159_SSC9_multiome | SSc | fibroblast | *NLRP1* | 512 | 187 | 2.12e+06 | 88.28 | 6.4803 | True | True |
| GSM9338159_SSC9_multiome | SSc | fibroblast | *PYCARD* | 512 | 44 | 2.12e+06 | 20.7718 | 4.4444 | True | True |
| GSM9338159_SSC9_multiome | SSc | fibroblast | *CASP4* | 512 | 87 | 2.12e+06 | 41.0715 | 5.3948 | True | True |
| GSM9338159_SSC9_multiome | SSc | fibroblast | *GSDMD* | 512 | 39 | 2.12e+06 | 18.4113 | 4.2788 | True | True |
| GSM9338161_SSC10_multiome | SSc | keratinocyte | *NLRP1* | 2401 | 1650 | 2.48e+07 | 66.6459 | 6.0799 | True | True |
| GSM9338161_SSC10_multiome | SSc | keratinocyte | *PYCARD* | 2401 | 2520 | 2.48e+07 | 101.7864 | 6.6835 | True | True |
| GSM9338161_SSC10_multiome | SSc | keratinocyte | *CASP4* | 2401 | 1465 | 2.48e+07 | 59.1735 | 5.9111 | True | True |
| GSM9338161_SSC10_multiome | SSc | keratinocyte | *GSDMD* | 2401 | 79 | 2.48e+07 | 3.1909 | 2.0673 | True | True |
| GSM9338161_SSC10_multiome | SSc | endothelial_strict | *NLRP1* | 131 | 154 | 1.19e+06 | 129.7603 | 7.0308 | True | True |
| GSM9338161_SSC10_multiome | SSc | endothelial_strict | *PYCARD* | 131 | 47 | 1.19e+06 | 39.6022 | 5.3435 | True | True |
| GSM9338161_SSC10_multiome | SSc | endothelial_strict | *CASP4* | 131 | 92 | 1.19e+06 | 77.5191 | 6.295 | True | True |
| GSM9338161_SSC10_multiome | SSc | endothelial_strict | *GSDMD* | 131 | 61 | 1.19e+06 | 51.3985 | 5.7115 | True | True |
| GSM9338161_SSC10_multiome | SSc | fibroblast | *NLRP1* | 344 | 152 | 2.74e+06 | 55.521 | 5.8207 | True | True |
| GSM9338161_SSC10_multiome | SSc | fibroblast | *PYCARD* | 344 | 83 | 2.74e+06 | 30.3174 | 4.9689 | True | True |
| GSM9338161_SSC10_multiome | SSc | fibroblast | *CASP4* | 344 | 160 | 2.74e+06 | 58.4432 | 5.8934 | True | True |

#### Supplementary Table S4. Donor cell-class pseudobulk and class contrasts (continued)

| Sample | Group | Class | Gene | Nuclei | Raw count | Library 18,703 | CPM | log2(CPM+1) | >=20 | >=50 |
|---|---|---|---|---|---|---|---|---|---|---|
| GSM9338161_SSC10_multiome | SSc | fibroblast | *GSDMD* | 344 | 55 | 2.74e+06 | 20.0898 | 4.3985 | True | True |
| GSM9338163_HC1_multiome | Control | fibroblast | *NLRP1* | 4501 | 423 | 6.73e+06 | 62.8255 | 5.9961 | True | True |
| GSM9338163_HC1_multiome | Control | fibroblast | *PYCARD* | 4501 | 175 | 6.73e+06 | 25.9916 | 4.7544 | True | True |
| GSM9338163_HC1_multiome | Control | fibroblast | *CASP4* | 4501 | 316 | 6.73e+06 | 46.9335 | 5.583 | True | True |
| GSM9338163_HC1_multiome | Control | fibroblast | *GSDMD* | 4501 | 84 | 6.73e+06 | 12.476 | 3.7523 | True | True |
| GSM9338163_HC1_multiome | Control | keratinocyte | *NLRP1* | 1013 | 251 | 4.18e+06 | 59.9919 | 5.9305 | True | True |
| GSM9338163_HC1_multiome | Control | keratinocyte | *PYCARD* | 1013 | 184 | 4.18e+06 | 43.9782 | 5.4912 | True | True |
| GSM9338163_HC1_multiome | Control | keratinocyte | *CASP4* | 1013 | 196 | 4.18e+06 | 46.8463 | 5.5803 | True | True |
| GSM9338163_HC1_multiome | Control | keratinocyte | *GSDMD* | 1013 | 18 | 4.18e+06 | 4.3022 | 2.4066 | True | True |
| GSM9338163_HC1_multiome | Control | endothelial_strict | *NLRP1* | 133 | 54 | 5.13e+05 | 105.2234 | 6.731 | True | True |
| GSM9338163_HC1_multiome | Control | endothelial_strict | *PYCARD* | 133 | 13 | 5.13e+05 | 25.3316 | 4.7187 | True | True |
| GSM9338163_HC1_multiome | Control | endothelial_strict | *CASP4* | 133 | 32 | 5.13e+05 | 62.3546 | 5.9854 | True | True |
| GSM9338163_HC1_multiome | Control | endothelial_strict | *GSDMD* | 133 | 12 | 5.13e+05 | 23.383 | 4.6078 | True | True |
| GSM9338165_HC2_multiome | Control | fibroblast | *NLRP1* | 2565 | 351 | 4.85e+06 | 72.2994 | 6.1957 | True | True |
| GSM9338165_HC2_multiome | Control | fibroblast | *PYCARD* | 2565 | 132 | 4.85e+06 | 27.1895 | 4.8171 | True | True |
| GSM9338165_HC2_multiome | Control | fibroblast | *CASP4* | 2565 | 251 | 4.85e+06 | 51.7013 | 5.7198 | True | True |
| GSM9338165_HC2_multiome | Control | fibroblast | *GSDMD* | 2565 | 101 | 4.85e+06 | 20.8041 | 4.4465 | True | True |
| GSM9338165_HC2_multiome | Control | keratinocyte | *NLRP1* | 1296 | 213 | 4.42e+06 | 48.1469 | 5.619 | True | True |
| GSM9338165_HC2_multiome | Control | keratinocyte | *PYCARD* | 1296 | 239 | 4.42e+06 | 54.024 | 5.782 | True | True |
| GSM9338165_HC2_multiome | Control | keratinocyte | *CASP4* | 1296 | 214 | 4.42e+06 | 48.373 | 5.6256 | True | True |
| GSM9338165_HC2_multiome | Control | keratinocyte | *GSDMD* | 1296 | 24 | 4.42e+06 | 5.425 | 2.6837 | True | True |
| GSM9338165_HC2_multiome | Control | endothelial_strict | *NLRP1* | 132 | 63 | 5.23e+05 | 120.4723 | 6.9245 | True | True |
| GSM9338165_HC2_multiome | Control | endothelial_strict | *PYCARD* | 132 | 9 | 5.23e+05 | 17.2103 | 4.1867 | True | True |
| GSM9338165_HC2_multiome | Control | endothelial_strict | *CASP4* | 132 | 24 | 5.23e+05 | 45.8942 | 5.5513 | True | True |
| GSM9338165_HC2_multiome | Control | endothelial_strict | *GSDMD* | 132 | 10 | 5.23e+05 | 19.1226 | 4.3307 | True | True |
| GSM9338167_HC3_multiome | Control | keratinocyte | *NLRP1* | 3054 | 836 | 1.56e+07 | 53.5077 | 5.7684 | True | True |
| GSM9338167_HC3_multiome | Control | keratinocyte | *PYCARD* | 3054 | 1056 | 1.56e+07 | 67.5887 | 6.0999 | True | True |

#### Supplementary Table S4. Donor cell-class pseudobulk and class contrasts (continued)

| Sample | Group | Class | Gene | Nuclei | Raw count | Library 18,703 | CPM | log2(CPM+1) | >=20 | >=50 |
|---|---|---|---|---|---|---|---|---|---|---|
| GSM9338167_HC3_multiome | Control | keratinocyte | *CASP4* | 3054 | 621 | 1.56e+07 | 39.7468 | 5.3486 | True | True |
| GSM9338167_HC3_multiome | Control | keratinocyte | *GSDMD* | 3054 | 61 | 1.56e+07 | 3.9043 | 2.294 | True | True |
| GSM9338167_HC3_multiome | Control | fibroblast | *NLRP1* | 1769 | 247 | 4.44e+06 | 55.5934 | 5.8226 | True | True |
| GSM9338167_HC3_multiome | Control | fibroblast | *PYCARD* | 1769 | 105 | 4.44e+06 | 23.6328 | 4.6225 | True | True |
| GSM9338167_HC3_multiome | Control | fibroblast | *CASP4* | 1769 | 159 | 4.44e+06 | 35.7868 | 5.2011 | True | True |
| GSM9338167_HC3_multiome | Control | fibroblast | *GSDMD* | 1769 | 67 | 4.44e+06 | 15.08 | 4.0072 | True | True |
| GSM9338167_HC3_multiome | Control | endothelial_strict | *NLRP1* | 185 | 82 | 1.02e+06 | 80.5919 | 6.3504 | True | True |
| GSM9338167_HC3_multiome | Control | endothelial_strict | *PYCARD* | 185 | 39 | 1.02e+06 | 38.3303 | 5.2976 | True | True |
| GSM9338167_HC3_multiome | Control | endothelial_strict | *CASP4* | 185 | 49 | 1.02e+06 | 48.1586 | 5.6194 | True | True |
| GSM9338167_HC3_multiome | Control | endothelial_strict | *GSDMD* | 185 | 21 | 1.02e+06 | 20.6394 | 4.4356 | True | True |
| GSM9338169_HC4_multiome | Control | fibroblast | *NLRP1* | 1652 | 359 | 4.83e+06 | 74.3198 | 6.235 | True | True |
| GSM9338169_HC4_multiome | Control | fibroblast | *PYCARD* | 1652 | 155 | 4.83e+06 | 32.0879 | 5.0482 | True | True |
| GSM9338169_HC4_multiome | Control | fibroblast | *CASP4* | 1652 | 245 | 4.83e+06 | 50.7196 | 5.6926 | True | True |
| GSM9338169_HC4_multiome | Control | fibroblast | *GSDMD* | 1652 | 125 | 4.83e+06 | 25.8774 | 4.7483 | True | True |
| GSM9338169_HC4_multiome | Control | keratinocyte | *NLRP1* | 3008 | 1101 | 1.87e+07 | 58.9537 | 5.9058 | True | True |
| GSM9338169_HC4_multiome | Control | keratinocyte | *PYCARD* | 3008 | 1454 | 1.87e+07 | 77.8553 | 6.3011 | True | True |
| GSM9338169_HC4_multiome | Control | keratinocyte | *CASP4* | 3008 | 849 | 1.87e+07 | 45.4602 | 5.5379 | True | True |

#### Supplementary Table S4. Donor cell-class pseudobulk and class contrasts (continued)

| Sample | Group | Class | Gene | Nuclei | Raw count | Library 18,703 | CPM | log2(CPM+1) | >=20 | >=50 |
|---|---|---|---|---|---|---|---|---|---|---|
| GSM9338169_HC4_multiome | Control | keratinocyte | *GSDMD* | 3008 | 87 | 1.87e+07 | 4.6585 | 2.5004 | True | True |
| GSM9338169_HC4_multiome | Control | endothelial_strict | *NLRP1* | 656 | 760 | 5.74e+06 | 132.4404 | 7.0601 | True | True |
| GSM9338169_HC4_multiome | Control | endothelial_strict | *PYCARD* | 656 | 159 | 5.74e+06 | 27.7079 | 4.8434 | True | True |
| GSM9338169_HC4_multiome | Control | endothelial_strict | *CASP4* | 656 | 453 | 5.74e+06 | 78.9415 | 6.3209 | True | True |
| GSM9338169_HC4_multiome | Control | endothelial_strict | *GSDMD* | 656 | 275 | 5.74e+06 | 47.9225 | 5.6124 | True | True |

K, F and E denote donor-specific keratinocyte, fibroblast and endothelial_strict log2(CPM+1) values, respectively. The expected-direction contrast is K − (F + E)/2 for *NLRP1*, *PYCARD* and *CASP4*, and (F + E)/2 − K for *GSDMD*. The reported median summarizes these donor contrasts; +, 0 and − count positive, zero and negative contrasts. This contrast averages F and E after log transformation and is distinct from the linear-CPM mixture reconstruction.

Raw count is the sum of target transcripts within a donor and cell class. Library 18,703 is the summed library count over the same 18,703 retained features for that donor and class; CPM = raw count / library count × 10^6. Floors 20 and 50 require at least 20 and 50 nuclei, respectively, in each of the three classes. HC and floor-50 results are descriptive; blank P and q cells denote tests that were not performed.

#### Supplementary Table S5. Composition-adjustment status and diagnostics

| Legacy ID | Corrected ID | n epi | n derm | Eligible | Reason | Cond no. | VIF layer | VIF K | VIF F | VIF E | Max VIF |
|---|---|---|---|---|---|---|---|---|---|---|---|
| HC01 | HC01 | 398 | 1443 | True | PASS | 4.258 | 4.093 | 4.261 | 1.202 | 1.26 | 4.261 |
| HC02 | HC02 | 160 | 484 | True | PASS | 4.311 | 4.066 | 4.159 | 1.299 | 1.257 | 4.159 |
| HC03 | HC03 | 614 | 1191 | False | NOT_ESTIMABLE_COLLINEARITY | 7.557 | 6.521 | 12.641 | 4.914 | 2.793 | 12.641 |
| HC05 | HC05 | 572 | 187 | True | PASS | 2.429 | 1.333 | 1.888 | 1.374 | 1.407 | 1.888 |
| SSc-HL01 | SSc-HL01 | 99 | 354 | True | PASS | 5.364 | 4.569 | 6.706 | 2.356 | 1.551 | 6.706 |
| SSc-HL05 | SSc-HL05 | 71 | 414 | False | NOT_ESTIMABLE_COLLINEARITY | 8.384 | 10.055 | 15.545 | 4.158 | 1.661 | 15.545 |
| SSc-HL06 | SSc-HL06 | 69 | 539 | True | PASS | 5.362 | 5.573 | 6.723 | 1.555 | 1.601 | 6.723 |
| SSc-HL11 | SSc-HL11 | 68 | 604 | True | PASS | 6.328 | 5.654 | 9.519 | 2.916 | 1.548 | 9.519 |
| SSc-HL13 | SSc-HL13 | 73 | 477 | False | NOT_ESTIMABLE_COLLINEARITY | 7.907 | 12.041 | 13.545 | 1.908 | 1.65 | 13.545 |
| SSc-HL25 | SSc-HL25 | 83 | 665 | False | NOT_ESTIMABLE_COLLINEARITY | 7.534 | 10.474 | 12.533 | 2.807 | 1.648 | 12.533 |
| SSc-HL33 | SSc-HL35 | 61 | 712 | False | NOT_ESTIMABLE_COLLINEARITY | 7.219 | 10.282 | 11.63 | 1.766 | 1.446 | 11.63 |
| SSc-HL35 | SSc4994 | 133 | 1509 | True | PASS | 4.518 | 4.694 | 5.097 | 1.482 | 1.338 | 5.097 |
| SSc4994 | SSc5380 | 239 | 645 | True | PASS | 5.413 | 4.715 | 6.854 | 2.31 | 1.738 | 6.854 |

| Gene | Route | Status | n | Median | + | 0 | − | P | BH q | Interpretation |
|---|---|---|---|---|---|---|---|---|---|---|
| *NLRP1* | simultaneous_direct | NOT_ESTIMABLE_COLLINEARITY | 8 | -0.05445 | 3 | 0 | 5 | | | No spot-level P values. Dermis context and inactive sensitivity routes are descriptive only. |
| *PYCARD* | simultaneous_direct | NOT_ESTIMABLE_COLLINEARITY | 8 | -0.05901 | 4 | 0 | 4 | | | No spot-level P values. Dermis context and inactive sensitivity routes are descriptive only. |
| *CASP4* | simultaneous_direct | NOT_ESTIMABLE_COLLINEARITY | 8 | -0.00213 | 4 | 0 | 4 | | | No spot-level P values. Dermis context and inactive sensitivity routes are descriptive only. |
| *GSDMD* | simultaneous_direct | NOT_ESTIMABLE_COLLINEARITY | 8 | 0.00895 | 5 | 0 | 3 | | | No spot-level P values. Dermis context and inactive sensitivity routes are descriptive only. |

#### Supplementary Table S5. Composition-adjustment status and diagnostics (continued)

| Gene | Route | Status | n | Median | + | 0 | − | P | BH q | Interpretation |
|---|---|---|---|---|---|---|---|---|---|---|
| *NLRP1* | two_stage_residual | ESTIMABLE_FORMAL | 13 | 0.0024 | 7 | 0 | 6 | 1 | 1 | No spot-level P values. Dermis context and inactive sensitivity routes are descriptive only. |
| *PYCARD* | two_stage_residual | ESTIMABLE_FORMAL | 13 | -0.00509 | 4 | 0 | 9 | 0.2668457 | 1 | No spot-level P values. Dermis context and inactive sensitivity routes are descriptive only. |
| *CASP4* | two_stage_residual | ESTIMABLE_FORMAL | 13 | 0.00564 | 7 | 0 | 6 | 1 | 1 | No spot-level P values. Dermis context and inactive sensitivity routes are descriptive only. |
| *GSDMD* | two_stage_residual | ESTIMABLE_FORMAL | 13 | 0.00181 | 8 | 0 | 5 | 0.58105469 | 1 | No spot-level P values. Dermis context and inactive sensitivity routes are descriptive only. |
| *NLRP1* | dermis_endothelial_context | ESTIMABLE_DESCRIPTIVE | 13 | 0.16782 | 13 | 0 | 0 | | | No spot-level P values. Dermis context and inactive sensitivity routes are descriptive only. |
| *PYCARD* | dermis_endothelial_context | ESTIMABLE_DESCRIPTIVE | 13 | 0.06883 | 12 | 0 | 1 | | | No spot-level P values. Dermis context and inactive sensitivity routes are descriptive only. |
| *CASP4* | dermis_endothelial_context | ESTIMABLE_DESCRIPTIVE | 13 | 0.14111 | 13 | 0 | 0 | | | No spot-level P values. Dermis context and inactive sensitivity routes are descriptive only. |
| *GSDMD* | dermis_endothelial_context | ESTIMABLE_DESCRIPTIVE | 13 | 0.1773 | 13 | 0 | 0 | | | No spot-level P values. Dermis context and inactive sensitivity routes are descriptive only. |

| Module | Status | Reason | Replacement score | P value |
|---|---|---|---|---|
| A2 | NOT_ESTIMABLE_ENDOTHELIAL_CONTEXT | The only available endothelium_score is the expression-derived layer-generator score; no independent endothelial context score was created. | False | |

#### Supplementary Table S6. All M1/M2 donor-gene predictions and directional categories

| Donor | Visium sample | snMultiome sample | Model | Gene | Observed E−D | O sign | Proxy E−D | P sign | Exact | Category | Inference |
|---|---|---|---|---|---|---|---|---|---|---|---|
| SSc1 | GSM9338171_SSC1_visium | GSM9338143_SSC1_multiome | M1 | *NLRP1* | 0.21671 | 1 | 0.10876 | 1 | True | DIRECTION_REPRODUCED | NONE |
| SSc1 | GSM9338171_SSC1_visium | GSM9338143_SSC1_multiome | M1 | *PYCARD* | 0.18047 | 1 | 1.72211 | 1 | True | DIRECTION_REPRODUCED | NONE |
| SSc1 | GSM9338171_SSC1_visium | GSM9338143_SSC1_multiome | M1 | *CASP4* | 0.1066 | 1 | -0.04857 | -1 | False | OBSERVED_MORE_EPIDERMAL_THAN_PROXY | NONE |
| SSc1 | GSM9338171_SSC1_visium | GSM9338143_SSC1_multiome | M1 | *GSDMD* | -0.10055 | -1 | -2.94925 | -1 | True | DIRECTION_REPRODUCED | NONE |
| SSc1 | GSM9338171_SSC1_visium | GSM9338143_SSC1_multiome | M2 | *NLRP1* | 0.21671 | 1 | -0.31042 | -1 | False | OBSERVED_MORE_EPIDERMAL_THAN_PROXY | NONE |
| SSc1 | GSM9338171_SSC1_visium | GSM9338143_SSC1_multiome | M2 | *PYCARD* | 0.18047 | 1 | 1.73432 | 1 | True | DIRECTION_REPRODUCED | NONE |
| SSc1 | GSM9338171_SSC1_visium | GSM9338143_SSC1_multiome | M2 | *CASP4* | 0.1066 | 1 | -0.19391 | -1 | False | OBSERVED_MORE_EPIDERMAL_THAN_PROXY | NONE |
| SSc1 | GSM9338171_SSC1_visium | GSM9338143_SSC1_multiome | M2 | *GSDMD* | -0.10055 | -1 | -3.16724 | -1 | True | DIRECTION_REPRODUCED | NONE |
| SSc2 | GSM9338172_SSC2_visium | GSM9338145_SSC2_multiome | M1 | *NLRP1* | 0.37394 | 1 | -0.44421 | -1 | False | OBSERVED_MORE_EPIDERMAL_THAN_PROXY | NONE |
| SSc2 | GSM9338172_SSC2_visium | GSM9338145_SSC2_multiome | M1 | *PYCARD* | 0.1666 | 1 | 1.5023 | 1 | True | DIRECTION_REPRODUCED | NONE |
| SSc2 | GSM9338172_SSC2_visium | GSM9338145_SSC2_multiome | M1 | *CASP4* | 0.15939 | 1 | -0.66787 | -1 | False | OBSERVED_MORE_EPIDERMAL_THAN_PROXY | NONE |
| SSc2 | GSM9338172_SSC2_visium | GSM9338145_SSC2_multiome | M1 | *GSDMD* | 0.07922 | 1 | -2.05254 | -1 | False | OBSERVED_MORE_EPIDERMAL_THAN_PROXY | NONE |
| SSc2 | GSM9338172_SSC2_visium | GSM9338145_SSC2_multiome | M2 | *NLRP1* | 0.37394 | 1 | -0.56045 | -1 | False | OBSERVED_MORE_EPIDERMAL_THAN_PROXY | NONE |
| SSc2 | GSM9338172_SSC2_visium | GSM9338145_SSC2_multiome | M2 | *PYCARD* | 0.1666 | 1 | 1.49012 | 1 | True | DIRECTION_REPRODUCED | NONE |
| SSc2 | GSM9338172_SSC2_visium | GSM9338145_SSC2_multiome | M2 | *CASP4* | 0.15939 | 1 | -0.73727 | -1 | False | OBSERVED_MORE_EPIDERMAL_THAN_PROXY | NONE |
| SSc2 | GSM9338172_SSC2_visium | GSM9338145_SSC2_multiome | M2 | *GSDMD* | 0.07922 | 1 | -2.20202 | -1 | False | OBSERVED_MORE_EPIDERMAL_THAN_PROXY | NONE |

#### Supplementary Table S6. All M1/M2 donor-gene predictions and directional categories (continued)

| Donor | Visium sample | snMultiome sample | Model | Gene | Observed E−D | O sign | Proxy E−D | P sign | Exact | Category | Inference |
|---|---|---|---|---|---|---|---|---|---|---|---|
| SSc3 | GSM9338173_SSC3_visium | GSM9338147_SSC3_multiome | M1 | *NLRP1* | 0.14461 | 1 | -0.06815 | -1 | False | OBSERVED_MORE_EPIDERMAL_THAN_PROXY | NONE |
| SSc3 | GSM9338173_SSC3_visium | GSM9338147_SSC3_multiome | M1 | *PYCARD* | 0.14693 | 1 | 0.97532 | 1 | True | DIRECTION_REPRODUCED | NONE |
| SSc3 | GSM9338173_SSC3_visium | GSM9338147_SSC3_multiome | M1 | *CASP4* | 0.0953 | 1 | -0.83728 | -1 | False | OBSERVED_MORE_EPIDERMAL_THAN_PROXY | NONE |
| SSc3 | GSM9338173_SSC3_visium | GSM9338147_SSC3_multiome | M1 | *GSDMD* | -0.10109 | -1 | -1.75031 | -1 | True | DIRECTION_REPRODUCED | NONE |
| SSc3 | GSM9338173_SSC3_visium | GSM9338147_SSC3_multiome | M2 | *NLRP1* | 0.14461 | 1 | -0.28551 | -1 | False | OBSERVED_MORE_EPIDERMAL_THAN_PROXY | NONE |
| SSc3 | GSM9338173_SSC3_visium | GSM9338147_SSC3_multiome | M2 | *PYCARD* | 0.14693 | 1 | 0.63472 | 1 | True | DIRECTION_REPRODUCED | NONE |
| SSc3 | GSM9338173_SSC3_visium | GSM9338147_SSC3_multiome | M2 | *CASP4* | 0.0953 | 1 | -1.05649 | -1 | False | OBSERVED_MORE_EPIDERMAL_THAN_PROXY | NONE |
| SSc3 | GSM9338173_SSC3_visium | GSM9338147_SSC3_multiome | M2 | *GSDMD* | -0.10109 | -1 | -2.1399 | -1 | True | DIRECTION_REPRODUCED | NONE |
| SSc4 | GSM9338174_SSC4_visium | GSM9338149_SSC4_multiome | M1 | *NLRP1* | 0.16999 | 1 | -0.93895 | -1 | False | OBSERVED_MORE_EPIDERMAL_THAN_PROXY | NONE |
| SSc4 | GSM9338174_SSC4_visium | GSM9338149_SSC4_multiome | M1 | *PYCARD* | -0.03564 | -1 | 0.02794 | 1 | False | OBSERVED_MORE_DERMAL_THAN_PROXY | NONE |
| SSc4 | GSM9338174_SSC4_visium | GSM9338149_SSC4_multiome | M1 | *CASP4* | -0.04562 | -1 | -0.67229 | -1 | True | DIRECTION_REPRODUCED | NONE |
| SSc4 | GSM9338174_SSC4_visium | GSM9338149_SSC4_multiome | M1 | *GSDMD* | -0.09458 | -1 | -3.52744 | -1 | True | DIRECTION_REPRODUCED | NONE |
| SSc4 | GSM9338174_SSC4_visium | GSM9338149_SSC4_multiome | M2 | *NLRP1* | 0.16999 | 1 | -0.41498 | -1 | False | OBSERVED_MORE_EPIDERMAL_THAN_PROXY | NONE |
| SSc4 | GSM9338174_SSC4_visium | GSM9338149_SSC4_multiome | M2 | *PYCARD* | -0.03564 | -1 | 0.22176 | 1 | False | OBSERVED_MORE_DERMAL_THAN_PROXY | NONE |
| SSc4 | GSM9338174_SSC4_visium | GSM9338149_SSC4_multiome | M2 | *CASP4* | -0.04562 | -1 | -0.19247 | -1 | True | DIRECTION_REPRODUCED | NONE |
| SSc4 | GSM9338174_SSC4_visium | GSM9338149_SSC4_multiome | M2 | *GSDMD* | -0.09458 | -1 | -3.05968 | -1 | True | DIRECTION_REPRODUCED | NONE |

#### Supplementary Table S6. All M1/M2 donor-gene predictions and directional categories (continued)

| Donor | Visium sample | snMultiome sample | Model | Gene | Observed E−D | O sign | Proxy E−D | P sign | Exact | Category | Inference |
|---|---|---|---|---|---|---|---|---|---|---|---|
| SSc5 | GSM9338175_SSC5_visium | GSM9338151_SSC5_multiome | M1 | *NLRP1* | 0.07599 | 1 | -0.48791 | -1 | False | OBSERVED_MORE_EPIDERMAL_THAN_PROXY | NONE |
| SSc5 | GSM9338175_SSC5_visium | GSM9338151_SSC5_multiome | M1 | *PYCARD* | 0.23742 | 1 | 2.18388 | 1 | True | DIRECTION_REPRODUCED | NONE |
| SSc5 | GSM9338175_SSC5_visium | GSM9338151_SSC5_multiome | M1 | *CASP4* | 0.07832 | 1 | -0.57018 | -1 | False | OBSERVED_MORE_EPIDERMAL_THAN_PROXY | NONE |
| SSc5 | GSM9338175_SSC5_visium | GSM9338151_SSC5_multiome | M1 | *GSDMD* | -0.16379 | -1 | -2.32372 | -1 | True | DIRECTION_REPRODUCED | NONE |
| SSc5 | GSM9338175_SSC5_visium | GSM9338151_SSC5_multiome | M2 | *NLRP1* | 0.07599 | 1 | -0.60357 | -1 | False | OBSERVED_MORE_EPIDERMAL_THAN_PROXY | NONE |
| SSc5 | GSM9338175_SSC5_visium | GSM9338151_SSC5_multiome | M2 | *PYCARD* | 0.23742 | 1 | 2.18517 | 1 | True | DIRECTION_REPRODUCED | NONE |
| SSc5 | GSM9338175_SSC5_visium | GSM9338151_SSC5_multiome | M2 | *CASP4* | 0.07832 | 1 | -0.62983 | -1 | False | OBSERVED_MORE_EPIDERMAL_THAN_PROXY | NONE |
| SSc5 | GSM9338175_SSC5_visium | GSM9338151_SSC5_multiome | M2 | *GSDMD* | -0.16379 | -1 | -2.44583 | -1 | True | DIRECTION_REPRODUCED | NONE |
| SSc6 | GSM9338176_SSC6_visium | GSM9338153_SSC6_multiome | M1 | *NLRP1* | 0.01495 | 1 | -0.27329 | -1 | False | OBSERVED_MORE_EPIDERMAL_THAN_PROXY | NONE |
| SSc6 | GSM9338176_SSC6_visium | GSM9338153_SSC6_multiome | M1 | *PYCARD* | 0.04386 | 1 | 1.97307 | 1 | True | DIRECTION_REPRODUCED | NONE |
| SSc6 | GSM9338176_SSC6_visium | GSM9338153_SSC6_multiome | M1 | *CASP4* | -0.03735 | -1 | -0.05705 | -1 | True | DIRECTION_REPRODUCED | NONE |
| SSc6 | GSM9338176_SSC6_visium | GSM9338153_SSC6_multiome | M1 | *GSDMD* | -0.16643 | -1 | -3.31292 | -1 | True | DIRECTION_REPRODUCED | NONE |
| SSc6 | GSM9338176_SSC6_visium | GSM9338153_SSC6_multiome | M2 | *NLRP1* | 0.01495 | 1 | -0.00547 | -1 | False | OBSERVED_MORE_EPIDERMAL_THAN_PROXY | NONE |
| SSc6 | GSM9338176_SSC6_visium | GSM9338153_SSC6_multiome | M2 | *PYCARD* | 0.04386 | 1 | 2.33913 | 1 | True | DIRECTION_REPRODUCED | NONE |
| SSc6 | GSM9338176_SSC6_visium | GSM9338153_SSC6_multiome | M2 | *CASP4* | -0.03735 | -1 | 0.12761 | 1 | False | OBSERVED_MORE_DERMAL_THAN_PROXY | NONE |
| SSc6 | GSM9338176_SSC6_visium | GSM9338153_SSC6_multiome | M2 | *GSDMD* | -0.16643 | -1 | -3.15055 | -1 | True | DIRECTION_REPRODUCED | NONE |

#### Supplementary Table S6. All M1/M2 donor-gene predictions and directional categories (continued)

| Donor | Visium sample | snMultiome sample | Model | Gene | Observed E−D | O sign | Proxy E−D | P sign | Exact | Category | Inference |
|---|---|---|---|---|---|---|---|---|---|---|---|
| SSc7 | GSM9338177_SSC7_visium | GSM9338155_SSC7_multiome | M1 | *NLRP1* | 0.12133 | 1 | -0.60562 | -1 | False | OBSERVED_MORE_EPIDERMAL_THAN_PROXY | NONE |
| SSc7 | GSM9338177_SSC7_visium | GSM9338155_SSC7_multiome | M1 | *PYCARD* | 0.02176 | 1 | 2.72158 | 1 | True | DIRECTION_REPRODUCED | NONE |
| SSc7 | GSM9338177_SSC7_visium | GSM9338155_SSC7_multiome | M1 | *CASP4* | 0.01944 | 1 | 0.48405 | 1 | True | DIRECTION_REPRODUCED | NONE |
| SSc7 | GSM9338177_SSC7_visium | GSM9338155_SSC7_multiome | M1 | *GSDMD* | -0.05531 | -1 | -2.98324 | -1 | True | DIRECTION_REPRODUCED | NONE |
| SSc7 | GSM9338177_SSC7_visium | GSM9338155_SSC7_multiome | M2 | *NLRP1* | 0.12133 | 1 | -0.50152 | -1 | False | OBSERVED_MORE_EPIDERMAL_THAN_PROXY | NONE |
| SSc7 | GSM9338177_SSC7_visium | GSM9338155_SSC7_multiome | M2 | *PYCARD* | 0.02176 | 1 | 3.19282 | 1 | True | DIRECTION_REPRODUCED | NONE |
| SSc7 | GSM9338177_SSC7_visium | GSM9338155_SSC7_multiome | M2 | *CASP4* | 0.01944 | 1 | 0.50193 | 1 | True | DIRECTION_REPRODUCED | NONE |
| SSc7 | GSM9338177_SSC7_visium | GSM9338155_SSC7_multiome | M2 | *GSDMD* | -0.05531 | -1 | -3.39333 | -1 | True | DIRECTION_REPRODUCED | NONE |
| SSc8 | GSM9338178_SSC8_visium | GSM9338157_SSC8_multiome | M1 | *NLRP1* | 0.25715 | 1 | -0.42895 | -1 | False | OBSERVED_MORE_EPIDERMAL_THAN_PROXY | NONE |
| SSc8 | GSM9338178_SSC8_visium | GSM9338157_SSC8_multiome | M1 | *PYCARD* | 0.05277 | 1 | 2.32713 | 1 | True | DIRECTION_REPRODUCED | NONE |
| SSc8 | GSM9338178_SSC8_visium | GSM9338157_SSC8_multiome | M1 | *CASP4* | 0.07419 | 1 | -0.28596 | -1 | False | OBSERVED_MORE_EPIDERMAL_THAN_PROXY | NONE |
| SSc8 | GSM9338178_SSC8_visium | GSM9338157_SSC8_multiome | M1 | *GSDMD* | 0.00494 | 1 | -1.45986 | -1 | False | OBSERVED_MORE_EPIDERMAL_THAN_PROXY | NONE |
| SSc8 | GSM9338178_SSC8_visium | GSM9338157_SSC8_multiome | M2 | *NLRP1* | 0.25715 | 1 | -0.53381 | -1 | False | OBSERVED_MORE_EPIDERMAL_THAN_PROXY | NONE |
| SSc8 | GSM9338178_SSC8_visium | GSM9338157_SSC8_multiome | M2 | *PYCARD* | 0.05277 | 1 | 2.34566 | 1 | True | DIRECTION_REPRODUCED | NONE |
| SSc8 | GSM9338178_SSC8_visium | GSM9338157_SSC8_multiome | M2 | *CASP4* | 0.07419 | 1 | -0.34963 | -1 | False | OBSERVED_MORE_EPIDERMAL_THAN_PROXY | NONE |
| SSc8 | GSM9338178_SSC8_visium | GSM9338157_SSC8_multiome | M2 | *GSDMD* | 0.00494 | 1 | -1.55472 | -1 | False | OBSERVED_MORE_EPIDERMAL_THAN_PROXY | NONE |

#### Supplementary Table S6. All M1/M2 donor-gene predictions and directional categories (continued)

| Donor | Visium sample | snMultiome sample | Model | Gene | Observed E−D | O sign | Proxy E−D | P sign | Exact | Category | Inference |
|---|---|---|---|---|---|---|---|---|---|---|---|
| SSc9 | GSM9338179_SSC9_visium | GSM9338159_SSC9_multiome | M1 | *NLRP1* | 0.19704 | 1 | -0.20041 | -1 | False | OBSERVED_MORE_EPIDERMAL_THAN_PROXY | NONE |
| SSc9 | GSM9338179_SSC9_visium | GSM9338159_SSC9_multiome | M1 | *PYCARD* | 0.06883 | 1 | 1.7619 | 1 | True | DIRECTION_REPRODUCED | NONE |
| SSc9 | GSM9338179_SSC9_visium | GSM9338159_SSC9_multiome | M1 | *CASP4* | -0.08729 | -1 | -0.75328 | -1 | True | DIRECTION_REPRODUCED | NONE |
| SSc9 | GSM9338179_SSC9_visium | GSM9338159_SSC9_multiome | M1 | *GSDMD* | -0.09336 | -1 | -1.42625 | -1 | True | DIRECTION_REPRODUCED | NONE |
| SSc9 | GSM9338179_SSC9_visium | GSM9338159_SSC9_multiome | M2 | *NLRP1* | 0.19704 | 1 | -0.10996 | -1 | False | OBSERVED_MORE_EPIDERMAL_THAN_PROXY | NONE |
| SSc9 | GSM9338179_SSC9_visium | GSM9338159_SSC9_multiome | M2 | *PYCARD* | 0.06883 | 1 | 1.76198 | 1 | True | DIRECTION_REPRODUCED | NONE |
| SSc9 | GSM9338179_SSC9_visium | GSM9338159_SSC9_multiome | M2 | *CASP4* | -0.08729 | -1 | -0.60624 | -1 | True | DIRECTION_REPRODUCED | NONE |
| SSc9 | GSM9338179_SSC9_visium | GSM9338159_SSC9_multiome | M2 | *GSDMD* | -0.09336 | -1 | -1.29459 | -1 | True | DIRECTION_REPRODUCED | NONE |
| SSc10 | GSM9338180_SSC10_visium | GSM9338161_SSC10_multiome | M1 | *NLRP1* | 0.17127 | 1 | -0.18677 | -1 | False | OBSERVED_MORE_EPIDERMAL_THAN_PROXY | NONE |
| SSc10 | GSM9338180_SSC10_visium | GSM9338161_SSC10_multiome | M1 | *PYCARD* | 0.37694 | 1 | 1.60123 | 1 | True | DIRECTION_REPRODUCED | NONE |
| SSc10 | GSM9338180_SSC10_visium | GSM9338161_SSC10_multiome | M1 | *CASP4* | 0.13262 | 1 | -0.10473 | -1 | False | OBSERVED_MORE_EPIDERMAL_THAN_PROXY | NONE |

#### Supplementary Table S6. All M1/M2 donor-gene predictions and directional categories (continued)

| Donor | Visium sample | snMultiome sample | Model | Gene | Observed E−D | O sign | Proxy E−D | P sign | Exact | Category | Inference |
|---|---|---|---|---|---|---|---|---|---|---|---|
| SSc10 | GSM9338180_SSC10_visium | GSM9338161_SSC10_multiome | M1 | *GSDMD* | -0.19848 | -1 | -2.82631 | -1 | True | DIRECTION_REPRODUCED | NONE |
| SSc10 | GSM9338180_SSC10_visium | GSM9338161_SSC10_multiome | M2 | *NLRP1* | 0.17127 | 1 | -0.46913 | -1 | False | OBSERVED_MORE_EPIDERMAL_THAN_PROXY | NONE |
| SSc10 | GSM9338180_SSC10_visium | GSM9338161_SSC10_multiome | M2 | *PYCARD* | 0.37694 | 1 | 1.51519 | 1 | True | DIRECTION_REPRODUCED | NONE |
| SSc10 | GSM9338180_SSC10_visium | GSM9338161_SSC10_multiome | M2 | *CASP4* | 0.13262 | 1 | -0.19707 | -1 | False | OBSERVED_MORE_EPIDERMAL_THAN_PROXY | NONE |
| SSc10 | GSM9338180_SSC10_visium | GSM9338161_SSC10_multiome | M2 | *GSDMD* | -0.19848 | -1 | -3.13218 | -1 | True | DIRECTION_REPRODUCED | NONE |

Observed E−D uses natural-log1p Visium values. Proxy E−D uses log2(CPM+1) after mixing class CPM on the linear scale. O denotes observed and P denotes proxy; positive, zero and negative signs encode the corresponding directions. The two numerical scales must not be subtracted or interpreted as raw residuals, amplitude agreement or a proportion explained.

#### Supplementary Table S7. Directional concordance and leave-one-donor-out ranges

| Model | Gene | n | Exact | Discordant | Zero | O+ | O− | P+ | P− | Inference |
|---|---|---|---|---|---|---|---|---|---|---|
| M1 | *NLRP1* | 10 | 1 | 9 | 0 | 10 | 0 | 1 | 9 | NONE |
| M1 | *PYCARD* | 10 | 9 | 1 | 0 | 9 | 1 | 10 | 0 | NONE |
| M1 | *CASP4* | 10 | 4 | 6 | 0 | 7 | 3 | 1 | 9 | NONE |
| M1 | *GSDMD* | 10 | 8 | 2 | 0 | 2 | 8 | 0 | 10 | NONE |
| M2 | *NLRP1* | 10 | 0 | 10 | 0 | 10 | 0 | 0 | 10 | NONE |
| M2 | *PYCARD* | 10 | 9 | 1 | 0 | 9 | 1 | 10 | 0 | NONE |
| M2 | *CASP4* | 10 | 3 | 7 | 0 | 7 | 3 | 2 | 8 | NONE |
| M2 | *GSDMD* | 10 | 8 | 2 | 0 | 2 | 8 | 0 | 10 | NONE |

| Model | Gene | Minimum exact | Maximum exact | Denominator | Role |
|---|---|---|---|---|---|
| M1 | *NLRP1* | 0 | 1 | 9 | descriptive; no refit; no P value |
| M1 | *PYCARD* | 8 | 9 | 9 | descriptive; no refit; no P value |
| M1 | *CASP4* | 3 | 4 | 9 | descriptive; no refit; no P value |
| M1 | *GSDMD* | 7 | 8 | 9 | descriptive; no refit; no P value |
| M2 | *NLRP1* | 0 | 0 | 9 | descriptive; no refit; no P value |
| M2 | *PYCARD* | 8 | 9 | 9 | descriptive; no refit; no P value |
| M2 | *CASP4* | 2 | 3 | 9 | descriptive; no refit; no P value |
| M2 | *GSDMD* | 7 | 8 | 9 | descriptive; no refit; no P value |

The complete 80-row concordance/discordance ledger and all 80 leave-one-donor-out rows are available in the Supporting Data workbook; no favourable-only rows were selected. LOO denotes leave-one-donor-out descriptive influence summaries without refitting or formal inference; the denominator is nine donors.

#### Supplementary Table S8. Post hoc endothelial-relative-abundance sensitivity analysis of the dermal IFNγ-response–*GSDMD* association

| Report section | Legacy source label | Group | Dermal spots | Unadjusted rho | Historical score-adjusted rho | Cell2location-adjusted rho |
|---|---|---|---|---|---|---|
| HC01 | HC01 | HC | 1443 | -0.017 | -0.016 | -0.021 |
| HC02 | HC02 | HC | 484 | 0.081 | 0.077 | 0.065 |
| HC03 | HC03 | HC | 1191 | 0.092 | 0.071 | 0.051 |
| HC05 | HC05 | HC | 187 | 0.050 | 0.036 | 0.026 |
| SSc-HL01 | SSc-HL01 | SSc | 354 | 0.068 | 0.066 | 0.027 |
| SSc-HL05 | SSc-HL05 | SSc | 414 | 0.245 | 0.244 | 0.219 |
| SSc-HL06 | SSc-HL06 | SSc | 539 | 0.293 | 0.266 | 0.236 |
| SSc-HL11 | SSc-HL11 | SSc | 604 | 0.109 | 0.109 | 0.072 |
| SSc-HL13 | SSc-HL13 | SSc | 477 | 0.073 | 0.052 | 0.034 |
| SSc-HL25 | SSc-HL25 | SSc | 665 | 0.182 | 0.156 | 0.107 |
| SSc-HL35 | SSc-HL33 | SSc | 712 | 0.155 | 0.130 | 0.114 |
| SSc4994 | SSc-HL35 | SSc | 1509 | 0.132 | 0.107 | 0.094 |
| SSc5380 | SSc4994 | SSc | 645 | 0.059 | 0.060 | 0.036 |

| Model | Sections | Median rho | Positive / zero / negative | Range |
|---|---|---|---|---|
| Unadjusted | 13 | 0.092 | 12 / 0 / 1 | -0.017 to 0.293 |
| Historical score-adjusted | 13 | 0.077 | 12 / 0 / 1 | -0.016 to 0.266 |
| Cell2location-adjusted | 13 | 0.065 | 12 / 0 / 1 | -0.021 to 0.236 |

For the Cell2location-adjusted coefficients, the median was 0.065 (95% section-bootstrap CI 0.034–0.114), with positive values in 12 of 13 sections. The two-sided section-level Wilcoxon signed-rank statistic was 1 and P=0.00049 (unadjusted post hoc single test). The bootstrap interval concerns the median and is not the historical Fisher-z pooled interval. All 13 sections are shown; HC denotes healthy control. Legacy source labels are linked to the unchanged SampleCrosswalk in Supporting Data; no sections or numerical values were reassigned by row position. Historical score-adjusted values refer to endothelial_score, whose original construction remains unverified. Full-precision values and code provenance are provided in Supporting Data and Supplementary Software 1.

#### Supplementary Table S9. Dermal endothelial-weight sensitivity of exact sign concordance

Each entry is the number of concordant donors out of 10 for the observed Visium epidermis-minus-dermis sign and the restricted three-class proxy sign.

| Endothelial share w | NLRP1 | PYCARD | CASP4 | GSDMD | Interpretation |
|---|---|---|---|---|---|
| 0.00 | 8 | 9 | 4 | 8 | Fixed-weight grid |
| 0.05 | 7 | 9 | 3 | 8 | Fixed-weight grid |
| 0.10 | 7 | 9 | 2 | 8 | Fixed-weight grid |
| 0.15 | 5 | 9 | 3 | 8 | Fixed-weight grid |
| 0.20 | 4 | 9 | 2 | 8 | Fixed-weight grid |
| 0.25 | 4 | 9 | 2 | 8 | Fixed-weight grid |
| 0.30 | 2 | 9 | 2 | 8 | Fixed-weight grid |
| 0.40 | 1 | 9 | 3 | 8 | Fixed-weight grid |
| 0.50 | 0 | 9 | 3 | 8 | M2 equal-weight control |
| 0.75 | 0 | 9 | 4 | 8 | Fixed-weight grid |
| 1.00 | 0 | 10 | 4 | 8 | Fixed-weight grid |
| M1: donor-specific | 1 | 9 | 4 | 8 | Nucleus-ratio control |
| 0.0965 | 7 | 9 | 2 | 8 | Discovery q05 reference |

Post hoc descriptive analysis; no P values. w is the endothelial share in a linear-CPM dermal proxy, not a measured cell or RNA fraction. M1 uses each donor's endothelial/(fibroblast + endothelial) nucleus ratio (range 0.137–0.864, median 0.384). The discovery reference uses the layer=Dermis mean relative abundances from Supporting Data Fig4C (q05): fibroblast 0.5424553, endothelial_strict 0.0579216, giving w = 0.0964754. It is not a composition estimate for the 10 replication sections.

**Supplementary Figure S1. Full discovery-cohort single-channel *PYCARD* maps on a locked cross-sample scale.**

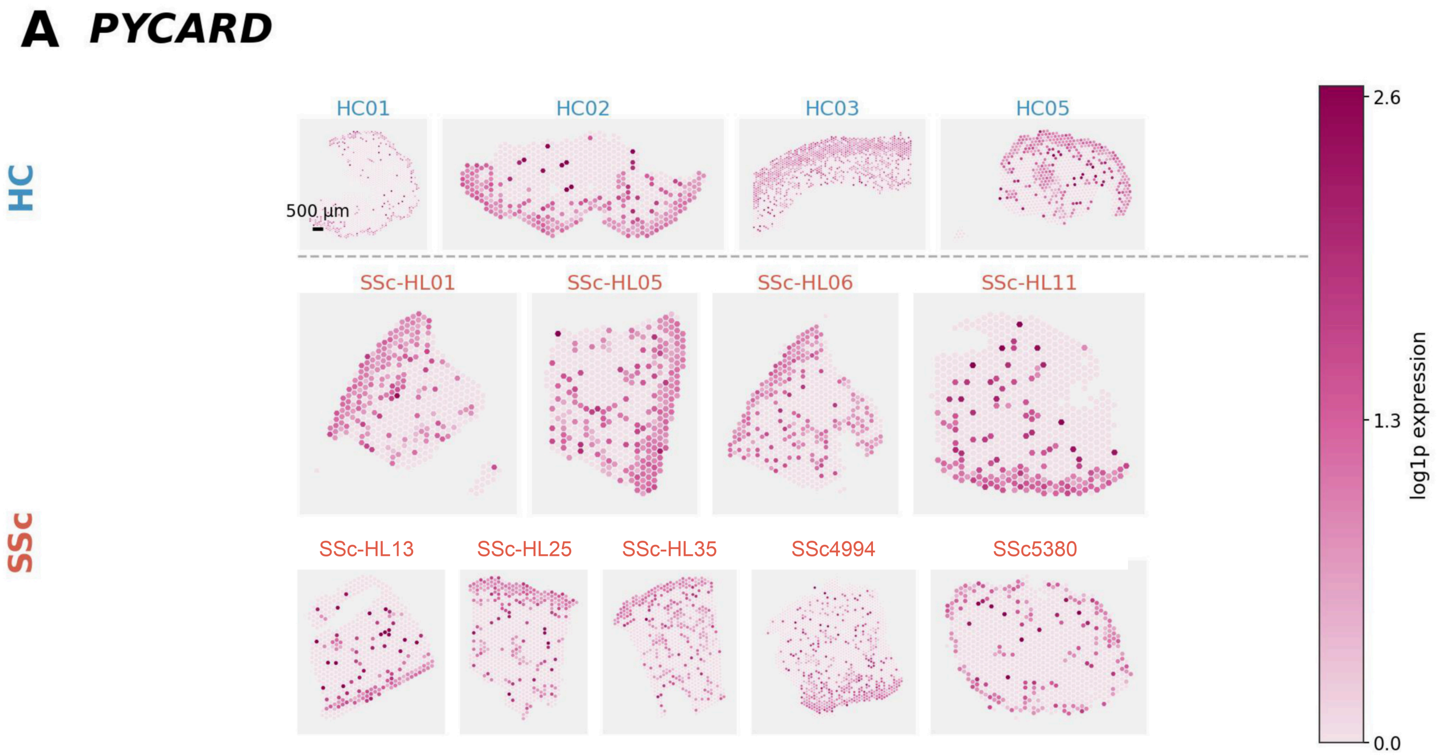


(A) Single-channel hex-spot maps for *PYCARD* across all 13 QC-passing discovery sections (4 HC and 9 SSc), displayed on the same cross-sample intensity scale to facilitate visual comparison of epidermal-versus-dermal localization. Off-tissue or unavailable regions are shown separately from in-tissue low-expression spots.

**Supplementary Figure S2. Full discovery-cohort single-channel *GSDMD* maps on a locked cross-sample scale.**

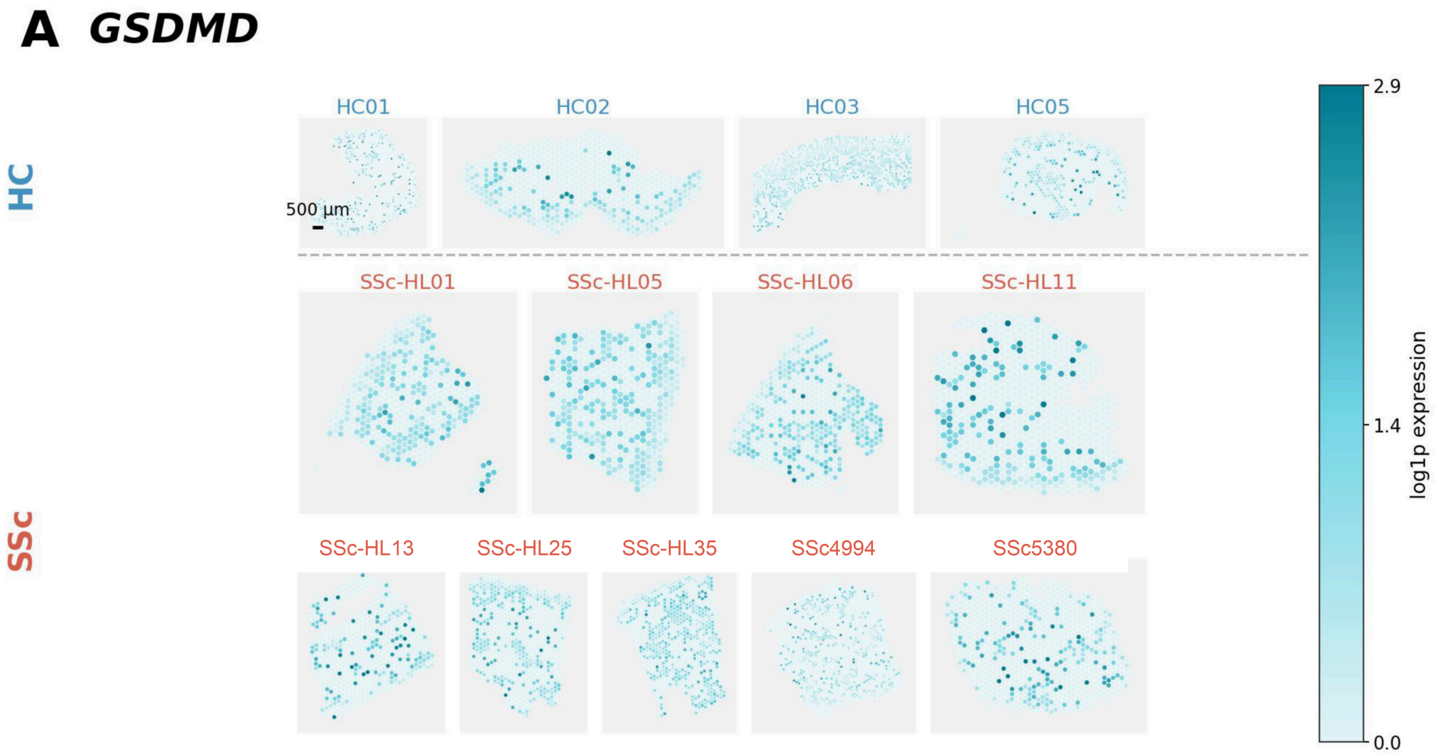


(A) Single-channel hex-spot maps for *GSDMD* across all 13 QC-passing discovery sections (4 HC and 9 SSc), displayed on the same cross-sample intensity scale to facilitate visual comparison of epidermal-versus-dermal localization. Off-tissue or unavailable regions are shown separately from in-tissue low-expression spots.

#### Supplementary Figure S3. Exploratory discovery-gene layer-bias heatmap.

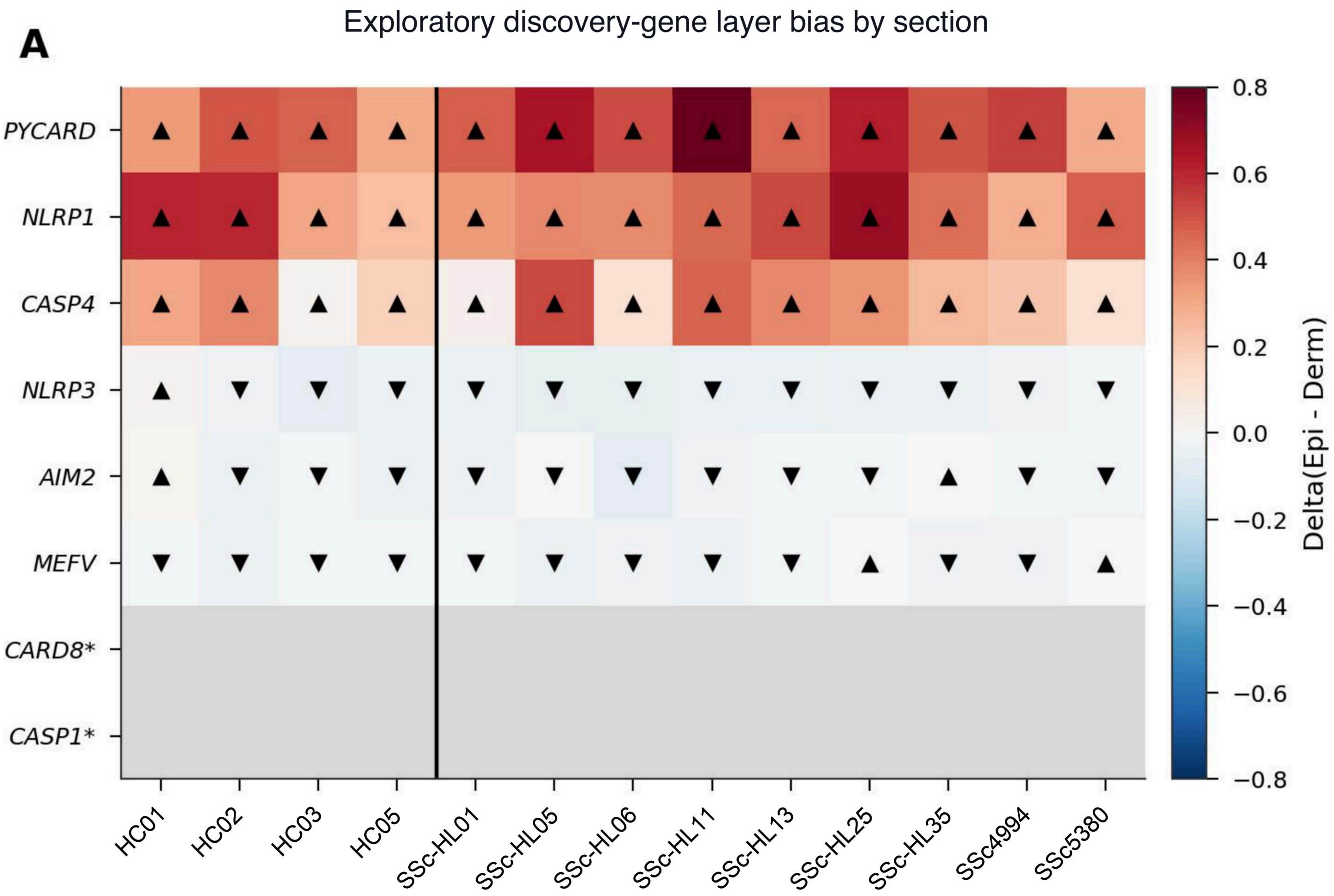


* CARD8 and CASP1 were excluded during probe filtering and were absent from the filtered feature space; shown as NA.

(A) Heatmap of section-level Δ(Epidermis − Dermis) across 13 discovery sections for eight screened inflammasome-related genes. *CARD8* and *CASP1* had been excluded during probe filtering and were absent from the filtered FFPE feature space; they are shown as explicit missingness (NA), not biological zero. The screen and displayed direction summaries are exploratory and post-selection.

**Supplementary Figure S4. Detection context and section-level layer bias for *NLRP3* in the discovery cohort.**

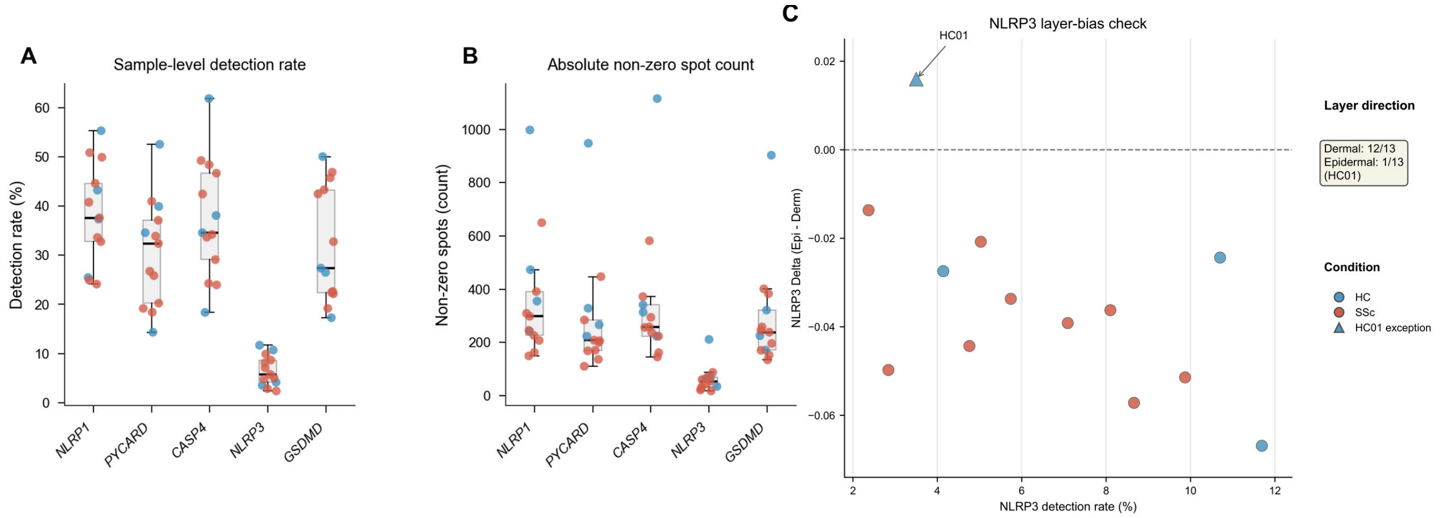


(A) Section-level detection rate (%) across 13 discovery sections for *NLRP1*, *PYCARD*, *CASP4*, *NLRP3*, and *GSDMD*. (B) Absolute non-zero spot counts for the same genes and sections. (C) *NLRP3* detection rate versus section-level Δ(Epidermis − Dermis), showing dermal direction in 12 of 13 sections and HC01 as the sole epidermal exception. Panels A and B provide detection context relative to the direction-concordant epidermal triad and *GSDMD*; panel C summarizes the opposing *NLRP3* layer-bias pattern.

**Supplementary Figure S5. Majority-direction concordance in the exploratory discovery screen.**

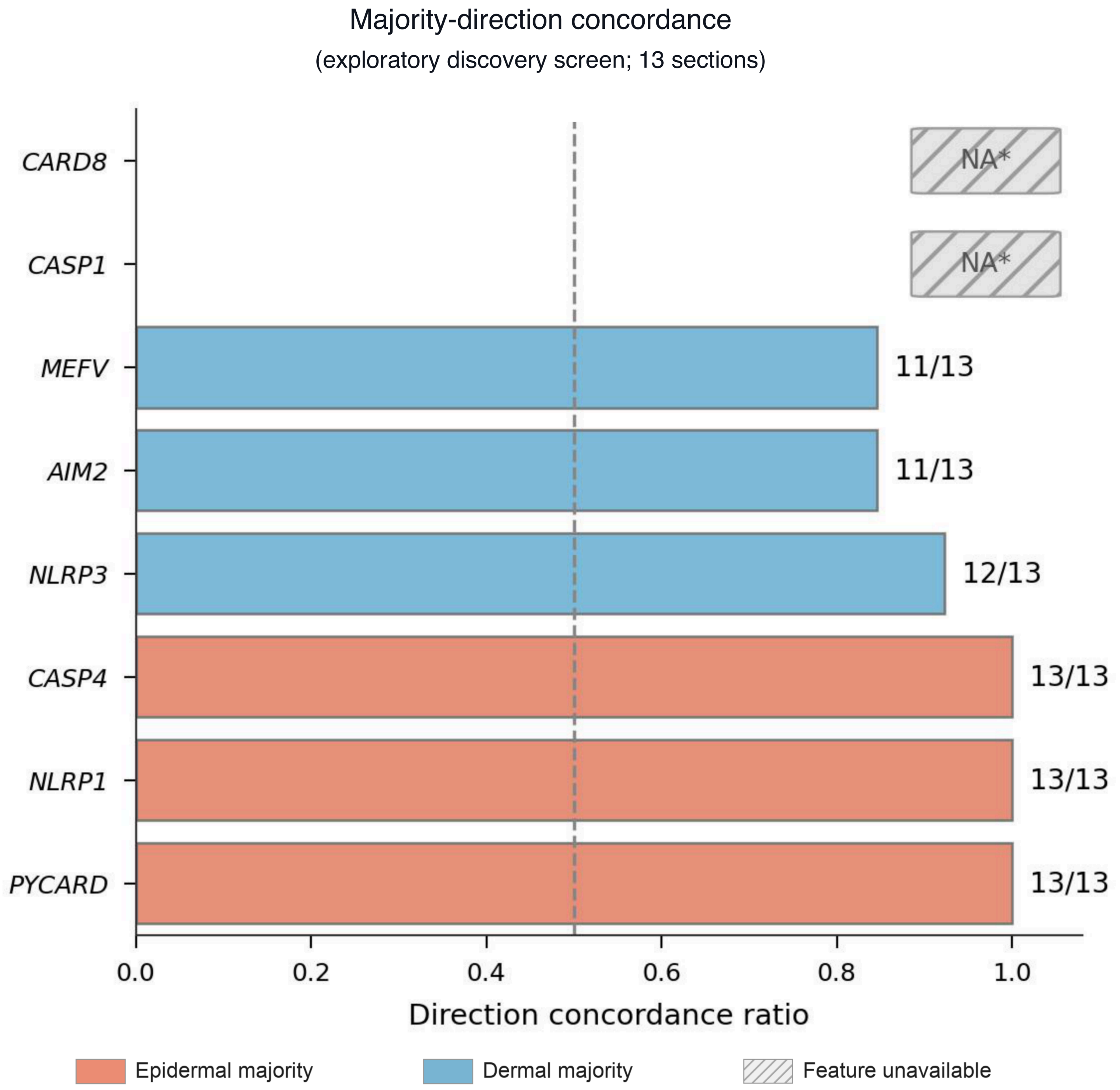


* CARD8 and CASP1 were excluded during probe filtering and were absent from the filtered feature space; shown as NA.

Concordance was defined as max(n_epidermal, n_dermal) / n_evaluable, with the majority direction shown by colour. *NLRP3* was dermal in 12/13 sections, *AIM2* and *MEFV* were dermal in 11/13, and *PYCARD*, *NLRP1*, and *CASP4* were epidermal in 13/13. This differs from Figure 2A, which reports the signed epidermal fraction n[Δ(Epidermis − Dermis) > 0] / n_evaluable. *CARD8* and *CASP1* had been excluded during probe filtering and were absent from the filtered feature space; they are shown as NA.

#### Supplementary Figure S6. Discovery-cohort module-score summaries.

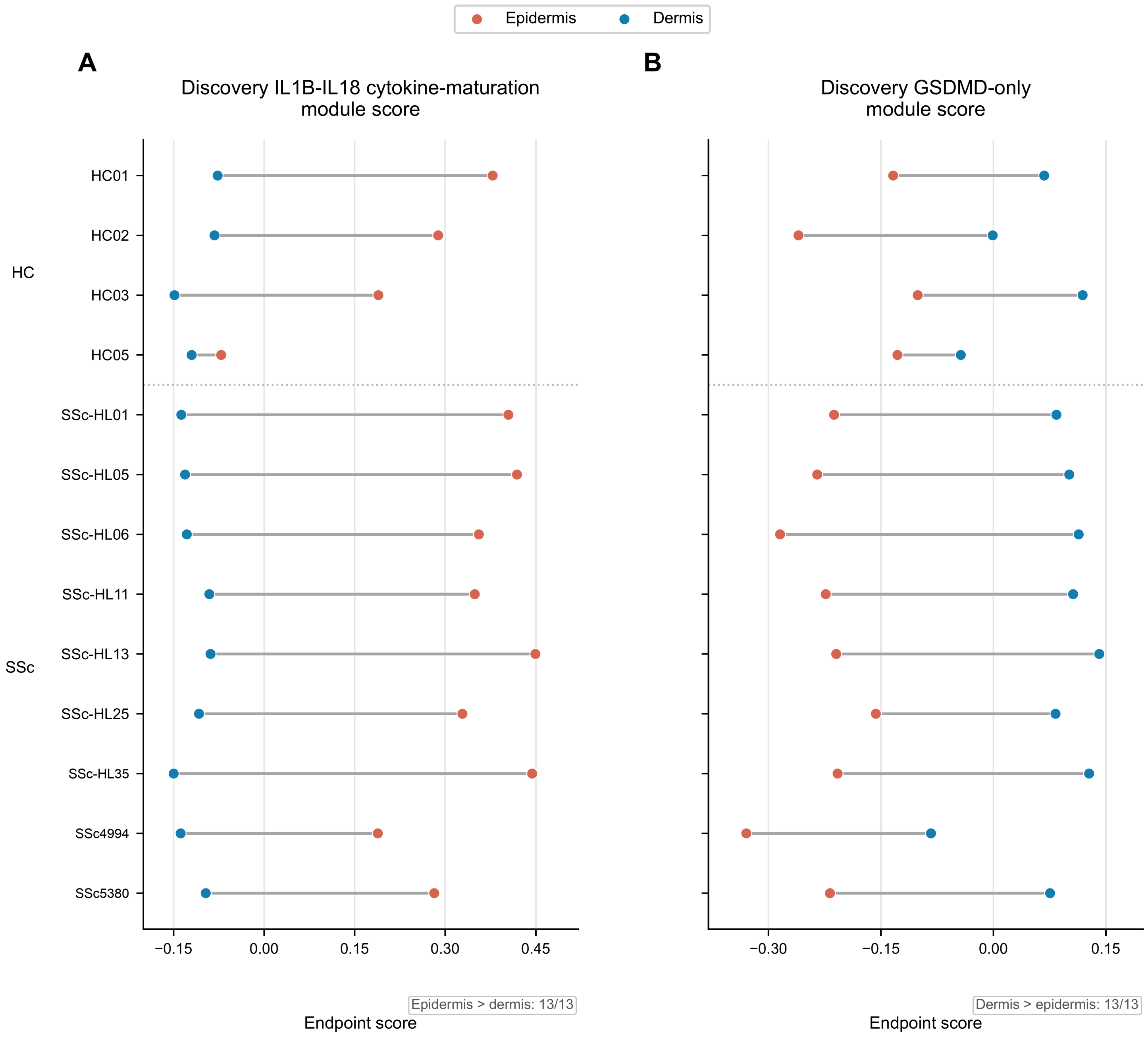


(A) Section-level epidermal and dermal values for the Scanpy *IL1B–IL18* cytokine-maturation module score (pyro_B_maturation_score), summarized by compartment median. (B) Corresponding values for the Scanpy *GSDMD*-only module score (pyro_gasdermin_score), summarized by compartment mean. Both axes show endpoint-score units. Panel A was epidermally directed in 13/13 sections and panel B was dermally directed in 13/13. These discovery scores are not the Whitfield five-gene and direct-*GSDMD* replication endpoints.

**Supplementary Figure S7. Descriptive comparison of pyroptosis-related and compact comparator cell-death modules.**

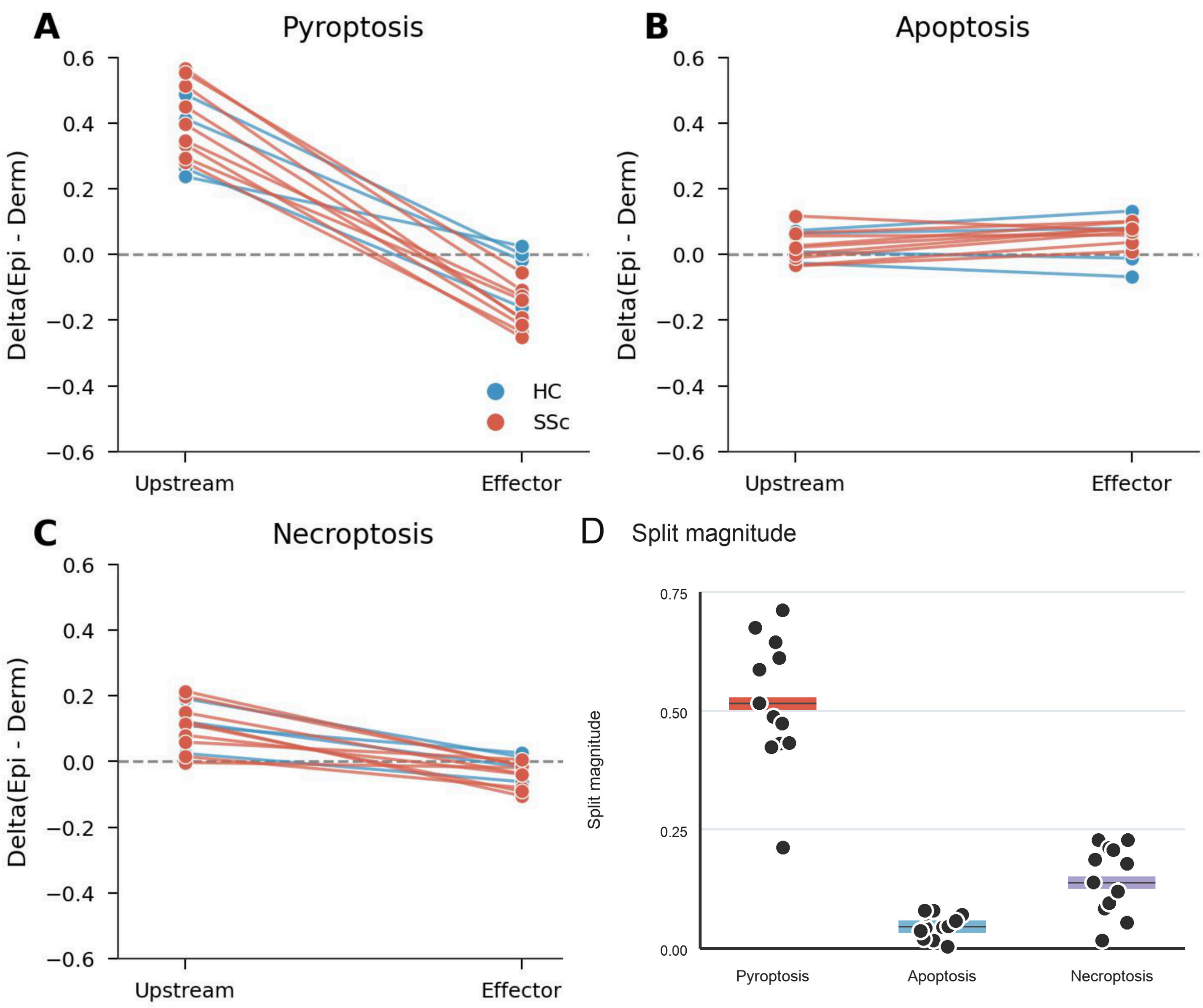


(A) Section-level layer-bias summary for the pyroptosis-related module. (B) Matching section-level summary for the apoptosis comparator module. (C) Matching section-level summary for the necroptosis comparator module. (D) Split magnitudes across the three module families; points show individual sections and horizontal lines show medians (n = 13 sections per family). The same section-level layer-bias framework was applied across families. The comparison is descriptive, the component sets differ in size and composition, and no additional statistical test was introduced for this display.

**Supplementary Figure S8. Independent-cohort comparison of prespecified related endpoints and targeted post hoc evaluation of the discovery-defined signed triad in the Whitfield/Jarnagin dataset.**

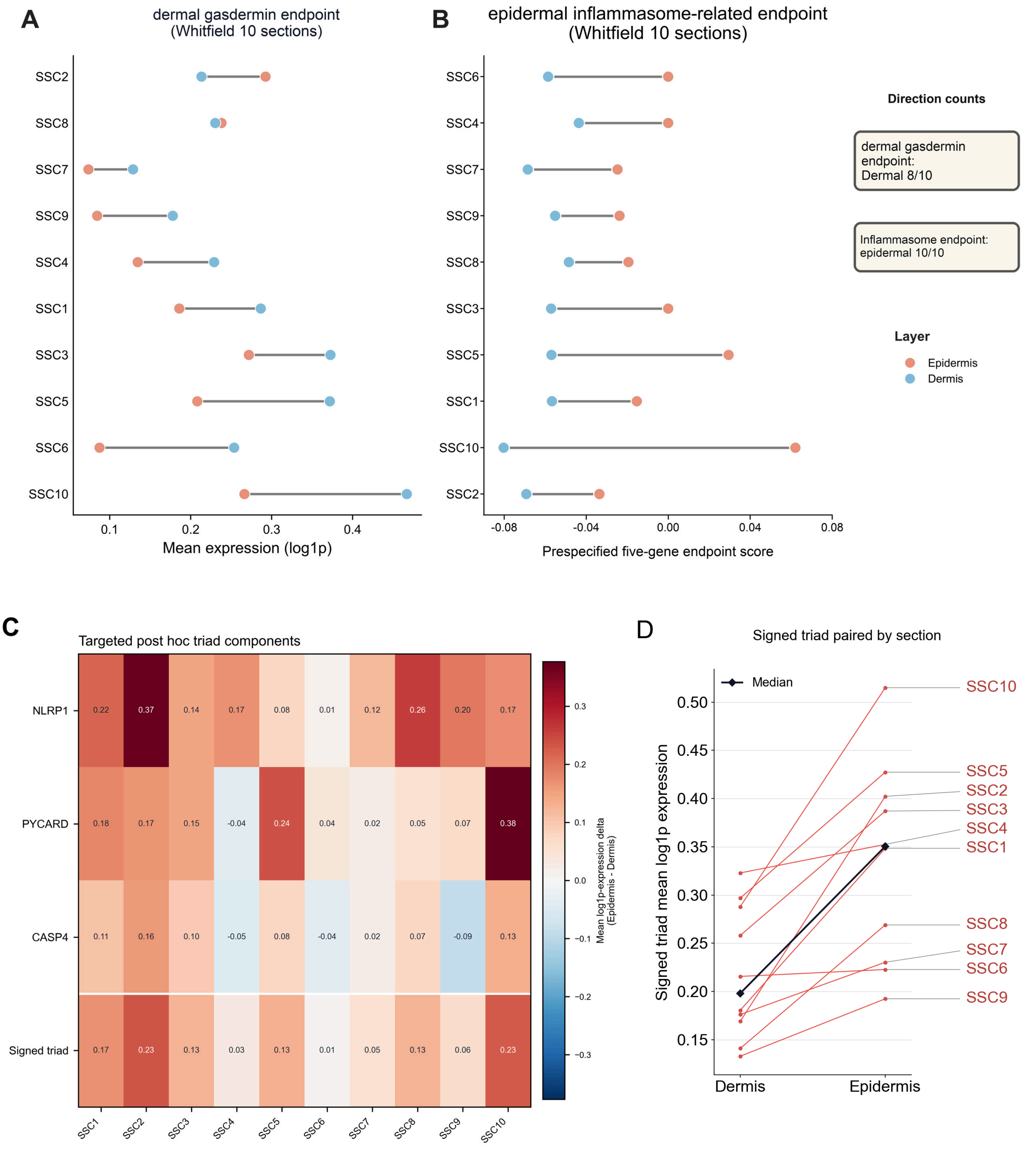


| | SSC1 | SSC2 | SSC3 | SSC4 | SSC5 | SSC6 | SSC7 | SSC8 | SSC9 | SSC10 |
|---|---|---|---|---|---|---|---|---|---|---|
| NLRP1 | 0.22 | 0.37 | 0.14 | 0.17 | 0.08 | 0.01 | 0.12 | 0.26 | 0.20 | 0.17 |
| PYCARD | 0.18 | 0.17 | 0.15 | -0.04 | 0.24 | 0.04 | 0.02 | 0.05 | 0.07 | 0.38 |
| CASP4 | 0.11 | 0.16 | 0.10 | -0.05 | 0.08 | -0.04 | 0.02 | 0.07 | -0.09 | 0.13 |
| Signed triad | 0.17 | 0.23 | 0.13 | 0.03 | 0.13 | 0.01 | 0.05 | 0.13 | 0.06 | 0.23 |

Epidermal direction: NLRP1 10/10; PYCARD 9/10; CASP4 7/10; signed triad 10/10. Section is the inferential unit.

#### Supplementary Figure S8 (continued).

(A) Paired epidermal and dermal mean expression (log1p) for the dermal gasdermin endpoint across the 10 Whitfield systemic sclerosis sections, with dermal-majority alignment in 8 of 10 sections. (B) Paired epidermal and dermal values of the prespecified five-gene endpoint (*PYCARD*, *NLRP3*, *CASP1*, *IL1B*, *IL18*), summarized by compartment median across the same 10 sections, with epidermal-majority alignment in 10 of 10 sections. This score-based endpoint remained analytically distinct from the direct gene-mean discovery triad. (C) Section-level heatmap of mean log1p-normalized expression differences defined as Epidermis − Dermis for *NLRP1*, *PYCARD*, *CASP4*, and the fixed signed triad; the triad value is the sectionwise mean of the three fixed gene contrasts. (D) Paired dermal and epidermal signed-triad summaries for the same sections; each line represents one section and the black line and diamonds show cohort medians. In this targeted post hoc analysis, *NLRP1*, *PYCARD*, and *CASP4* showed epidermal direction in 10 of 10, 9 of 10, and 7 of 10 sections, respectively, whereas the fixed signed triad was epidermally directed in 10 of 10 sections. The triad median delta was 0.1285 (IQR 0.0555–0.1586; range 0.0072–0.2333). A two-sided one-sample Wilcoxon signed-rank test of the 10 section-level triad deltas against zero gave P = 0.001953 and is reported as exploratory and secondary. The tissue section was the inferential unit; no pooled spot-level inference was performed. These mRNA-level results do not establish protein activation or functional pyroptosis.

**Supplementary Figure S9. Ranking of marker-defined dermal tissue-context scores by association with *GSDMD*.**

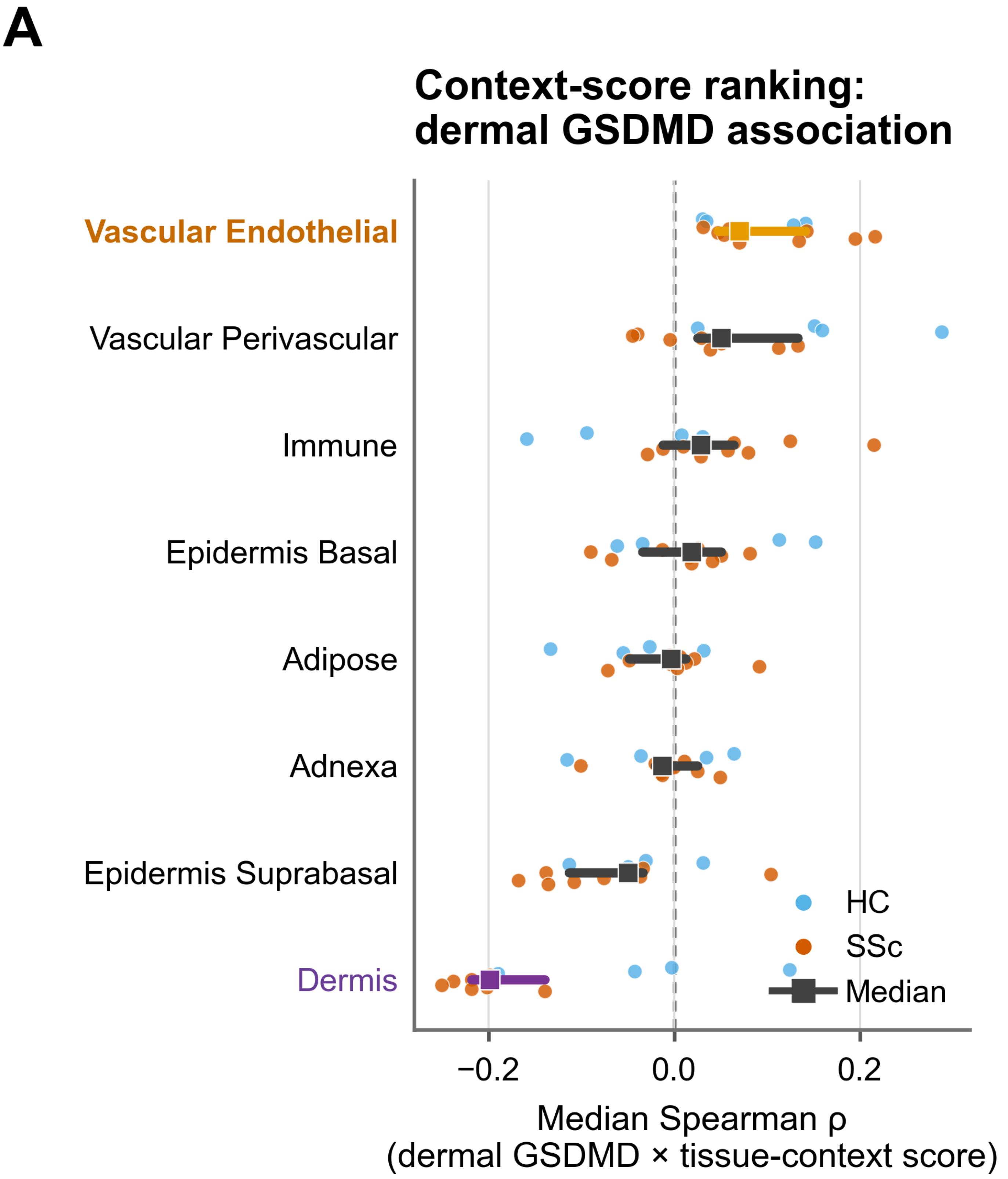


(A) Section-level ranking of Spearman associations between dermal *GSDMD* and marker-defined tissue-context scores within QC-passing dermal spots in the discovery cohort. Context scores are estimated spatial summaries and are interpreted as tissue-context associations rather than direct cell counts or cell-of-origin assignments. Circles show individual discovery sections (HC, n=4; SSc, n=9); squares show the across-section median, and thick horizontal segments span the 25th to 75th percentiles (interquartile range). These segments are not confidence intervals.

**Supplementary Figure S10. Ancillary source-score adjustment for the dermal IFNγ-response–*GSDMD* association.**

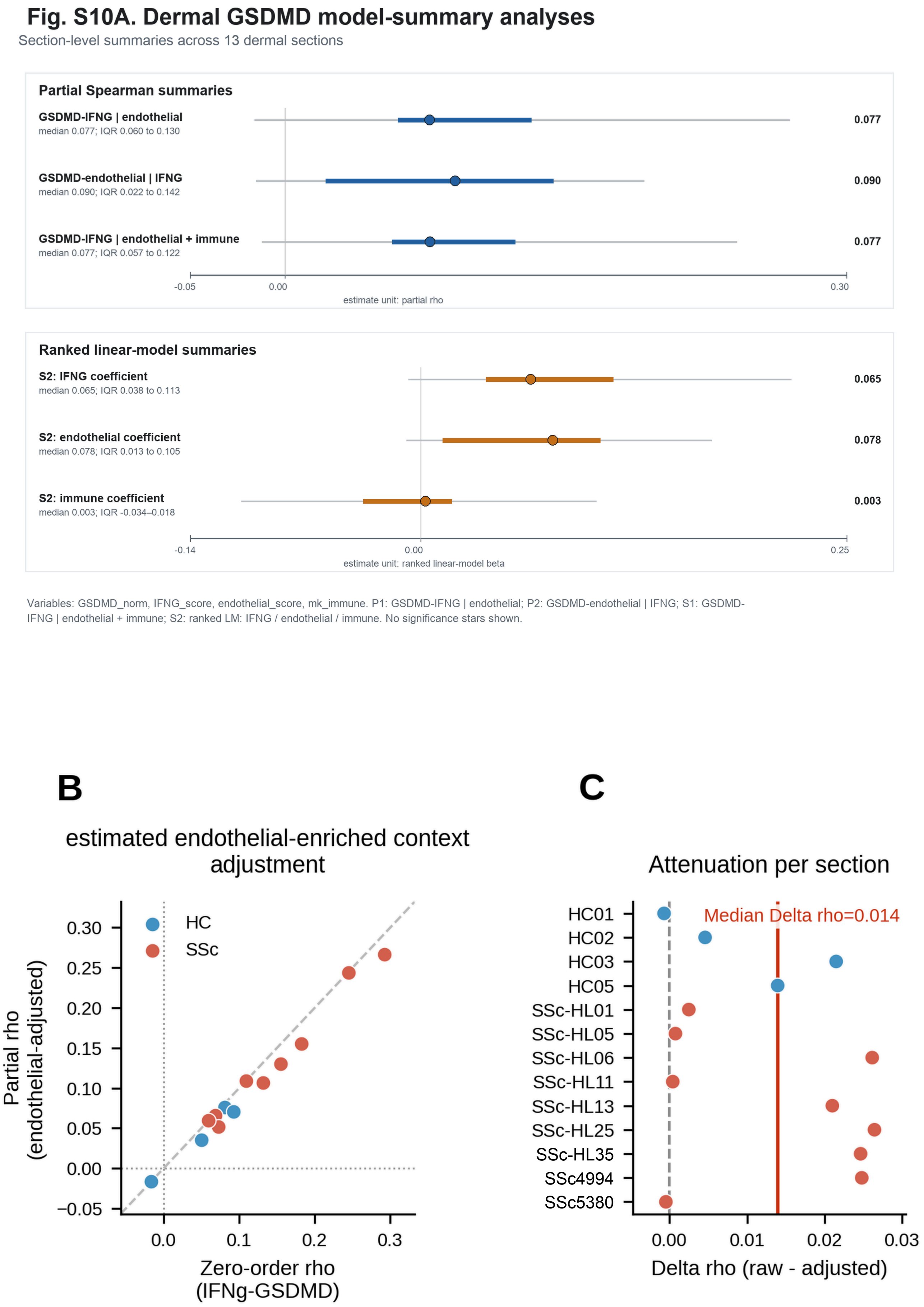

#### Supplementary Figure S10 (continued).

(A) Model-summary view for GSDMD_norm, IFNG_score, endothelial_score, and mk_immune. P1 shows rho(GSDMD_norm, IFNG_score | endothelial_score), P2 shows rho(GSDMD_norm, endothelial_score | IFNG_score), S1 adds the implemented 12-gene immune-score sensitivity covariate mk_immune, and S2 summarizes ranked linear-model coefficients. The frozen analysis implemented the planned Fisher-z P1/P2 family. At reporting, the two-sided one-sample Wilcoxon test across 13 P1 section-level partial rho values was designated primary, while planned pooled Fisher-z summaries were retained as secondary. (B) Raw-versus-adjusted section-level correlations. (C) Sectionwise attenuation, with median $\Delta\rho$ [raw − adjusted] = +0.014. Values remain section-level quantities rather than pooled spot-level inference. In panel A, filled circles show the across-section median, thick colored segments show the interquartile range, and thin gray segments span the minimum to maximum section-level values. The upper block shows partial Spearman coefficients, whereas the lower block shows ranked linear-model coefficients; neither set of segments is a confidence interval. In the retained model labels, endothelial refers to the precomputed endothelial_score column, whose original construction could not be verified (Supplementary Methods Section 11); these ancillary results are not evidence of adjustment for endothelial cell abundance. The separate post hoc adjustment using Cell2location-estimated endothelial relative abundance is reported in Supplementary Table S8 and is not depicted in this figure.

**Supplementary Figure S11. Ancillary source-score adjustment of the interferon-gamma–*GSDMD* association and integrative translayer architecture.**

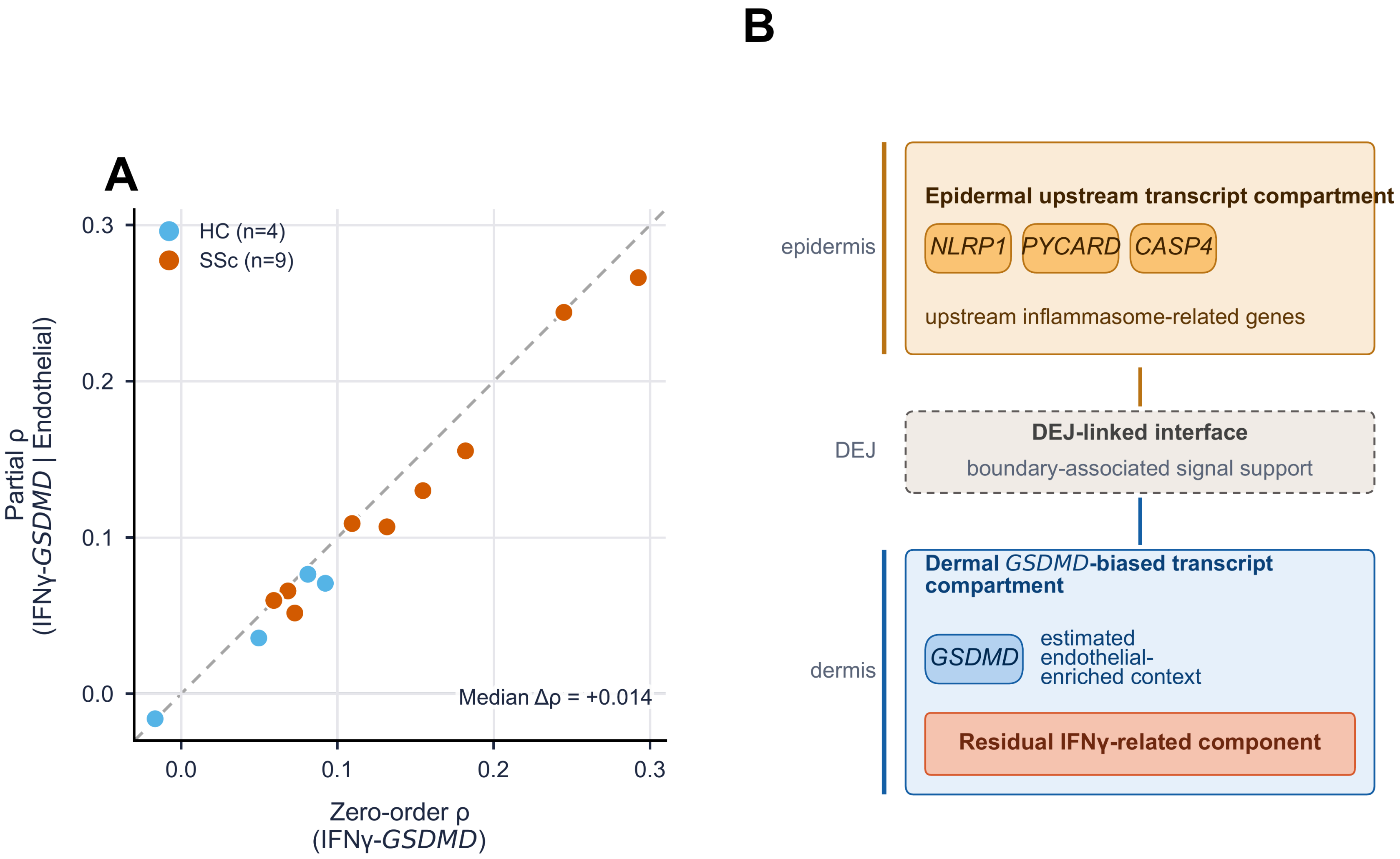

#### Supplementary Figure S11 (continued).

(A) Zero-order versus endothelial_score-adjusted Spearman rho for the dermal interferon-gamma–*GSDMD* association across the 13 discovery sections (HC, n=4; SSc, n=9). The diagonal is the identity line; proximity to the diagonal indicates modest attenuation after adjustment (median sectionwise partial rho=0.077; reporting-stage primary two-sided one-sample Wilcoxon signed-rank P=0.000488; median delta rho [raw − adjusted]=+0.014). The planned pooled Fisher-z summary of the adjusted partial rho (0.094 [95% CI 0.073–0.114]) is retained as a secondary summary aggregating sections. Supplementary Fig. S10 provides model-summary and sectionwise sensitivity details. Adjusted denotes use of the precomputed endothelial_score, whose original construction could not be verified; it does not establish adjustment for endothelial cell abundance. (B) Hypothesis-generating integrative spatial summary of an upstream transcript compartment biased toward the epidermis, an interface near the dermal–epidermal junction, and a dermal transcript compartment biased toward *GSDMD*. This panel integrates direct spatial observations and computationally inferred contexts and does not assert pathway directionality or mechanism beyond the mRNA-level architecture. IFNG denotes the implemented 13-gene interferon-gamma-response score. The separate post hoc adjustment using Cell2location-estimated endothelial relative abundance is reported in Supplementary Table S8 and is not depicted in this figure.

**Supplementary Figure S12. Related but nonidentical endpoints show directional consistency in an independent cohort.**

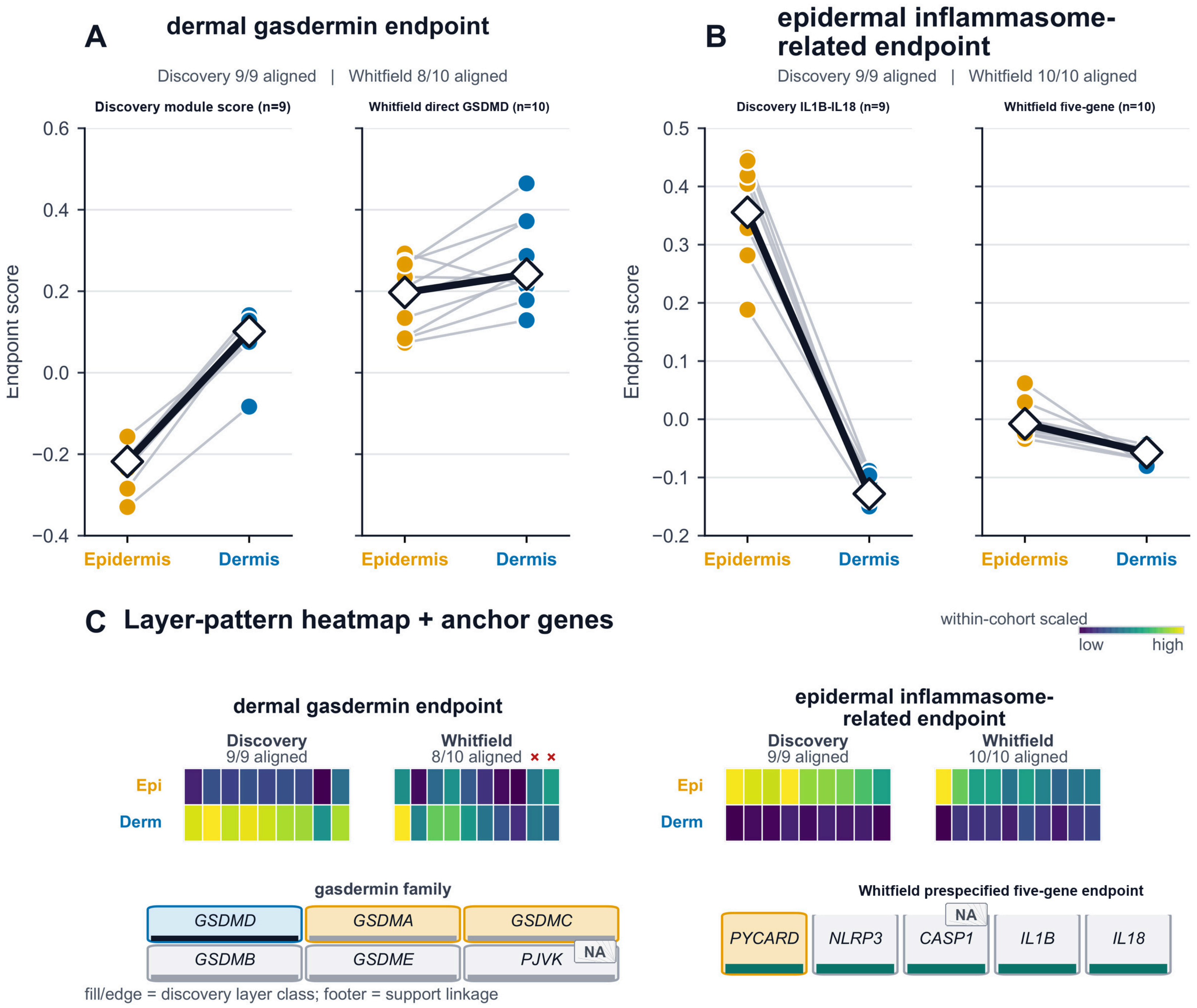

#### Supplementary Figure S12 (continued).

(A) Paired epidermal and dermal values for the discovery *GSDMD*-only module score (compartment means; discovery SSc subset, n=9) and the prespecified Whitfield direct log1p-normalized *GSDMD* endpoint (compartment means; n=10). (B) Paired epidermal and dermal values for the discovery *IL1B*-*IL18* cytokine-maturation module score (compartment medians) and the prespecified Whitfield five-gene *PYCARD*-*NLRP3*-*CASP1*-*IL1B*-*IL18* endpoint (compartment medians). (C) Layer-pattern heatmaps scaled within each cohort with anchor-gene context for the dermal and epidermal endpoints. Discovery alignment counts are 9 of 9 for both discovery module scores; Whitfield alignment counts are 8 of 10 for direct *GSDMD* and 10 of 10 for the five-gene epidermal endpoint. The Whitfield five-gene endpoint was distinct from both the discovery *IL1B*-*IL18* module score and the discovery *NLRP1*-*PYCARD*-*CASP4* triad; these panels provide endpoint-level directional support rather than exact construct replication. Y-axis regimes in panels A and B are locked within endpoint using pooled discovery and Whitfield values, whereas panel C is scaled within cohort to emphasize directional patterning rather than absolute magnitude across cohorts. Red x symbols mark the two Whitfield sections not aligned with the dermal endpoint direction. White diamonds and the thick black line in panels A and B show cohort medians. *CASP1* and *PJVK* are marked by hatched NA tags because they were unavailable for discovery layer classification. Component agreement for the post hoc triad was *NLRP1* 10 of 10, *PYCARD* 9 of 10 and *CASP4* 7 of 10.

#### Supplementary Figure S13. Source-score illustration and deconvolution-derived tissue contexts.

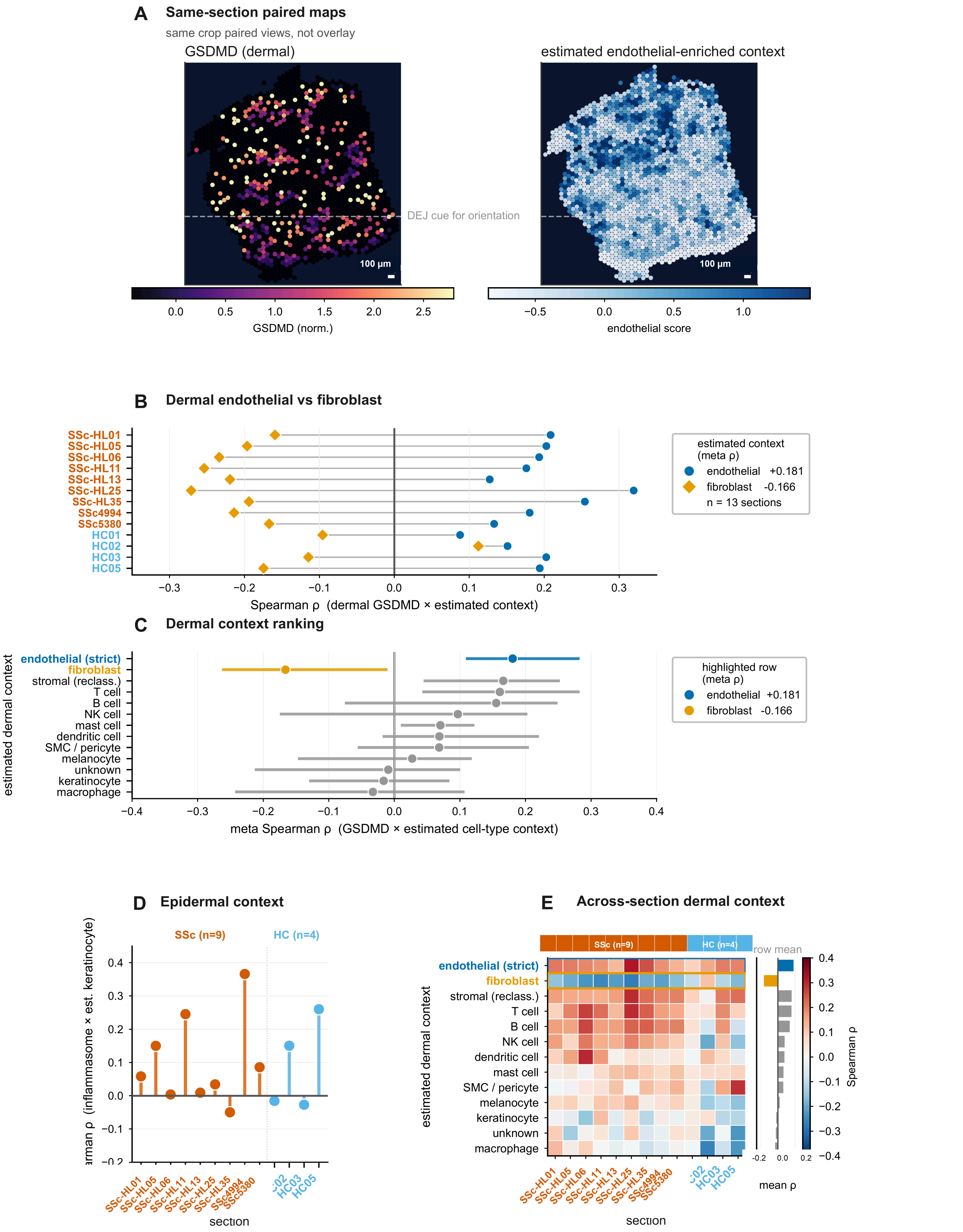

#### Supplementary Figure S13 (continued).

(A) Paired maps from a representative SSc4994 crop show dermal *GSDMD* signal alongside the stored endothelial_score. The original construction of this score could not be verified (Supplementary Methods Section 11); this panel is retained as a source-score illustration, not as a map of cell2location abundance or evidence of cellular origin. (B) Paired sectionwise Spearman rho values compare dermal *GSDMD* with estimated endothelial context and the fibroblast comparator across the 13 discovery sections. (C) Points show secondary fixed-effect Fisher-z pooled estimates of the estimated dermal context associations. Horizontal segments span the 5th to 95th percentiles of the section-level Spearman coefficients, not confidence intervals for the pooled estimates. The ranking does not test whether one context is significantly stronger than another. (D) Each point and stem represents one section's Spearman rho between the epidermal inflammasome score and the estimated keratinocyte context. (E) Heatmap across sections of dermal context associations. For the deconvolution-derived analyses in panels B–E, reference labels were initialized with CellTypist 1.7.1 and Immune_All_Low.pkl using majority voting, followed by the implemented manual consolidation map, marker fallback, and an endothelial_strict override requiring *PECAM1* and *CDH5*; model-file hash unavailable. Deconvolution-derived contexts are inferred quantities and are not direct cell counts or assignments of cellular origin. The frozen P1-P4 family remains a formal result in the main Results.

#### Supplementary Figure S14. Donor-resolved spatial and cell-class context of *NLRP1* in the replication cohort

This figure provides a donor-resolved view of the *NLRP1* results already reported in Figures 3-5 and Supplementary Tables S3–S7; it is not an additional validation cohort. The following two atlas pages show SSc1-5 and SSc6-10, respectively, in the fixed donor order. Spatial maps (left two columns). Each row displays stored expression-derived layer labels and spot-level *NLRP1* mRNA from the same Visium section. All 10,217 retained spots are shown, including zeros and the endothelium-label and other-label spots. Outlined spots are excluded only from the existing epidermis/dermis summary. Expression uses the stored natural-log1p values after normalization to 10,000, with the same linear color range of 0-5.5 throughout. Each donor retains its full coordinate extent; the paired maps share coordinates, extent and spot radius. The labels are not independently adjudicated morphology, and the maps do not establish cellular origin. No histological image, physical scale bar, interpolation or smoothing is used; map dimensions across donors are not common physical lengths. Class-pseudobulk (third column). K, keratinocyte; F, fibroblast; E, endothelial_strict. The 30 frozen values summarize the same 10 donors, using log2(CPM+1) and a library denominator of 18,703 retained features. Nuclear counts are shown for each class; the primary floor is at least 20 nuclei per class. These are cell-class summaries, not per-cell or per-nucleus expression. The low-count classes SSc4 F=22 and SSc7 E=20 remain displayed. E exceeds K in all donors, but E is not always the highest class: SSc7 retains F > E > K. Directions (right column). Observed denotes the frozen Visium epidermis-minus-dermis direction. M1 represents epidermis by keratinocyte expression and dermis by donor-specific fibroblast:endothelial nucleus-ratio weights; M2 uses equal fibroblast:endothelial weights. Class CPM values are mixed on the linear scale before log2(CPM+1) transformation. Both are restricted proxies, not measured Visium composition. Upward orange and downward blue triangles indicate epidermal and dermal directions, respectively. Observed direction is epidermal in all 10 donors. M1 reproduces that direction only in SSc1 (1/10); M2 does so in none (0/10). Observed Visium and class-pseudobulk/reconstruction values use different measurement scales; only directions are compared, without cross-platform residuals or amplitude comparisons. This explanatory display adds no new inferential endpoint. For context, the existing M1/M2 direction-concordance counts are *PYCARD* 9/10 and 9/10, *GSDMD* 8/10 and 8/10, and *CASP4* 4/10 and 3/10. These genes are descriptive comparators. Omitted cell classes, differential nuclear recovery and class-dependent platform capture remain alternative explanations for non-reproduction; the display does not resolve them. The displayed epidermal and dermal means are the 20 stored *NLRP1* layer means transcribed from the frozen section-value table and rounded to three decimal places; they were not recomputed for this display.

#### Supplementary Figure S14 (continued). SSc1–SSc5

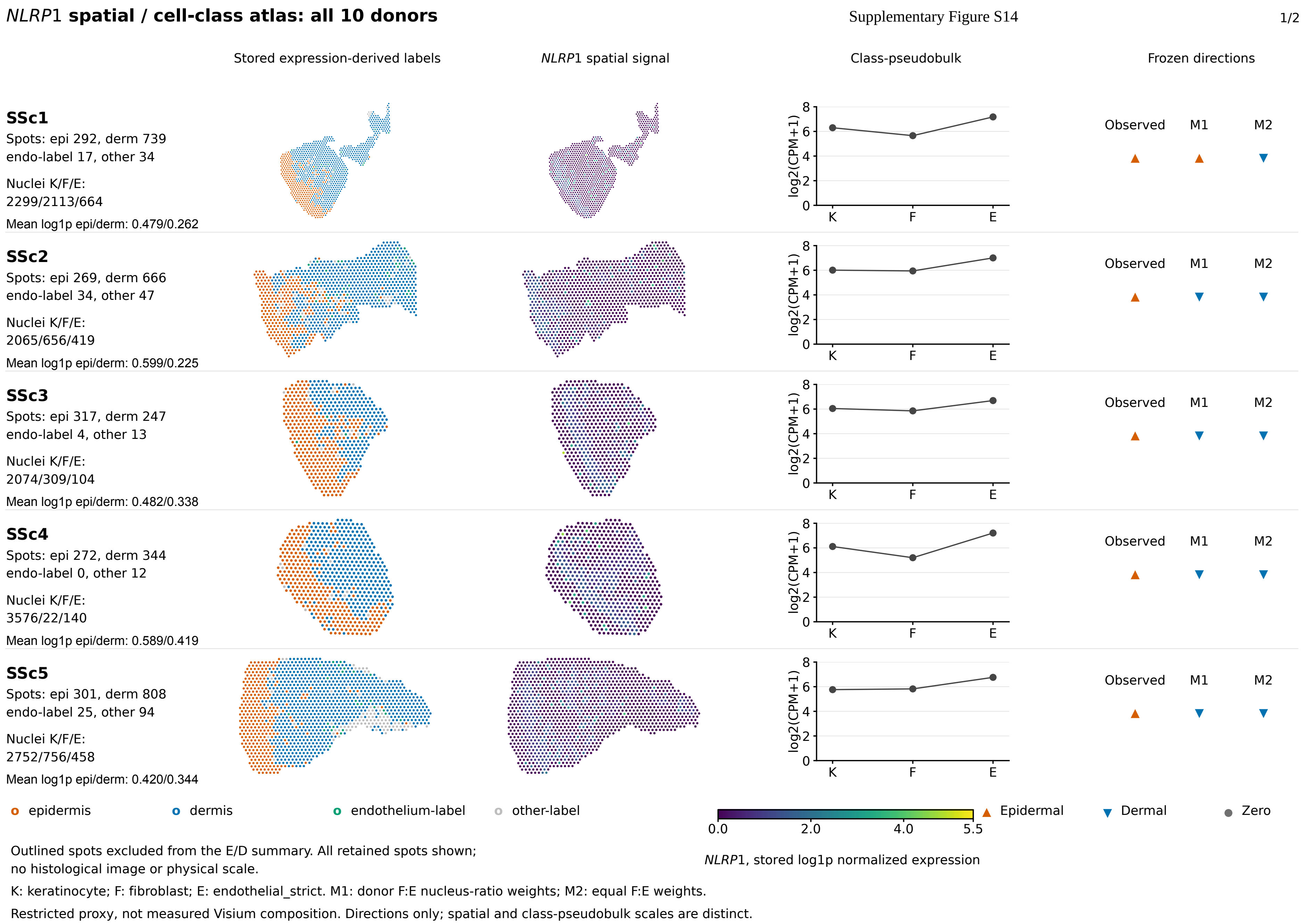

### Supplementary Figure S14 (continued). SSc6–SSc10



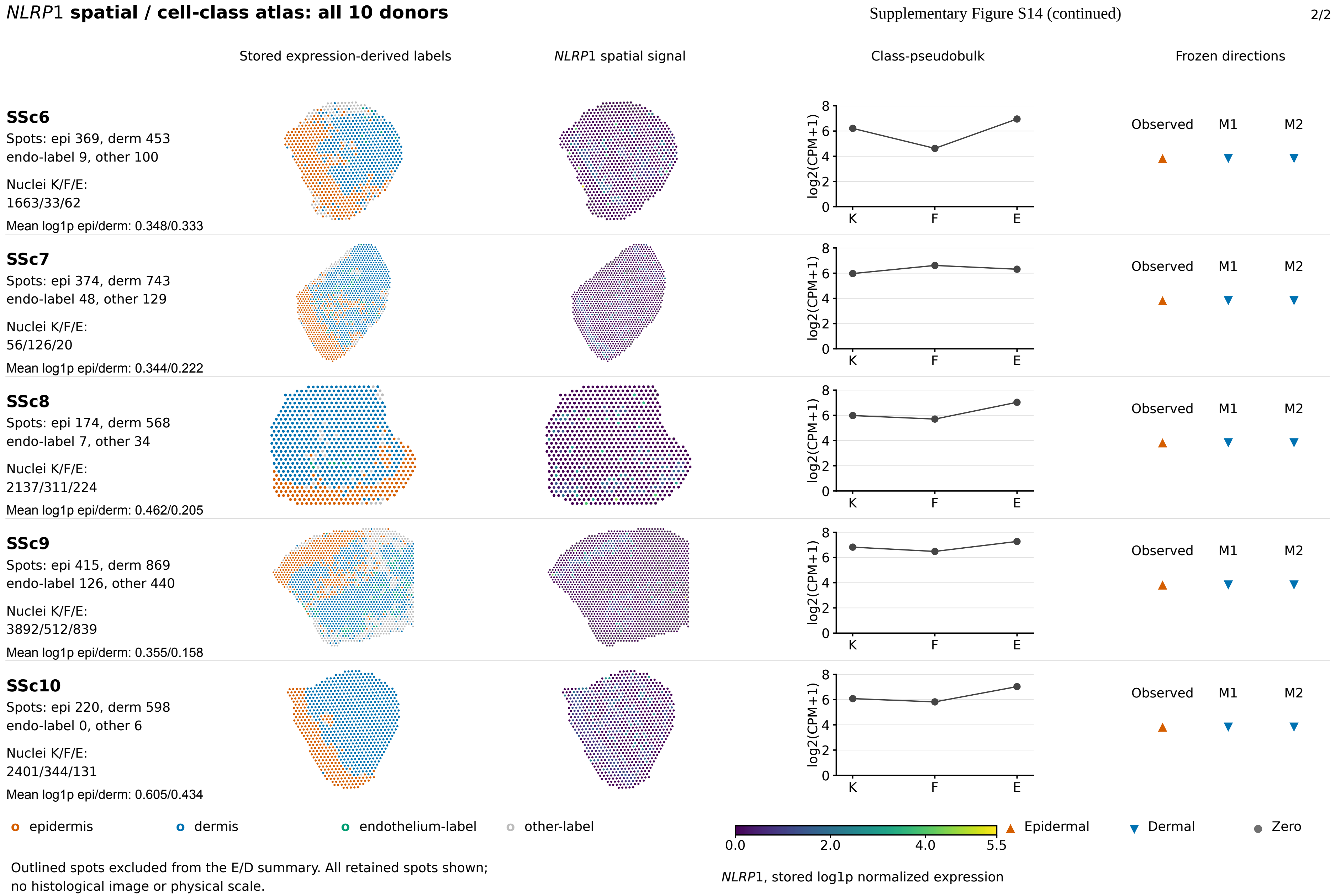


Outlined spots excluded from the E/D summary. All retained spots shown; no histological image or physical scale.

K: keratinocyte; F: fibroblast; E: endothelial_strict. M1: donor F:E nucleus-ratio weights; M2: equal F:E weights.

Restricted proxy, not measured Visium composition. Directions only; spatial and class-pseudobulk scales are distinct.
