## Supplementary Software 1 for "Layer-to-cell-class correspondence is gene-specific for pyroptosis-related transcription in human skin": render.py

```
import argparse, importlib.util
from pathlib import Path
import pandas as pd
from matplotlib import font_manager

p=argparse.ArgumentParser()
p.add_argument('--output',type=Path,required=True)
p.add_argument('--font',type=Path,required=True)
a=p.parse_args()
font_manager.fontManager.addfont(str(a.font))
source=Path(__file__).parent/'scripts/figures/build_figures_3_to_5.py'
spec=importlib.util.spec_from_file_location('frozen_figure5',source)
m=importlib.util.module_from_spec(spec);spec.loader.exec_module(m)
m.OUT=a.output;m.WORK=a.output/'source_formats';m.QA=a.output/'metadata'
for d in [m.OUT,m.WORK,m.QA]:d.mkdir(parents=True,exist_ok=True)
m.require_inputs();m.configure_style()
tables=[pd.read_csv(m.FILES[k][0],sep='\t') for k in ['pred','summary','loo']]
m.build_figure5(*tables)
```
