## Supplementary Software 1 for "Layer-to-cell-class correspondence is gene-specific for pyroptosis-related transcription in human skin": build_figures_3_to_5.py

```
#!/usr/bin/env python3
from __future__ import annotations

import hashlib
import json
import platform
import re
from pathlib import Path

import matplotlib as mpl
import matplotlib.pyplot as plt
from matplotlib import font_manager, ft2font
from matplotlib.lines import Line2D
from matplotlib.patches import FancyBboxPatch
import numpy as np
import pandas as pd
from PIL import Image


RUN = Path(__file__).resolve().parents[2]
SRC = RUN / "intermediate/extended_analysis"
OUT = RUN / "generated/figures_3_to_5"
WORK = OUT / "source_formats"
QA = OUT / "metadata"
GENES = ["NLRP1", "PYCARD", "CASP4", "GSDMD"]
DONORS = [f"SSc{i}" for i in range(1, 11)]

FIGURE_CONTRACT = {
    3: {"figsize_inches": [7.2, 7.15], "dpi": 300, "bbox_inches": "tight"},
    4: {"figsize_inches": [7.2, 6.75], "dpi": 300, "bbox_inches": "tight"},
    5: {"figsize_inches": [7.2, 8.15], "dpi": 300, "bbox_inches": "tight"},
}

FILES = {
    "vis_summary": (SRC / "VISIUM_CORE_SUMMARY.tsv", "65a9e01f74f503a1f53372d43f77759a0442500a730eec3ebbfc00c4818d444f"),
    "vis_sections": (SRC / "VISIUM_CORE_SECTION_VALUES.tsv", "0ea6e6d1a5fc7ef548f547735267cbb25d46a557884e11e4a0c050d78ec4f236"),
    "pseudo": (SRC / "MULTIOME_PSEUDOBULK_VALUES.tsv", "6b95ea8f125c146ab736c7df2925b42fc03f2810f7ab7e015efa5620abf79bac"),
    "contrasts": (SRC / "MULTIOME_CLASS_CONTRASTS.tsv", "11c3fd662c975ffa357057d74573b58259e214cbd5058acf255dd1ae7aac5c59"),
    "component": (SRC / "PART_A_RUN1_COMPONENT_MATRIX.tsv", "447e724e36c52adf9131670e406899e0bd41bedbe516f97c6ae5f954481038e8"),
    "pred": (SRC / "MIXTURE_PREDICTIONS.tsv", "17425a1e19fc86fe30e6f07de45bb3ed2f87900c2069eedc92ec0a7d7fe74a97"),
    "summary": (SRC / "SIGN_CONCORDANCE_SUMMARY.tsv", "bfb15a47aa4ac9219601a4dbf0f8a7a1355ce7861ba05a5331f7f275d90fbb6d"),
    "loo": (SRC / "LOO_DIRECTION_SENSITIVITY.tsv", "8f1631c0f60757926c307d4e82dce28da5a8f476faa493def60e025767fd1c01"),
}

EPI = "#C55A2D"
DERM = "#2F6F9F"
NEUTRAL = "#6F6F6F"
M1 = "#3B6FB6"
M2 = "#E38B2C"
CLASS_COLORS = {
    "keratinocyte": "#D65F2E",
    "fibroblast": "#2C8C74",
    "endothelial_strict": "#6A67A5",
}


def sha256(path: Path) -> str:
    h = hashlib.sha256()
    with path.open("rb") as handle:
        for block in iter(lambda: handle.read(1024 * 1024), b""):
            h.update(block)
    return h.hexdigest()


def pixel_record(path: Path) -> dict:
    with Image.open(path) as image:
        rgb = image.convert("RGB")
        return {
            "file_sha256": sha256(path),
            "pixel_sha256_rgb": hashlib.sha256(rgb.tobytes()).hexdigest(),
            "pixel_dimensions": list(rgb.size),
            "pixel_mode_for_hash": "RGB",
        }


def renderer_record() -> dict:
    preference = ["Arial", "Helvetica", "DejaVu Sans"]
    properties = font_manager.FontProperties(
        family=preference,
        style="normal",
        weight="normal",
    )
    font_path = Path(font_manager.findfont(properties, fallback_to_default=True))
    selected = font_manager.FontProperties(fname=font_path)
    return {
        "operating_system": platform.system(),
        "operating_system_release": platform.release(),
        "architecture": platform.machine(),
        "python": platform.python_version(),
        "python_implementation": platform.python_implementation(),
        "matplotlib": mpl.__version__,
        "pillow": Image.__version__,
        "numpy": np.__version__,
        "freetype": ft2font.__freetype_version__,
        "font_preference": preference,
        "selected_font_family": selected.get_name(),
        "selected_font_file_basename": font_path.name,
        "selected_font_file_sha256": sha256(font_path),
        "selected_font_style": "normal",
        "selected_font_weight": "normal",
        "absolute_font_path_recorded": False,
    }


def require_inputs() -> None:
    for path, expected in FILES.values():
        actual = sha256(path)
        if actual != expected:
            raise RuntimeError(f"input SHA mismatch: {path}: {actual} != {expected}")


def require_absent(path: Path) -> None:
    if path.exists():
        raise RuntimeError(f"output collision: {path}")


def configure_style() -> None:
    mpl.rcParams.update(
        {
            "font.family": "sans-serif",
            "font.sans-serif": ["Arial", "Helvetica", "DejaVu Sans"],
            "font.size": 8,
            "axes.titlesize": 9,
            "axes.labelsize": 8,
            "xtick.labelsize": 7,
            "ytick.labelsize": 7,
            "legend.fontsize": 7,
            "axes.linewidth": 0.7,
            "pdf.fonttype": 42,
            "ps.fonttype": 42,
            "svg.fonttype": "none",
        }
    )


def donor_from_visium(sample: str) -> str:
    match = re.search(r"_(SSC\d+)_visium$", sample)
    if not match:
        raise RuntimeError(f"unexpected Visium sample id: {sample}")
    return match.group(1).replace("SSC", "SSc")


def donor_from_multiome(sample: str) -> str:
    match = re.search(r"_(SSC\d+)_multiome$", sample)
    if not match:
        raise RuntimeError(f"unexpected multiome sample id: {sample}")
    return match.group(1).replace("SSC", "SSc")


def save_figure(fig: plt.Figure, number: int) -> list[Path]:
    base = WORK / f"Figure_{number}"
    png = base.with_suffix(".png")
    pdf = base.with_suffix(".pdf")
    svg = base.with_suffix(".svg")
    tif = OUT / f"Figure_{number}.tif"
    for path in (png, pdf, svg, tif):
        require_absent(path)
    fig.savefig(pdf, facecolor="white", bbox_inches="tight")
    fig.savefig(svg, facecolor="white", bbox_inches="tight")
    fig.savefig(png, dpi=300, facecolor="white", bbox_inches="tight")
    plt.close(fig)
    with Image.open(png) as image:
        image.save(tif, compression="tiff_lzw", dpi=(300, 300))
    return [tif, png, pdf, svg]


def build_figure3(vis_sections: pd.DataFrame, vis_summary: pd.DataFrame) -> list[Path]:
    values = vis_sections[vis_sections["row_type"] == "section_value"].copy()
    values["donor"] = values["sample_id"].map(donor_from_visium)
    values["raw_E_minus_D"] = values["epidermis_mean_log1p"] - values["dermis_mean_log1p"]
    values["donor_order"] = values["donor"].str.replace("SSc", "", regex=False).astype(int)
    values = values.sort_values(["gene", "donor_order"])
    if len(values) != 40:
        raise RuntimeError(f"expected 40 direct section rows, found {len(values)}")

    fig = plt.figure(figsize=(7.2, 7.15))
    grid = fig.add_gridspec(3, 1, height_ratios=[1.7, 1.25, 1.55], hspace=0.52, left=0.105, right=0.98, top=0.945, bottom=0.08)

    ax_a = fig.add_subplot(grid[0, 0])
    positions = np.arange(len(GENES))
    for x, gene in enumerate(GENES):
        g = values[values["gene"] == gene].sort_values("donor_order")
        jitter = np.linspace(-0.13, 0.13, len(g))
        color = DERM if gene == "GSDMD" else EPI
        ax_a.scatter(np.full(len(g), x) + jitter, g["raw_E_minus_D"], s=28, color=color, edgecolor="white", linewidth=0.5, zorder=3)
        ax_a.plot([x - 0.22, x + 0.22], [g["raw_E_minus_D"].median()] * 2, color="black", lw=2.0, zorder=4)
    ax_a.axhline(0, color="#555555", lw=0.8)
    ax_a.set_xticks(positions, GENES, fontstyle="italic")
    ax_a.set_ylabel("Epidermis - Dermis\nmean log1p expression")
    ax_a.set_title("Section-level direct-gene effects (all 10 sections)", loc="left")
    ax_a.grid(axis="y", color="#E5E5E5", lw=0.55)
    ax_a.spines[["top", "right"]].set_visible(False)
    ax_a.text(-0.11, 1.08, "a", transform=ax_a.transAxes, fontsize=12, fontweight="bold", va="top")

    ax_b = fig.add_subplot(grid[1, 0])
    matrix = np.zeros((len(GENES), len(DONORS)), dtype=int)
    for gi, gene in enumerate(GENES):
        g = values[values["gene"] == gene].set_index("donor")
        matrix[gi] = np.sign([g.loc[donor, "raw_E_minus_D"] for donor in DONORS]).astype(int)
    for gi, gene in enumerate(GENES):
        for di, donor in enumerate(DONORS):
            sign = matrix[gi, di]
            color = EPI if sign > 0 else DERM if sign < 0 else NEUTRAL
            marker = "^" if sign > 0 else "v" if sign < 0 else "o"
            ax_b.scatter(di, len(GENES) - 1 - gi, marker=marker, s=52, color=color, edgecolor="black", linewidth=0.35)
    ax_b.set_xticks(range(10), DONORS, rotation=45, ha="right")
    ax_b.set_yticks(range(3, -1, -1), GENES, fontstyle="italic")
    ax_b.set_xlim(-0.55, 9.55)
    ax_b.set_ylim(-0.55, 3.55)
    ax_b.grid(axis="x", color="#ECECEC", lw=0.5)
    ax_b.spines[["top", "right", "left", "bottom"]].set_visible(False)
    ax_b.tick_params(axis="both", length=0)
    ax_b.set_title("Observed layer direction by section", loc="left")
    ax_b.text(-0.11, 1.12, "b", transform=ax_b.transAxes, fontsize=12, fontweight="bold", va="top")
    ax_b.legend(
        handles=[
            Line2D([0], [0], marker="^", color="none", markerfacecolor=EPI, markeredgecolor="black", markersize=6, label="epidermal"),
            Line2D([0], [0], marker="v", color="none", markerfacecolor=DERM, markeredgecolor="black", markersize=6, label="dermal"),
        ],
        frameon=False,
        ncol=2,
        loc="upper right",
        bbox_to_anchor=(1.0, 1.16),
    )

    ax_c = fig.add_subplot(grid[2, 0])
    discovery_counts = {"NLRP1": 13, "PYCARD": 13, "CASP4": 13, "GSDMD": 13}
    replication_counts = {
        row["gene"]: int(row["positive"] if row["gene"] != "GSDMD" else row["positive"])
        for _, row in vis_summary.iterrows()
    }
    q_values = {row["gene"]: float(row["BH_q_m4"]) for _, row in vis_summary.iterrows()}
    y = np.arange(len(GENES))[::-1]
    ax_c.barh(y + 0.16, [discovery_counts[g] / 13 for g in GENES], height=0.26, color="white", edgecolor="#555555", hatch="///", label="Discovery direction context (of 13)")
    ax_c.barh(y - 0.16, [replication_counts[g] / 10 for g in GENES], height=0.26, color="#4C78A8", label="GSE312129 direct family (of 10)")
    for yi, gene in zip(y, GENES):
        ax_c.text(discovery_counts[gene] / 13 + 0.015, yi + 0.16, f"{discovery_counts[gene]}/13", va="center", fontsize=7)
        ax_c.text(replication_counts[gene] / 10 + 0.015, yi - 0.16, f"{replication_counts[gene]}/10; q={q_values[gene]:.4g}", va="center", fontsize=7)
    ax_c.set_yticks(y, GENES, fontstyle="italic")
    ax_c.set_xlim(0, 1.34)
    ax_c.set_xlabel("Fraction in expected direction")
    ax_c.set_title("Discovery context and fixed direct replication remain distinct families", loc="left")
    ax_c.grid(axis="x", color="#E5E5E5", lw=0.5)
    ax_c.spines[["top", "right", "left"]].set_visible(False)
    ax_c.tick_params(axis="y", length=0)
    ax_c.legend(frameon=False, loc="upper right", bbox_to_anchor=(1.0, 1.16), fontsize=6.8)
    ax_c.text(-0.11, 1.08, "c", transform=ax_c.transAxes, fontsize=12, fontweight="bold", va="top")
    ax_c.text(0.0, -0.37, "Discovery: exploratory/post-selection context. Replication: exact two-sided sign tests, BH m=4.", transform=ax_c.transAxes, fontsize=6.8)

    fig.suptitle("Direct four-gene spatial replication in GSE312129", x=0.105, y=0.99, ha="left", fontsize=10.2, fontweight="bold")
    return save_figure(fig, 3)


def build_figure4(pseudo: pd.DataFrame, contrasts: pd.DataFrame) -> list[Path]:
    data = pseudo[(pseudo["condition"] == "SSc") & pseudo["gene"].isin(GENES)].copy()
    data["donor"] = data["sample_id"].map(donor_from_multiome)
    data["donor_order"] = data["donor"].str.replace("SSc", "", regex=False).astype(int)
    data = data.sort_values(["gene", "donor_order"])
    class_order = ["keratinocyte", "fibroblast", "endothelial_strict"]
    class_labels = ["Keratinocyte", "Fibroblast", "Endothelial"]
    annotation = {
        "NLRP1": "K>F 8/10, median +0.270\nK<E 10/10, median -0.915",
        "PYCARD": "K>F 10/10, median +1.738\nK>E 9/10, median +1.597",
        "CASP4": "K>F 7/10, median +0.097\nK<E 9/10, median -0.753",
        "GSDMD": "F>K 10/10, median K-F -1.925\nE>K 10/10, median K-E -3.184",
    }
    fig, axes = plt.subplots(2, 2, figsize=(7.2, 6.75), sharex=True)
    for ax, gene in zip(axes.flat, GENES):
        g = data[data["gene"] == gene]
        pivot = g.pivot(index="donor", columns="cell_type_strict", values="log2_CPM_plus_1").reindex(DONORS)
        if pivot[class_order].isna().any().any():
            raise RuntimeError(f"missing donor/class values for {gene}")
        for donor, row in pivot[class_order].iterrows():
            ax.plot(range(3), row.values, color="#B6B6B6", lw=0.75, alpha=0.75, zorder=1)
        for x, cell_class in enumerate(class_order):
            vals = pivot[cell_class].to_numpy()
            jitter = np.linspace(-0.07, 0.07, len(vals))
            ax.scatter(x + jitter, vals, s=24, color=CLASS_COLORS[cell_class], edgecolor="white", linewidth=0.45, zorder=3)
            ax.scatter(x, np.median(vals), marker="D", s=34, color="black", edgecolor="white", linewidth=0.5, zorder=4)
        ax.set_title(gene, loc="left", fontstyle="italic", fontweight="bold")
        ax.text(0.22, 1.015, annotation[gene].replace("\n", "; "), transform=ax.transAxes, ha="left", va="bottom", fontsize=6.5)
        ax.set_xticks(range(3), class_labels, rotation=18, ha="right")
        ax.set_ylabel("Donor-level log2(CPM + 1)")
        ax.grid(axis="y", color="#E5E5E5", lw=0.55)
        ax.spines[["top", "right"]].set_visible(False)
    axes[0, 0].text(-0.2, 1.11, "a", transform=axes[0, 0].transAxes, fontsize=12, fontweight="bold", va="top")
    axes[0, 1].text(-0.2, 1.11, "b", transform=axes[0, 1].transAxes, fontsize=12, fontweight="bold", va="top")
    axes[1, 0].text(-0.2, 1.11, "c", transform=axes[1, 0].transAxes, fontsize=12, fontweight="bold", va="top")
    axes[1, 1].text(-0.2, 1.11, "d", transform=axes[1, 1].transAxes, fontsize=12, fontweight="bold", va="top")
    fig.suptitle("Same-donor major-cell-class expression context", x=0.1, y=0.99, ha="left", fontsize=10.2, fontweight="bold")
    fig.text(0.1, 0.018, "All points are SSc donor-level pseudobulk values (n=10), not per-cell expression or cellular origin. Diamonds show medians.", fontsize=7)
    fig.tight_layout(rect=[0.03, 0.055, 0.99, 0.955], h_pad=2.3, w_pad=1.8)
    return save_figure(fig, 4)


def add_box(ax: plt.Axes, xy: tuple[float, float], width: float, height: float, text: str, face: str, edge: str = "#555555") -> None:
    box = FancyBboxPatch(xy, width, height, boxstyle="round,pad=0.018,rounding_size=0.015", facecolor=face, edgecolor=edge, linewidth=0.8)
    ax.add_patch(box)
    ax.text(xy[0] + width / 2, xy[1] + height / 2, text, ha="center", va="center", fontsize=7)


def build_figure5(pred: pd.DataFrame, summary: pd.DataFrame, loo: pd.DataFrame) -> list[Path]:
    m1 = pred[pred["model"] == "M1_DONOR_F_E_NUCLEUS_RATIO"].copy()
    m1["donor_order"] = m1["report_donor"].str.replace("SSc", "", regex=False).astype(int)
    m1 = m1.sort_values(["gene", "donor_order"])
    if len(m1) != 40 or len(summary) != 8:
        raise RuntimeError(f"unexpected reconstruction rows: M1={len(m1)}, summary={len(summary)}")

    fig = plt.figure(figsize=(7.2, 8.15))
    grid = fig.add_gridspec(3, 1, height_ratios=[1.25, 1.75, 1.4], hspace=0.52, left=0.105, right=0.98, top=0.945, bottom=0.08)

    ax_a = fig.add_subplot(grid[0, 0])
    ax_a.set_xlim(0, 1)
    ax_a.set_ylim(0, 1)
    ax_a.axis("off")
    add_box(ax_a, (0.02, 0.54), 0.25, 0.30, "Observed Visium layer direction\nEpidermis vs dermis\n(stored labels)", "#F4F4F4")
    ax_a.annotate("compare sign", xy=(0.47, 0.69), xytext=(0.36, 0.69), arrowprops={"arrowstyle": "->", "lw": 1.0}, ha="center", va="center", fontsize=7)
    add_box(ax_a, (0.48, 0.57), 0.22, 0.25, "M1 primary proxy\nEpidermis = K\nDermis = donor F:E ratio", "#E9F0FA", M1)
    add_box(ax_a, (0.74, 0.57), 0.22, 0.25, "M2 sensitivity proxy\nEpidermis = K\nDermis = 0.5F + 0.5E", "#FCF1E5", M2)
    ax_a.text(0.48, 0.38, "Class CPMs mixed on the linear scale, then transformed to log2(CPM+1)", fontsize=7)
    ax_a.text(0.02, 0.13, "Restricted proxy: keratinocyte, fibroblast and endothelial_strict only. It is not measured Visium spot composition.", fontsize=7.1, fontweight="bold")
    ax_a.text(-0.08, 1.02, "a", transform=ax_a.transAxes, fontsize=12, fontweight="bold", va="top")
    ax_a.set_title("Frozen direction-reconstruction model", loc="left")

    ax_b = fig.add_subplot(grid[1, 0])
    y_map = {gene: 3 - i for i, gene in enumerate(GENES)}
    x_map = {donor: i for i, donor in enumerate(DONORS)}
    for _, row in m1.iterrows():
        x = x_map[row["report_donor"]]
        y = y_map[row["gene"]]
        obs = int(row["observed_sign"])
        proxy = int(row["predicted_sign"])
        if obs != proxy:
            ax_b.add_patch(plt.Rectangle((x - 0.38, y - 0.34), 0.76, 0.68, facecolor="#F7ECE6", edgecolor="#B66A47", linewidth=0.45, hatch="//", zorder=0))
        ax_b.scatter(x - 0.14, y, marker="^" if obs > 0 else "v", s=40, color=EPI if obs > 0 else DERM, edgecolor="black", linewidth=0.35, zorder=3)
        ax_b.scatter(x + 0.14, y, marker="^" if proxy > 0 else "v", s=40, facecolor="white", edgecolor=EPI if proxy > 0 else DERM, linewidth=1.25, zorder=3)
    ax_b.set_xticks(range(10), DONORS, rotation=45, ha="right")
    ax_b.set_yticks([y_map[g] for g in GENES], GENES, fontstyle="italic")
    ax_b.set_xlim(-0.55, 9.55)
    ax_b.set_ylim(-0.55, 3.55)
    ax_b.grid(axis="x", color="#ECECEC", lw=0.5)
    ax_b.spines[["top", "right", "left", "bottom"]].set_visible(False)
    ax_b.tick_params(axis="both", length=0)
    ax_b.set_title("Observed and M1 proxy directions for every paired donor", loc="left")
    ax_b.text(-0.08, 1.07, "b", transform=ax_b.transAxes, fontsize=12, fontweight="bold", va="top")
    ax_b.legend(
        handles=[
            Line2D([0], [0], marker="^", color="none", markerfacecolor=EPI, markeredgecolor="black", markersize=6, label="observed epidermal"),
            Line2D([0], [0], marker="v", color="none", markerfacecolor=DERM, markeredgecolor="black", markersize=6, label="observed dermal"),
            Line2D([0], [0], marker="^", color="none", markerfacecolor="white", markeredgecolor=EPI, markersize=6, label="proxy epidermal"),
            Line2D([0], [0], marker="v", color="none", markerfacecolor="white", markeredgecolor=DERM, markersize=6, label="proxy dermal"),
        ],
        frameon=False,
        ncol=2,
        loc="upper right",
        bbox_to_anchor=(1.0, 1.16),
        columnspacing=0.8,
    )

    ax_c = fig.add_subplot(grid[2, 0])
    model_color = {"M1_DONOR_F_E_NUCLEUS_RATIO": M1, "M2_EQUAL_F_E": M2}
    offset = {"M1_DONOR_F_E_NUCLEUS_RATIO": 0.13, "M2_EQUAL_F_E": -0.13}
    for _, row in summary.iterrows():
        y = y_map[row["gene"]] + offset[row["model"]]
        count = int(row["n_exact_sign_concordant"])
        ax_c.scatter(count, y, s=43, color=model_color[row["model"]], edgecolor="white", linewidth=0.5, zorder=3)
        ax_c.text(count + 0.16, y, f"{count}/10", va="center", fontsize=7)
    grouped = loo.groupby(["model", "gene"])["n_exact_sign_concordant"].agg(["min", "max"]).reset_index()
    for _, row in grouped.iterrows():
        y = y_map[row["gene"]] + offset[row["model"]]
        color = model_color[row["model"]]
        ax_c.plot([row["min"], row["max"]], [y, y], color=color, lw=2.1, alpha=0.9)
    ax_c.set_yticks([y_map[g] for g in GENES], GENES, fontstyle="italic")
    ax_c.set_xlim(-0.4, 10.8)
    ax_c.set_ylim(-0.6, 3.6)
    ax_c.set_xticks(range(0, 11, 2))
    ax_c.set_xlabel("Exact sign concordance (of 10); horizontal segment = LOO range (of 9)")
    ax_c.set_title("M1 and M2 concordance with descriptive donor influence", loc="left")
    ax_c.grid(axis="x", color="#E5E5E5", lw=0.55)
    ax_c.spines[["top", "right", "left"]].set_visible(False)
    ax_c.tick_params(axis="y", length=0)
    ax_c.legend(
        handles=[
            Line2D([0], [0], marker="o", color=M1, markerfacecolor=M1, lw=2, label="M1 donor-specific F:E"),
            Line2D([0], [0], marker="o", color=M2, markerfacecolor=M2, lw=2, label="M2 equal F:E"),
        ],
        frameon=False,
        loc="upper right",
    )
    ax_c.text(-0.08, 1.08, "c", transform=ax_c.transAxes, fontsize=12, fontweight="bold", va="top")
    ax_c.text(0.0, -0.33, "NLRP1: observed more epidermal than M1 in 9/10; CASP4 in 6/10. Counts are descriptive; no P values.", transform=ax_c.transAxes, fontsize=6.8)

    fig.suptitle("Restricted same-donor direction reconstruction", x=0.105, y=0.99, ha="left", fontsize=10.2, fontweight="bold")
    return save_figure(fig, 5)


def main() -> None:
    require_inputs()
    OUT.mkdir(parents=True, exist_ok=True)
    WORK.mkdir(parents=True, exist_ok=True)
    QA.mkdir(parents=True, exist_ok=True)
    configure_style()
    renderer = renderer_record()

    vis_summary = pd.read_csv(FILES["vis_summary"][0], sep="\t")
    vis_sections = pd.read_csv(FILES["vis_sections"][0], sep="\t")
    pseudo = pd.read_csv(FILES["pseudo"][0], sep="\t")
    contrasts = pd.read_csv(FILES["contrasts"][0], sep="\t")
    pred = pd.read_csv(FILES["pred"][0], sep="\t")
    summary = pd.read_csv(FILES["summary"][0], sep="\t")
    loo = pd.read_csv(FILES["loo"][0], sep="\t")

    generated = {
        3: build_figure3(vis_sections, vis_summary),
        4: build_figure4(pseudo, contrasts),
        5: build_figure5(pred, summary, loo),
    }

    manifest = {
        "status": "PASS",
        "script": str(Path(__file__).relative_to(RUN)),
        "script_status": "PUBLIC_DETERMINISTIC_FIGURES_3_TO_5_GENERATOR",
        "script_sha256": sha256(Path(__file__)),
        "frozen_inputs": {key: {"path": str(path.relative_to(RUN)), "sha256": expected} for key, (path, expected) in FILES.items()},
        "new_figures": {
            str(number): [{"path": str(path.relative_to(RUN)), "sha256": sha256(path), "bytes": path.stat().st_size} for path in paths]
            for number, paths in generated.items()
        },
        "contracts": {
            "all_sections_or_donors_visible": True,
            "new_p_values": False,
            "raw_cross_platform_residual": "not plotted",
            "percent_explained": "not plotted",
            "r_squared": "not plotted",
            "proxy_is_not_measured_visium_composition": True,
        },
        "renderer": renderer,
        "figure_contract": {str(number): contract for number, contract in FIGURE_CONTRACT.items()},
        "generated_tiff_pixels": {
            str(number): pixel_record(paths[0]) for number, paths in generated.items()
        },
    }
    (QA / "FIGURE_BUILD_MANIFEST.json").write_text(json.dumps(manifest, indent=2) + "\n", encoding="utf-8")
    renderer_manifest = {
        "status": "PASS",
        "scope": "recorded build renderer; pixel identity is not guaranteed under other renderers",
        "script": str(Path(__file__).relative_to(RUN)),
        "script_sha256": sha256(Path(__file__)),
        "renderer": renderer,
        "figure_contract": manifest["figure_contract"],
        "generated_tiff_pixels": manifest["generated_tiff_pixels"],
    }
    (QA / "RENDERER_MANIFEST.json").write_text(
        json.dumps(renderer_manifest, indent=2) + "\n",
        encoding="utf-8",
    )
    print(json.dumps(manifest, indent=2))


if __name__ == "__main__":
    main()
```
