## Supplementary Software 1 for "Layer-to-cell-class correspondence is gene-specific for pyroptosis-related transcription in human skin": README.md

# Saved-value sensitivity reproduction

Portable reproduction code for the frozen post hoc endothelial-relative-abundance sensitivity analysis, using saved observation-level inputs. This is a new entry point derived from the historical computational contract, not the original single-use runner. It does not reproduce raw-data normalization, score construction or Cell2location fitting. The frozen reference files retain their original pre-adoption status fields as historical records.

The fixed analysis concerns the same 13 discovery sections (4 HC, 9 SSc) and 9,224 Dermis spots. It is not an independent cohort. The historical endothelial_score construction remains unverified. The new covariate is an expression-derived estimate, not a measured cell fraction; posterior uncertainty was not propagated.

Use the versions in environment.json. From the extracted software root, run:

```bash
python -B ifng_c2l/reproduce.py --output reproduced_ifng_c2l
```

The output directory must be new and outside ifng_c2l. The entry point reads only supplied TSV/JSON files, reproduces the three fixed coefficient series, the fixed bootstrap interval and the single fixed Wilcoxon test, and compares them with the frozen references. No alternative analysis is selected. See verification.json for the recorded comparison.

Inputs retain legacy section labels alongside report identifiers; the mapping comes from the unchanged Supporting Data SampleCrosswalk. The row index is an export index, not a spot barcode. The scope is saved intermediate values, not raw-data-to-result reproducibility.

This extension is provided as Supplementary Software 1, not as content of the already public Zenodo version 1.1.0. Historical pre-adoption fields are preserved unchanged. MIT applies to code; source data retain their original rights and attribution.
