## Supplementary Software 1 for "Layer-to-cell-class correspondence is gene-specific for pyroptosis-related transcription in human skin": reproduce.py

```
"""New portable entry point for fixed saved-value reproduction, not the historical runner."""
import argparse, csv, hashlib, json
from pathlib import Path
import numpy as np
from scipy.stats import rankdata, wilcoxon, PermutationMethod

BASE = Path(__file__).resolve().parent
FRACTION = 'c2l_rel_q05cell_abundance_w_sf_endothelial_strict'
COLS = ['unadjusted_rho','historical_score_adjusted_rho','c2l_fraction_adjusted_rho']

def correlation(a,b):
    a,b = np.asarray(a,dtype=float),np.asarray(b,dtype=float)
    ac,bc = a-a.mean(),b-b.mean()
    scale = np.linalg.norm(ac)*np.linalg.norm(bc)
    assert scale > 0 and np.isfinite(scale)
    return float(np.dot(ac,bc)/scale)

def partial(x,y,z):
    rx,ry,rz = (rankdata(v,method='average') for v in (x,y,z))
    design = np.column_stack([np.ones(len(x)),rz])
    assert np.linalg.matrix_rank(design) == 2
    ex = rx-design@np.linalg.lstsq(design,rx,rcond=None)[0]
    ey = ry-design@np.linalg.lstsq(design,ry,rcond=None)[0]
    assert np.std(ex)>1e-12 and np.std(ey)>1e-12
    return correlation(ex,ey),correlation(rx,ry)

def read(name):
    return list(csv.DictReader((BASE/name).read_text().splitlines(),delimiter='\t'))

def run(output):
    output = Path(output).resolve()
    assert not output.exists(), 'Output directory must not exist'
    assert not output.is_relative_to(BASE), 'Outputs must be outside the immutable input/code directory'
    inputs = read('data/DERMIS_INPUT_VALUES.tsv')
    cross = read('data/SECTION_ID_CROSSWALK.tsv')
    reference = read('reference/SECTION_CORRELATIONS.tsv')
    summary = json.loads((BASE/'reference/SUMMARY.json').read_text())
    assert len(inputs)==9224 and len(cross)==len(reference)==13
    assert [int(r['row_index']) for r in inputs]==list(range(9224))
    assert [r['source_section_id'] for r in cross]==[r['section'] for r in reference]
    records, errors = [],[]
    for cw,ref in zip(cross,reference):
        data = [r for r in inputs if r['source_section_id']==cw['source_section_id']]
        assert len(data)==int(cw['n_dermis'])==int(ref['n_dermis'])
        assert {r['report_section_id'] for r in data}=={cw['report_section_id']}
        assert {r['group'] for r in data}=={cw['group']}=={ref['group']}
        x,y,old,new = [np.array([float(r[k]) for r in data]) for k in ['IFNG_score','GSDMD_norm','endothelial_score',FRACTION]]
        oldrho,raw = partial(x,y,old)
        newrho,raw2 = partial(x,y,new)
        assert raw==raw2
        record = {'section':ref['section'],'group':ref['group'],'n_dermis':len(data),
                  COLS[0]:raw,COLS[1]:oldrho,COLS[2]:newrho,
                  'c2l_minus_unadjusted_rho':newrho-raw,'c2l_minus_historical_rho':newrho-oldrho}
        for k in COLS+['c2l_minus_unadjusted_rho','c2l_minus_historical_rho']:
            err = abs(record[k]-float(ref[k])); assert err<=1e-10,(ref['section'],k,err)
            errors.append(err)
        records.append(record)
    descriptions = {}
    for k in COLS:
        v = np.array([r[k] for r in records])
        descriptions[k]={'median':float(np.median(v)),'min':float(v.min()),'max':float(v.max()),
            'positive_sections':int((v>0).sum()),'zero_sections':int((v==0).sum()),'negative_sections':int((v<0).sum())}
        for field,value in descriptions[k].items():
            expected=summary['descriptive'][k][field]
            assert (value==expected if isinstance(value,int) else abs(value-expected)<=1e-10)
    v=np.array([r[COLS[2]] for r in records])
    assert np.all(v!=0) and len(np.unique(np.abs(v)))==13
    result=wilcoxon(v,alternative='two-sided',zero_method='wilcox',correction=False,
                    method=PermutationMethod(n_resamples=np.inf,batch=8192))
    assert float(result.statistic)==summary['sole_new_test']['statistic']
    assert abs(float(result.pvalue)-summary['sole_new_test']['pvalue'])<=1e-15
    rng=np.random.default_rng(42)
    ci=np.percentile([np.median(rng.choice(v,size=13,replace=True)) for _ in range(2000)],[2.5,97.5])
    assert np.max(np.abs(ci-summary['new_median_bootstrap_95ci']))<=1e-10
    verification={'status':'PASS','purpose':'Reproduction QA only; frozen references remain authoritative',
        'input_rows':9224,'section_n':13,'coefficients_compared':39,'max_coefficient_or_delta_error':max(errors),
        'descriptive':descriptions,'median_bootstrap_95ci':ci.tolist(),
        'wilcoxon':{'statistic':float(result.statistic),'pvalue':float(result.pvalue),'nonzero_n':13,'zero_n':0,'absolute_ties':False},
        'reference_summary_sha256':hashlib.sha256((BASE/'reference/SUMMARY.json').read_bytes()).hexdigest(),
        'reference_sections_sha256':hashlib.sha256((BASE/'reference/SECTION_CORRELATIONS.tsv').read_bytes()).hexdigest(),
        'historical_runner_executed':False,'new_scientific_results_adopted_from_reproduction':False}
    output.mkdir(parents=True,exist_ok=False)
    (output/'verification.json').write_text(json.dumps(verification,indent=2,allow_nan=False)+'\n')
    with (output/'reproduced_sections.tsv').open('w',newline='') as f:
        w=csv.DictWriter(f,fieldnames=list(records[0]),delimiter='\t',lineterminator='\n'); w.writeheader(); w.writerows(records)
    print(json.dumps(verification))

if __name__=='__main__':
    p=argparse.ArgumentParser(description=__doc__)
    p.add_argument('--output',required=True,help='A new output directory outside the input/code directory')
    run(p.parse_args().output)
```
