## Supplementary Software 1 for "Layer-to-cell-class correspondence is gene-specific for pyroptosis-related transcription in human skin": PROVENANCE.md

# Plotting Provenance

This software supplement supplies additional display code, not a new scientific
analysis or a new public Zenodo version. The source hashes in FIGURE_CODE_MAP.tsv
identify the frozen predecessors. Their role-relative identifiers are provenance
labels, not paths required on a recipient's computer.

- Figure 5: the included generator is byte-identical to the fixed layout-only
  predecessor. A separate launcher selects Figure 5 only. Compared with the
  public v1.1.0 generator, only the compare-sign label x coordinate changed from
  0.29 to 0.36. No plotted data, axes or other artists changed.
- S8D: Replacement plotting code for Supplementary Figure S8D, generated from frozen source values. The original plotting script was not recovered.
  The render function is unchanged from that replacement script. Only font/input
  arguments and launch settings were made portable. The 20 points, 10 links,
  two display medians and 10 labels reproduce the recorded artist ledger.
- S10A: the fixed pure drawing functions and composition geometry were extracted
  without importing the historical analysis runner. Font selection is explicit.
  The saved model summaries and the previously retained immune-row label are
  unchanged. Upper-panel and lower-panel reference regions are preserved.
- S14: constants and eight drawing functions were extracted without invoking
  raw-data extraction. A loader reads the saved spot and donor tables. The
  native atlas, 20-label overlay and final header/page-placement transformation
  are separate reproducible steps. Comparison to the final figure uses its
  content region, not the surrounding Supplement prose.

Only paths, launch control, font arguments and provenance were adapted. The
drawing-function AST comparisons and rendered pixels matched their references.
S8D is a replacement renderer; S14 is an extracted saved-display renderer.
Neither is described as the recovered historical analysis runner.

The validation software reads no raw expression matrices and computes no new
P values, q values, correlations, regression coefficients or scientific means.
Coordinate bounds/radii and the already defined S8D display medians are drawing
operations. The submitted figures remain unchanged.

## Saved-value endothelial-relative-abundance sensitivity

Portable reproduction code for the frozen post hoc endothelial-relative-abundance sensitivity analysis, using saved observation-level inputs. This is a new entry point derived from the historical computational contract, not the original single-use runner. It does not reproduce raw-data normalization, score construction or Cell2location fitting. The frozen reference files retain their original pre-adoption status fields as historical records.

See ifng_c2l/SOURCE_PROVENANCE.json for source-file hashes, allowed observation fields, denominator definition and export hashes. See ifng_c2l/verification.json for the exact saved-result comparison.
