## Supplementary Software 1 for "Layer-to-cell-class correspondence is gene-specific for pyroptosis-related transcription in human skin": README.md

# Supplementary Software 1

Additional plotting code and frozen display inputs for Figure 5 and
Supplementary Figures S8D, S10A and S14. This bundle also includes portable
reproduction code and saved inputs for the separate post hoc endothelial-
relative-abundance sensitivity analysis. The accompanying manuscript is
"Layer-to-cell-class correspondence is gene-specific for pyroptosis-related
transcription in human skin".

The analysis repository is Zenodo version 1.1.0,
doi:10.5281/zenodo.22215741. This additional software is a separate supplement,
not a new version of that deposit. It is prepared for journal review; its local
creation does not imply upload, journal acceptance or public publication.

## Reproduce the Displays

Use Python 3.12.2 and the versions in environment/versions.json. Set PYTHON to
that executable. Set FONT_REGULAR to the locally available Arial regular font,
FONT_BOLD to Arial Bold, and FONT_SERIF to Liberation Serif Regular. Font files
are not redistributed; their reference names and hashes are in the environment
record. These fonts are needed for the recorded pixel comparisons. Different
fonts, operating systems or rendering-library versions can change pixels.

Run from the extracted software directory. Outputs go to a new directory of
your choosing. The commands below read only the supplied tables and images.
They do not download data, read an AnnData matrix or repeat statistical tests.

```bash
export PYTHONDONTWRITEBYTECODE=1
export MPLBACKEND=Agg
export MPLCONFIGDIR=reproduced/cache/mpl
export XDG_CACHE_HOME=reproduced/cache/xdg
"$PYTHON" -B figure5/render.py --output reproduced/figure5 --font "$FONT_REGULAR"
"$PYTHON" -B s8d/plot_s8d_from_frozen_values.py --output reproduced/s8d --font "$FONT_REGULAR"
"$PYTHON" -B s8d/assemble.py --panel reproduced/s8d/S8D.png --output reproduced/s8d/S08_replacement.png
"$PYTHON" -B s10a/render.py --output reproduced/s10a --font "$FONT_REGULAR" --bold-font "$FONT_BOLD"
"$PYTHON" -B s14/render.py --output reproduced/s14 --font "$FONT_REGULAR"
"$PYTHON" -B s14/place_final.py --atlas reproduced/s14/atlas_with_saved_means.pdf --output reproduced/s14/atlas_final_placement.pdf --serif-font "$FONT_SERIF"
```

Figure 5 calls only the Figure 5 function, retaining the frozen data and the
one-coordinate label correction. S8D supplies 20 saved layer means from 10
sections; its two median lines are display coordinates, not new scientific
estimates. S10A reads saved model summaries and reproduces the fixed lower-panel
layout while retaining the other image regions. S14 uses 10,217 saved spot
records, 10 donor display rows and 20 saved layer-mean labels. Its coordinate
processing determines circle radius and plot bounds, not expression estimates.
The native atlas, later labels and final page placement are separate steps.

## Provenance and Verification

Replacement plotting code for Supplementary Figure S8D, generated from frozen source values. The original plotting script was not recovered.

S8D is not presented as a recovered historical script. S14 is an extracted
renderer taking saved display tables, not a reproduction of raw-data extraction
or the original scientific analysis. The original source hashes and portable
entry points are listed in FIGURE_CODE_MAP.tsv and PROVENANCE.md.

Each component includes comparison assets. Pixel comparisons passed in the
recorded environment; see environment/verification.json. The submitted Figure 5
and Supplement were not replaced by these validation outputs. The full
Supplement text and unrelated source analyses are outside this plotting bundle.

MANIFEST.tsv lists every file except itself with byte size and SHA-256. MIT
applies to the code as stated in LICENSE, not automatically to third-party
data, comparison figures or fonts; source data remain subject to their source
repository terms.

## Reproduce the Separate Post Hoc Sensitivity Analysis

See ifng_c2l/README.md. Unlike the display commands above, this entry point
reproduces the fixed statistical calculations from supplied observation-level
values, including the single Wilcoxon test and section-median bootstrap.
It does not fit Cell2location, reconstruct scores, download data or import the
historical single-use runner. Its scope and environment are separate from
the plotting components and the public Zenodo version 1.1.0.

## Supplementary Table S9: post hoc dermal-weight sensitivity

The s9/ directory contains a standalone Python 3 script, frozen Supporting Data
extracts, the donor-level output, and a verification receipt. From an extracted
copy of this ZIP, run `python3 s9/weight_sweep.py`. The script compares its M1
and M2 controls and every fixed-weight row against frozen expected values before
writing results. It does not run new differential expression or compute P values.
The endothelial share w is an assumed linear-CPM mixing weight, not a measured
cell or RNA fraction in the ten replication sections. The q05 reference row is
from discovery Supporting Data Fig4C and is descriptive only.
