## Supplementary Software 1 for "Layer-to-cell-class correspondence is gene-specific for pyroptosis-related transcription in human skin": assemble.py

```
import argparse
from pathlib import Path
from PIL import Image
p=argparse.ArgumentParser();p.add_argument('--panel',type=Path,required=True);p.add_argument('--output',type=Path,required=True);a=p.parse_args()
old=Image.open(Path(__file__).parent/'reference/S08_parent.png').convert('RGB')
panel=Image.open(a.panel).convert('RGB');assert panel.size==(1000,1360)
old.paste(panel,(1550,1510));old.save(a.output)
```
