## Supplementary Software 1 for "Layer-to-cell-class correspondence is gene-specific for pyroptosis-related transcription in human skin": plot_s8d_from_frozen_values.py

```
"""Replacement plotting code for Supplementary Figure S8D, generated from frozen source values. The original plotting script was not recovered."""
import argparse, csv, hashlib, json, platform, sys
from decimal import Decimal
from pathlib import Path
import matplotlib
matplotlib.use('Agg')
import matplotlib.pyplot as plt
from matplotlib.font_manager import FontProperties
import numpy as np
from PIL import Image

PROVENANCE=__doc__
FONT=None

def tsv(path,rows):
    with path.open('w',encoding='utf-8',newline='') as f:
        w=csv.DictWriter(f,fieldnames=list(rows[0]),delimiter='\t',lineterminator='\n');w.writeheader();w.writerows(rows)

def render(inp,out):
    out.mkdir(parents=True,exist_ok=True)
    rows=list(csv.DictReader(inp.open(),delimiter='\t'));assert len(rows)==10
    ids=[f'GSM{9338170+i}_SSC{i}_visium' for i in range(1,11)]
    assert [r['section_id'] for r in rows]==ids
    d=[Decimal(r['dermal_mean_log1p']) for r in rows];e=[Decimal(r['epidermal_mean_log1p']) for r in rows]
    md=(sorted(d)[4]+sorted(d)[5])/2;me=(sorted(e)[4]+sorted(e)[5])/2
    assert md==Decimal('0.19811430945') and me==Decimal('0.35050543145')
    fp=FontProperties(fname=FONT,size=18)
    fig=plt.figure(figsize=(6.25,8.5),dpi=160,facecolor='white')
    ax=fig.add_axes([.22,.14,.49,.74]);ax.set_xlim(-.18,1.18);ax.set_ylim(.11383627857,.53402374803)
    ax.set_xticks([0,1],['Dermis','Epidermis']);ax.set_yticks([.15,.20,.25,.30,.35,.40,.45,.50])
    ax.spines[['top','right']].set_visible(False);ax.grid(axis='y',color='#dce0e6',linewidth=.65);ax.set_axisbelow(True)
    ax.set_ylabel('Signed triad mean log1p expression',fontproperties=fp,labelpad=7)
    ax.tick_params(axis='both',length=3)
    for t in ax.get_xticklabels()+ax.get_yticklabels():t.set_fontproperties(fp)
    title=fig.text(.55,.94,'Signed triad paired by section',ha='center',va='center',fontproperties=FontProperties(fname=FONT,size=16))
    panel=fig.text(.025,.97,'D',ha='left',va='top',fontproperties=FontProperties(fname=FONT,size=28,weight='bold'))
    donor=[]
    for r,di,ei in zip(rows,d,e):
        ln,=ax.plot([0,1],[float(di),float(ei)],'-o',color='#e64b45',linewidth=1.0,markersize=4,markeredgewidth=.4,markeredgecolor='#a83d39',zorder=3)
        donor.append(ln)
    med,=ax.plot([0,1],[float(md),float(me)],'-D',color='#111827',linewidth=2.0,markersize=6,label='Median',zorder=4)
    ax.legend(loc='upper left',frameon=False,prop=FontProperties(fname=FONT,size=14),handlelength=1.6,borderpad=.15)
    fig.canvas.draw();renderer=fig.canvas.get_renderer()
    order=sorted(range(10),key=lambda i:(e[i],i))
    xy=[ax.transData.transform((1,float(e[i]))) for i in order]
    gap=47.0;yy=[]
    for pt in xy:yy.append(max(pt[1],yy[-1]+gap if yy else pt[1]))
    ceiling=ax.bbox.y1-12
    if yy[-1]>ceiling:
        yy[-1]=ceiling
        for j in range(8,-1,-1):yy[j]=min(yy[j],yy[j+1]-gap)
    assert yy[0]>ax.bbox.y0
    labels=[]
    for i,y in zip(order,yy):
        end=ax.transData.transform((1,float(e[i])))/np.array(fig.canvas.get_width_height())
        labelxy=(.845,y/fig.bbox.height)
        annotation=ax.annotate('',xy=(1,float(e[i])),xycoords='data',xytext=(labelxy[0]-.01,labelxy[1]),textcoords=fig.transFigure,
                               arrowprops={'arrowstyle':'-','color':'#777777','linewidth':.6},annotation_clip=False)
        text=fig.text(*labelxy,rows[i]['label'],fontproperties=fp,va='center',ha='left',color='#b43f39')
        labels.append((i,text))
    fig.canvas.draw();renderer=fig.canvas.get_renderer()
    bboxes=[t.get_window_extent(renderer) for _,t in labels]
    assert all(b.width>0 and b.x1<fig.bbox.x1 and b.y0>0 and b.y1<fig.bbox.y1 for b in bboxes)
    assert all(not a.overlaps(b) for i,a in enumerate(bboxes) for b in bboxes[i+1:])
    ledger=[]
    for r,line in zip(rows,donor):
        xs,ys=line.get_data()
        for j,layer in enumerate(('Dermis','Epidermis')):
            ledger.append({'kind':'point','section_id':r['section_id'],'label':layer,'x':float(xs[j]),'y':float(ys[j]),'x2':'','y2':''})
        ledger.append({'kind':'connection','section_id':r['section_id'],'label':'paired','x':float(xs[0]),'y':float(ys[0]),'x2':float(xs[1]),'y2':float(ys[1])})
    xs,ys=med.get_data()
    for j,layer in enumerate(('Dermis','Epidermis')):ledger.append({'kind':'display_median','section_id':'cohort','label':layer,'x':float(xs[j]),'y':float(ys[j]),'x2':'','y2':''})
    for i,t in labels:
        x,y=t.get_position();ledger.append({'kind':'label','section_id':rows[i]['section_id'],'label':t.get_text(),'x':x,'y':y,'x2':'','y2':''})
    tsv(out/'S8D_PLOTTED_ELEMENTS.tsv',ledger)
    tsv(out/'S8D_DISPLAY_MEDIAN_COORDINATES.tsv',[{'layer':x,'value':str(v),'definition':'derived display coordinate: (5th+6th ordered frozen means)/2; not median_delta'} for x,v in [('Dermis',md),('Epidermis',me)]])
    fig.savefig(out/'S8D.png',dpi=160,facecolor='white');fig.savefig(out/'S8D.pdf',metadata={'Title':'Replacement S8D from frozen display values','Subject':PROVENANCE})
    style={'provenance':PROVENANCE,'public_deposit_status':'LOCAL_UNPUBLISHED; not included in Zenodo v1.1.0','executable':sys.executable,'python':platform.python_version(),'matplotlib':matplotlib.__version__,'numpy':np.__version__,'font':FONT,'font_sha256':hashlib.sha256(Path(FONT).read_bytes()).hexdigest(),'image_pixels':[1000,1360],'donor_rgba':[.9019607843137255,.29411764705882354,.27058823529411763,1.0],'donor_linewidth':1.0,'marker_size':4,'median_color':'#111827','median_linewidth':2.0,'label_gap_pixels':gap,'label_font_points':18,'xlim':list(ax.get_xlim()),'ylim':list(ax.get_ylim()),'ticks':list(ax.get_yticks()),'artist_records':len(ledger),'label_bbox_pixels':[list(b.bounds) for b in bboxes]}
    (out/'ENVIRONMENT.json').write_text(json.dumps(style,indent=2)+'\n');plt.close(fig)

if __name__=='__main__':
    p=argparse.ArgumentParser();p.add_argument('--input',type=Path,default=Path(__file__).parent/'data/S8D_FROZEN_DISPLAY_VALUES.tsv');p.add_argument('--output',type=Path,required=True);p.add_argument('--font',type=Path,required=True);a=p.parse_args();FONT=str(a.font);render(a.input,a.output)
```
