## Supplementary Software 1 for "Layer-to-cell-class correspondence is gene-specific for pyroptosis-related transcription in human skin": README.md

# Supplementary Table S9 reproduction

Run `python3 s9/weight_sweep.py` from any current directory after extraction.
Inputs in `data/` are frozen extracts from the accompanying Supporting Data
workbook (ReplicationDonorCrosswalk, MultiomePseudobulk,
Rep4GeneSections, Fig4C). Outputs under `results/` were produced by the script.

The script checks M1=1/9/4/8, M2=0/9/3/8, all fixed-weight grid rows, and the
discovery q05 reference row before writing. The analysis is post hoc and
descriptive; no P values are computed. `results/verification.json` contains
input hashes and the PASS receipt. The linear-CPM weight w is assumed, not
estimated from replication tissue composition.
