## Supplementary Software 1 for "Layer-to-cell-class correspondence is gene-specific for pyroptosis-related transcription in human skin": weight_sweep.py

```
"""Reproduce descriptive dermal-weight sign concordance from frozen TSV inputs.

The four input TSVs live next to this script in ``data/``. No raw-expression
matrix is needed. This analysis is post hoc and provides no P values.
"""

from __future__ import annotations

import csv
import hashlib
import json
import math
from pathlib import Path


GENES = ("NLRP1", "PYCARD", "CASP4", "GSDMD")
GRID = (0.0, 0.05, 0.10, 0.15, 0.20, 0.25, 0.30, 0.40, 0.50, 0.75, 1.0)
EXPECTED_GRID = (
    (8, 9, 4, 8), (7, 9, 3, 8), (7, 9, 2, 8), (5, 9, 3, 8),
    (4, 9, 2, 8), (4, 9, 2, 8), (2, 9, 2, 8), (1, 9, 3, 8),
    (0, 9, 3, 8), (0, 9, 4, 8), (0, 10, 4, 8),
)
EXPECTED_M1 = (1, 9, 4, 8)
EXPECTED_M2 = (0, 9, 3, 8)
EXPECTED_REFERENCE = (7, 9, 2, 8)


def tsv_rows(path: Path) -> list[dict[str, str]]:
    with path.open(encoding="utf-8", newline="") as stream:
        return list(csv.DictReader(stream, delimiter="\t"))


def sha256(path: Path) -> str:
    return hashlib.sha256(path.read_bytes()).hexdigest()


def sign(value: float) -> int:
    return (value > 0) - (value < 0)


def write_tsv(path: Path, fieldnames: list[str], rows: list[dict]) -> None:
    with path.open("w", encoding="utf-8", newline="") as stream:
        writer = csv.DictWriter(stream, fieldnames=fieldnames, delimiter="\t", lineterminator="\n")
        writer.writeheader()
        writer.writerows(rows)


def main() -> None:
    here = Path(__file__).resolve().parent
    inputs = {
        "crosswalk": here / "data/replication_donor_crosswalk.tsv",
        "pseudobulk": here / "data/multiome_pseudobulk.tsv",
        "visium": here / "data/replication_visium_4gene.tsv",
        "reference": here / "data/discovery_dermis_reference.tsv",
    }
    crosswalk = tsv_rows(inputs["crosswalk"])
    multi_rows = tsv_rows(inputs["pseudobulk"])
    visium_rows = tsv_rows(inputs["visium"])
    reference_rows = tsv_rows(inputs["reference"])
    if len(crosswalk) != 10 or len(multi_rows) != 120 or len(visium_rows) != 40:
        raise RuntimeError("Unexpected frozen input cardinality")
    if {r["gene"] for r in visium_rows} != set(GENES):
        raise RuntimeError("Four-gene input mismatch")

    multi = {(r["sample_id"], r["gene"], r["cell_type_strict"]): r for r in multi_rows}
    visium = {(r["sample_id"], r["gene"]): r for r in visium_rows}
    if len(multi) != 120 or len(visium) != 40:
        raise RuntimeError("Duplicate input key")
    reference = {r["cell_type"]: float(r["mean_rel_abundance"]) for r in reference_rows}
    reference_w = reference["endothelial_strict"] / (
        reference["fibroblast"] + reference["endothelial_strict"]
    )

    def calculate(label: str, fixed_w: float | None) -> tuple[tuple[int, ...], list[dict]]:
        details: list[dict] = []
        totals = {gene: 0 for gene in GENES}
        for mapping in crosswalk:
            vis_id = mapping["visium_sample_id"]
            mul_id = mapping["multiome_sample_id"]
            for gene in GENES:
                vals = {
                    cell: multi[(mul_id, gene, cell)]
                    for cell in ("keratinocyte", "fibroblast", "endothelial_strict")
                }
                k = float(vals["keratinocyte"]["CPM_retained_features"])
                f = float(vals["fibroblast"]["CPM_retained_features"])
                e = float(vals["endothelial_strict"]["CPM_retained_features"])
                nf = int(vals["fibroblast"]["n_nuclei"])
                ne = int(vals["endothelial_strict"]["n_nuclei"])
                weight = fixed_w if fixed_w is not None else ne / (nf + ne)
                measured = visium[(vis_id, gene)]
                observed_delta = (
                    float(measured["epidermis_mean_log1p"])
                    - float(measured["dermis_mean_log1p"])
                )
                proxy_delta = math.log2(k + 1) - math.log2((1 - weight) * f + weight * e + 1)
                if sign(observed_delta) == 0 or sign(proxy_delta) == 0:
                    raise RuntimeError(f"Zero sign at {label}/{mapping['report_donor']}/{gene}")
                concordant = int(sign(observed_delta) == sign(proxy_delta))
                totals[gene] += concordant
                details.append({
                    "weight_label": label,
                    "endothelial_weight": f"{weight:.12f}",
                    "donor": mapping["report_donor"],
                    "visium_sample_id": vis_id,
                    "multiome_sample_id": mul_id,
                    "gene": gene,
                    "observed_sign": sign(observed_delta),
                    "proxy_sign": sign(proxy_delta),
                    "concordant": concordant,
                    "keratinocyte_cpm": f"{k:.12f}",
                    "fibroblast_cpm": f"{f:.12f}",
                    "endothelial_cpm": f"{e:.12f}",
                    "fibroblast_nuclei": nf,
                    "endothelial_nuclei": ne,
                })
        return tuple(totals[g] for g in GENES), details

    m1, m1_detail = calculate("M1", None)
    m2, _ = calculate("0.50", 0.50)
    if m1 != EXPECTED_M1 or m2 != EXPECTED_M2:
        raise RuntimeError(f"M1/M2 control failure: M1={m1}, M2={m2}")

    summary: list[dict] = []
    all_details: list[dict] = []
    for weight, expected in zip(GRID, EXPECTED_GRID):
        label = f"{weight:.2f}"
        counts, detail = calculate(label, weight)
        if counts != expected:
            raise RuntimeError(f"Expected-grid failure at {label}: {counts} != {expected}")
        summary.append({"weight_label": label, "endothelial_weight": f"{weight:.12f}", **dict(zip(GENES, counts)), "scope": "replication fixed weight"})
        all_details.extend(detail)
    summary.append({"weight_label": "M1", "endothelial_weight": "donor-specific", **dict(zip(GENES, m1)), "scope": "nucleus-ratio control"})
    all_details.extend(m1_detail)
    reference_counts, reference_detail = calculate("discovery_reference", reference_w)
    if reference_counts != EXPECTED_REFERENCE:
        raise RuntimeError(f"Discovery reference failure: {reference_counts} != {EXPECTED_REFERENCE}")
    summary.append({"weight_label": "discovery_reference", "endothelial_weight": f"{reference_w:.12f}", **dict(zip(GENES, reference_counts)), "scope": "discovery q05 descriptive reference"})
    all_details.extend(reference_detail)

    out = here / "results"
    out.mkdir(exist_ok=True)
    write_tsv(out / "weight_sweep_summary.tsv", ["weight_label", "endothelial_weight", *GENES, "scope"], summary)
    write_tsv(out / "donor_sign_concordance.tsv", list(all_details[0]), all_details)
    verification = {
        "status": "PASS",
        "analysis_role": "post hoc descriptive sensitivity; no P values",
        "donors": len(crosswalk),
        "genes": list(GENES),
        "input_sha256": {name: sha256(path) for name, path in inputs.items()},
        "controls": {"M1": dict(zip(GENES, m1)), "M2": dict(zip(GENES, m2))},
        "expected_grid_match": True,
        "reference_w": reference_w,
        "reference_counts": dict(zip(GENES, reference_counts)),
        "n_donor_rows": len(all_details),
    }
    (out / "verification.json").write_text(json.dumps(verification, indent=2) + "\n", encoding="utf-8")
    print(json.dumps(verification, indent=2))


if __name__ == "__main__":
    main()
```
