## Supplementary Software 1 for "Layer-to-cell-class correspondence is gene-specific for pyroptosis-related transcription in human skin": render.py

```
import argparse, textwrap
from pathlib import Path
from typing import Any
import pandas as pd
from PIL import Image,ImageDraw,ImageFont
REGULAR=BOLD=None
def font(size: int, bold: bool=False):
    return ImageFont.truetype(str(BOLD if bold else REGULAR),size=size)

def fmt(x: Any, digits: int=3) -> str:
    try:
        return f'{float(x):.{digits}f}'
    except Exception:
        return str(x)

def create_active_figure(long_df: pd.DataFrame) -> tuple[Path, Path]:
    img = Image.new('RGB', (2400, 1820), 'white')
    draw = ImageDraw.Draw(img)
    black = (32, 34, 36)
    gray = (92, 102, 112)
    light = (232, 236, 239)
    blue = (37, 99, 160)
    orange = (198, 112, 27)
    green = (45, 128, 95)
    purple = (126, 87, 194)
    draw.text((100, 70), 'Fig. S10A. Dermal GSDMD model-summary analyses', fill=black, font=font(58, True))
    draw.text((100, 145), 'Section-level summaries across 13 dermal sections', fill=gray, font=font(32))
    p_rows = long_df[long_df['analysis_id'].isin(['P1', 'P2', 'S1'])].copy()
    s2_rows = long_df[long_df['analysis_id'].astype(str).str.startswith('S2_')].copy()

    def axis_block(rows: pd.DataFrame, x0: int, y0: int, w: int, h: int, title: str, unit: str, color: tuple[int, int, int]):
        draw.rectangle((x0, y0, x0 + w, y0 + h), outline=light, width=3)
        draw.text((x0 + 35, y0 + 25), title, fill=black, font=font(36, True))
        vals = []
        for _, r in rows.iterrows():
            vals.extend([float(r['min_estimate']), float(r['max_estimate'])])
        mn = min(-0.05, min(vals) - 0.03)
        mx = max(0.25, max(vals) + 0.03)
        left = x0 + 420
        right = x0 + w - 120
        top = y0 + 120
        row_gap = 155 if len(rows) <= 3 else 115

        def mapx(v: float) -> int:
            return int(left + (v - mn) / (mx - mn) * (right - left))
        y_axis = y0 + h - 85
        draw.line((left, y_axis, right, y_axis), fill=gray, width=3)
        for tick in [mn, 0, mx]:
            tx = mapx(tick)
            draw.line((tx, y_axis - 8, tx, y_axis + 8), fill=gray, width=2)
            draw.text((tx - 40, y_axis + 16), fmt(tick, 2), fill=gray, font=font(24))
        draw.text((left + 390, y_axis + 50), unit, fill=gray, font=font(24))
        if mn < 0 < mx:
            zx = mapx(0)
            draw.line((zx, top - 30, zx, y_axis), fill=(190, 190, 190), width=2)
        for i, (_, r) in enumerate(rows.iterrows()):
            y = top + i * row_gap
            label = str(r['analysis_label']).replace('P1: ', '').replace('P2: ', '').replace('S1: ', '')
            if str(r['analysis_id']) == 'S2_IFNG':
                label = 'S2: IFNG coefficient'
            elif str(r['analysis_id']) == 'S2_endothelial':
                label = 'S2: endothelial coefficient'
            elif str(r['analysis_id']) == 'S2_immune':
                label = 'S2: immune coefficient'
            draw.text((x0 + 35, y - 24), label, fill=black, font=font(29, True))
            draw.text((x0 + 35, y + 15), 'median 0.003; IQR -0.034–0.018' if str(r['analysis_id']) == 'S2_immune' else f'median {fmt(r['median_estimate'])}; IQR {fmt(r['iqr_low'])} to {fmt(r['iqr_high'])}', fill=gray, font=font(24))
            x_min = mapx(float(r['min_estimate']))
            x_max = mapx(float(r['max_estimate']))
            x_lo = mapx(float(r['iqr_low']))
            x_hi = mapx(float(r['iqr_high']))
            x_med = mapx(float(r['median_estimate']))
            draw.line((x_min, y, x_max, y), fill=(185, 190, 195), width=4)
            draw.line((x_lo, y, x_hi, y), fill=color, width=12)
            draw.ellipse((x_med - 13, y - 13, x_med + 13, y + 13), fill=color, outline=black, width=2)
            draw.text((right + 20, y - 16), fmt(r['median_estimate']), fill=black, font=font(27, True))
    axis_block(p_rows, 95, 245, 2210, 600, 'Partial Spearman summaries', 'estimate unit: partial rho', blue)
    axis_block(s2_rows, 95, 905, 2210, 610, 'Ranked linear-model summaries', 'estimate unit: ranked linear-model beta', orange)
    foot = 'Variables: GSDMD_norm, IFNG_score, endothelial_score, mk_immune. P1: GSDMD-IFNG | endothelial; P2: GSDMD-endothelial | IFNG; S1: GSDMD-IFNG | endothelial + immune; S2: ranked LM: IFNG / endothelial / immune. No significance stars shown.'
    lines = textwrap.wrap(foot, width=135)
    for i, line in enumerate(lines):
        draw.text((100, 1580 + i * 38), line, fill=gray, font=font(27))
    return img

def assemble(replacement,output):
    current=Image.open(Path(__file__).parent/'reference/S10_parent.png').convert('RGB')
    scale=(516.8701*600/72)/3000
    ax0=round((47.56497+120*516.8701/3000)*600/72)
    ay0=round((36+120*720/4179)*600/72)
    width=round(2760*scale);height=round(2093*scale);oldheight=round(1955*scale)
    anew=replacement.resize((width,height),Image.Resampling.LANCZOS)
    boundary=ay0+round(900*1.15*scale)
    oldend=ay0+oldheight;newend=ay0+height;delta=newend-oldend
    out=Image.new('RGB',(5100,6600+delta),'white')
    out.paste(current.crop((0,0,5100,boundary)),(0,0))
    out.paste(anew.crop((0,boundary-ay0,width,height)),(ax0,boundary))
    out.paste(current.crop((0,oldend,5100,6600)),(0,newend))

    out.save(output/'S10_replacement.png')

if __name__=='__main__':
    p=argparse.ArgumentParser();p.add_argument('--output',type=Path,required=True);p.add_argument('--font',type=Path,required=True);p.add_argument('--bold-font',type=Path,required=True);a=p.parse_args()
    REGULAR=a.font;BOLD=a.bold_font;a.output.mkdir(parents=True,exist_ok=True)
    df=pd.read_csv(Path(__file__).parent/'data/S10A_MODEL_SUMMARIES.csv')
    panel=create_active_figure(df);panel.save(a.output/'S10A_replacement_native.png')
    assemble(panel,a.output)
```
