## Supplementary Software 1 for "Layer-to-cell-class correspondence is gene-specific for pyroptosis-related transcription in human skin": place_final.py

```
"""Reproduce the fixed S14 header and final figure placement, not the Supplement."""
import argparse,json
from pathlib import Path
import fitz
p=argparse.ArgumentParser()
p.add_argument('--atlas',type=Path,required=True)
p.add_argument('--output',type=Path,required=True)
p.add_argument('--serif-font',type=Path,required=True)
a=p.parse_args()
doc=fitz.open(a.atlas)
placements=json.loads((Path(__file__).parent/'data/FINAL_PLACEMENT.json').read_text())
out=fitz.open()
for j,placement in enumerate(placements):
    page=doc[j]
    header='Supplementary Figure S14'+(' (continued)' if j else '')
    page.insert_text((545,17),header,fontname='S14Serif',fontfile=str(a.serif_font),fontsize=10)
    target=out.new_page(width=792,height=612)
    target.show_pdf_page(fitz.Rect(placement['rect']),doc,j,clip=fitz.Rect(placement['clip']),keep_proportion=True)
out.save(a.output,deflate=True)
```
