## Supplementary Software 1 for "Layer-to-cell-class correspondence is gene-specific for pyroptosis-related transcription in human skin": atlas_native.pdf

### *NLRP*1 **spatial / cell-class atlas: all 10 donors**

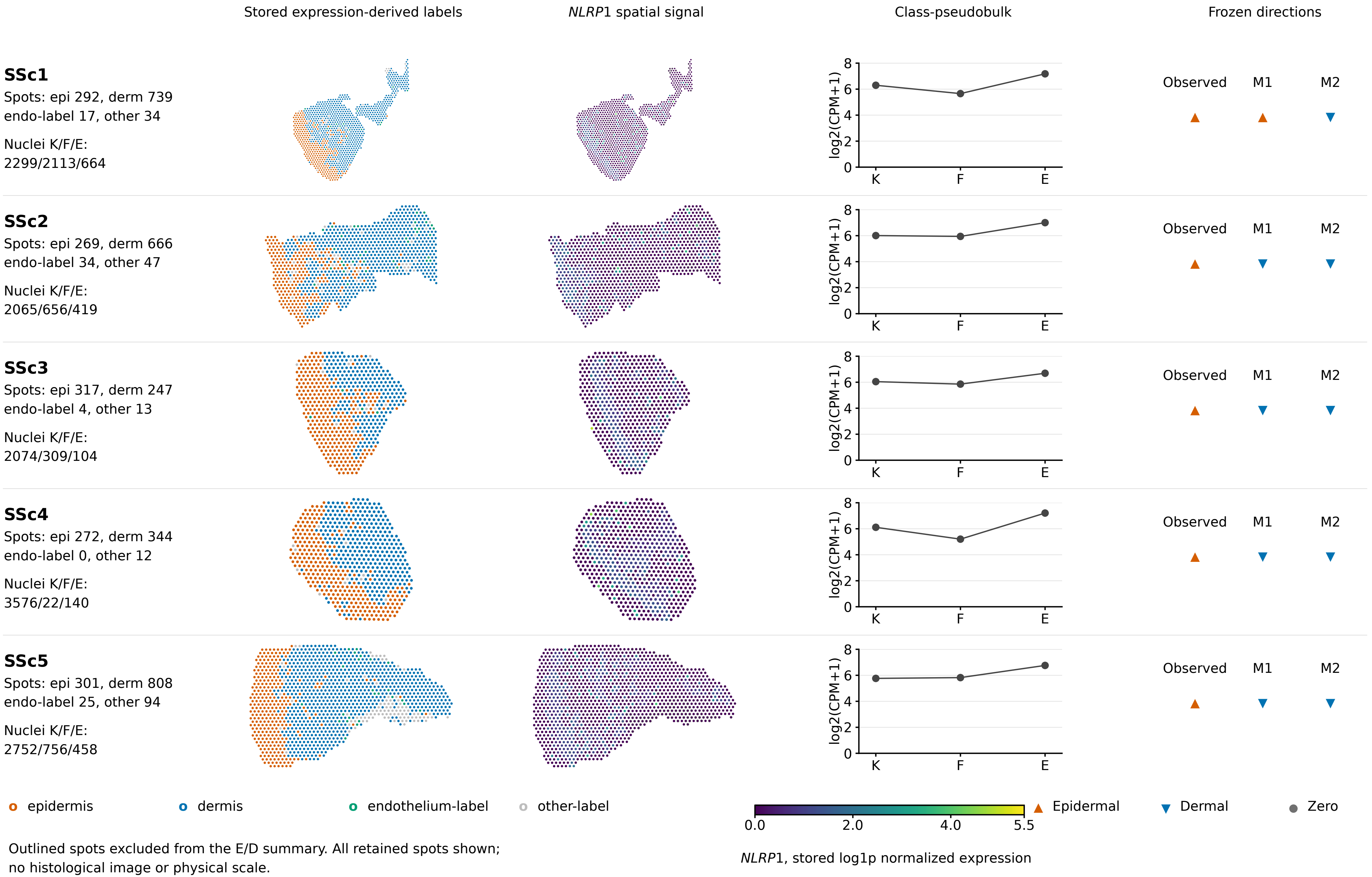


Outlined spots excluded from the E/D summary. All retained spots shown;
no histological image or physical scale.

K: keratinocyte; F: fibroblast; E: endothelial_strict. M1: donor F:E nucleus-ratio weights; M2: equal F:E weights.

Restricted proxy, not measured Visium composition. Directions only; spatial and class-pseudobulk scales are distinct.

### *NLRP*1 spatial / cell-class atlas: all 10 donors

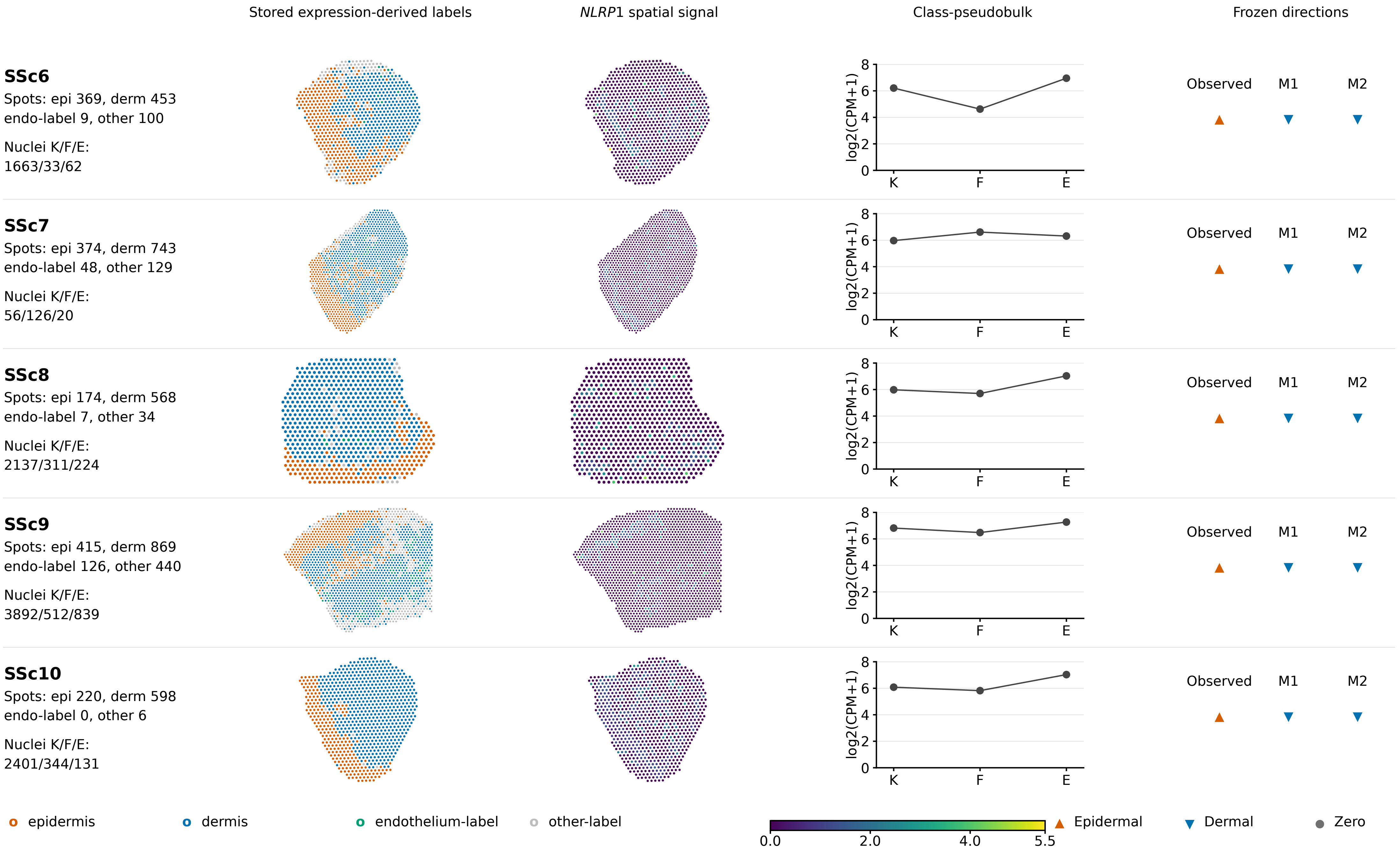


Outlined spots excluded from the E/D summary. All retained spots shown; no histological image or physical scale.

K: keratinocyte; F: fibroblast; E: endothelial_strict. M1: donor F:E nucleus-ratio weights; M2: equal F:E weights.

Restricted proxy, not measured Visium composition. Directions only; spatial and class-pseudobulk scales are distinct.
