## Supplementary Software 1 for "Layer-to-cell-class correspondence is gene-specific for pyroptosis-related transcription in human skin": final_figure_content.pdf

### *NLRP1* spatial / cell-class atlas: all 10 donors

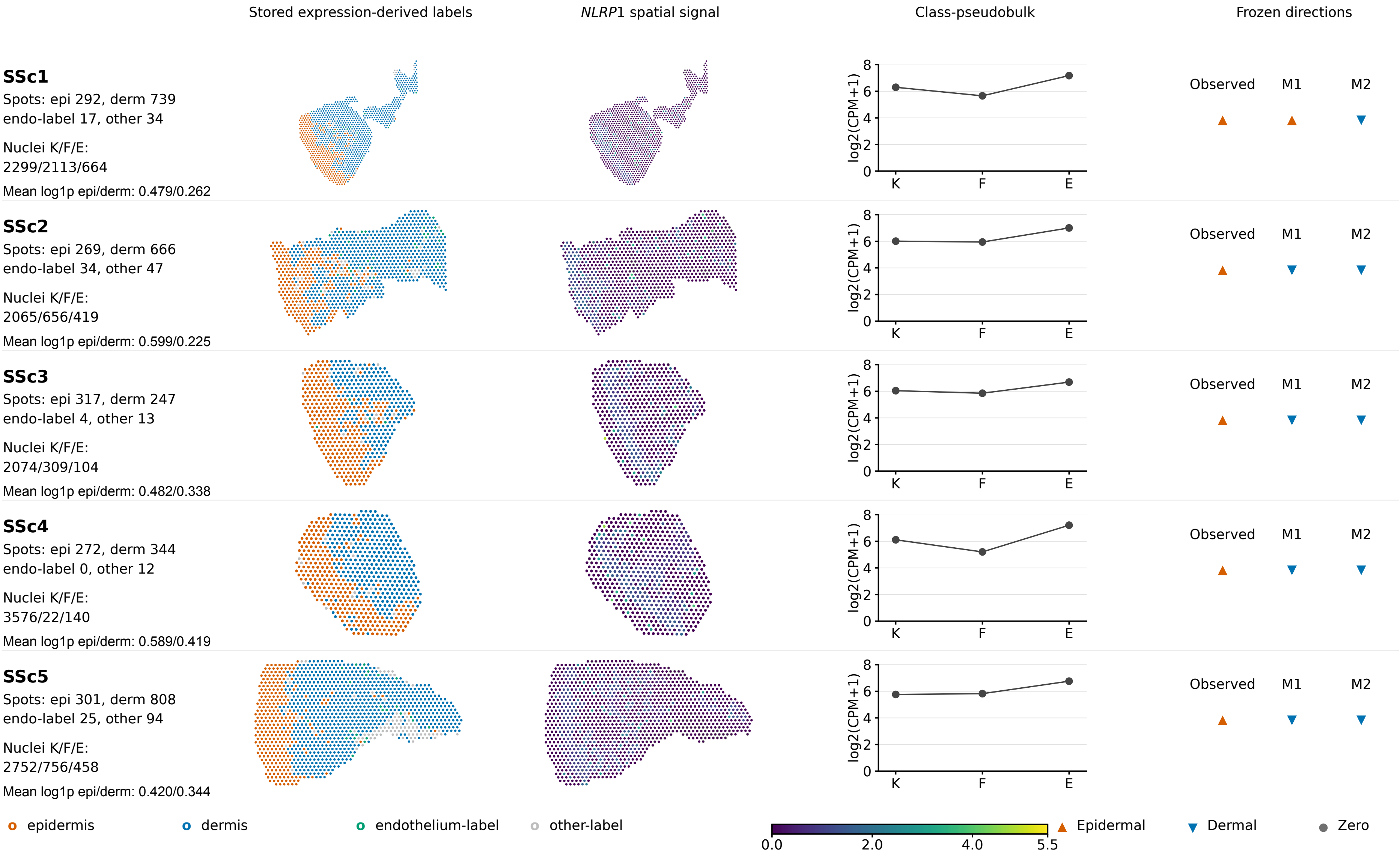


Outlined spots excluded from the E/D summary. All retained spots shown;
no histological image or physical scale.

K: keratinocyte; F: fibroblast; E: endothelial_strict. M1: donor F:E nucleus-ratio weights; M2: equal F:E weights.

Restricted proxy, not measured Visium composition. Directions only; spatial and class-pseudobulk scales are distinct.

### *NLRP1* spatial / cell-class atlas: all 10 donors



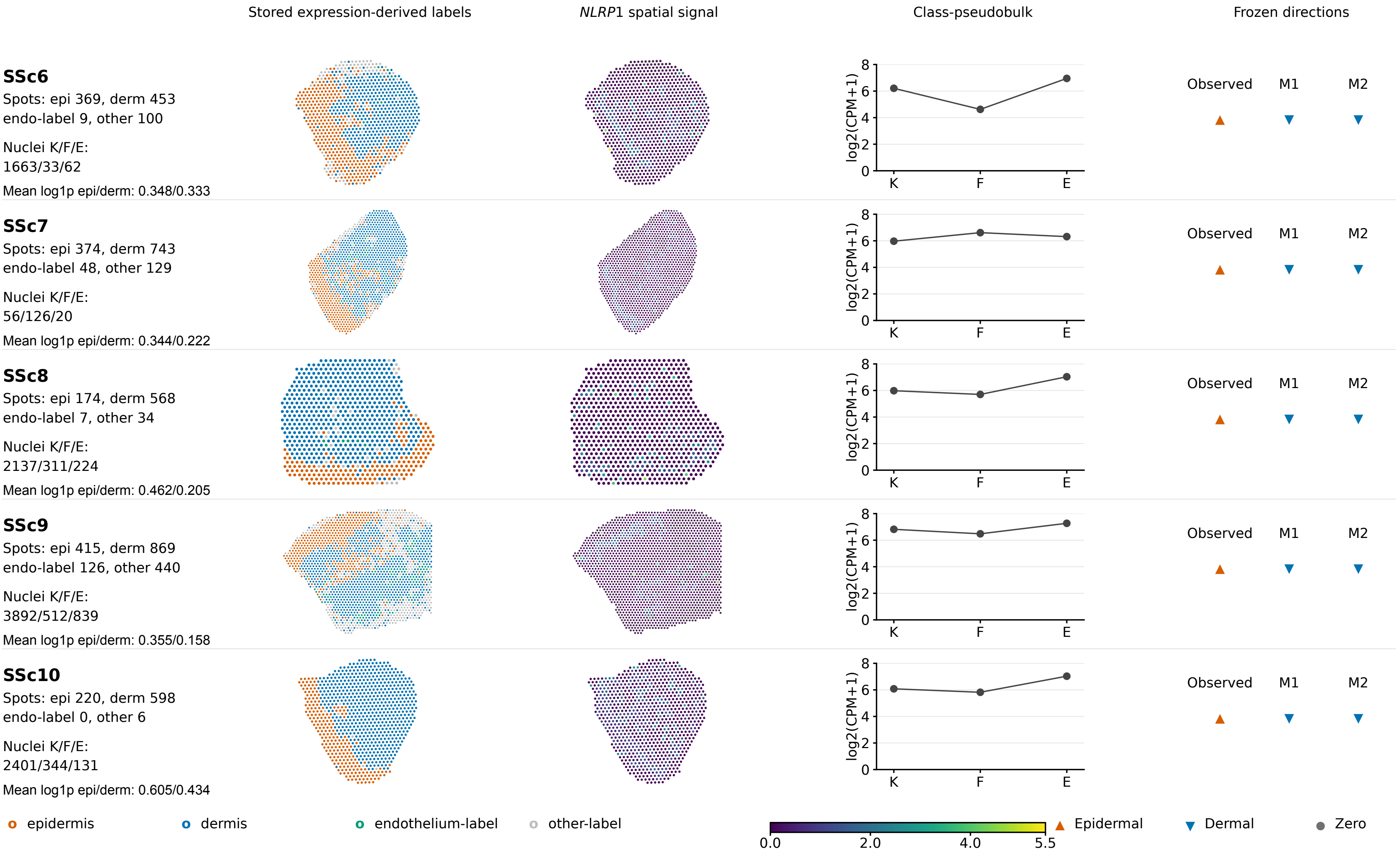


o epidermis o dermis o endothelium-label o other-label

0.0 2.0 4.0 5.5 ▲ Epidermal ▼ Dermal ● Zero

*NLRP1*, stored log1p normalized expression

Outlined spots excluded from the E/D summary. All retained spots shown; no histological image or physical scale.

K: keratinocyte; F: fibroblast; E: endothelial_strict. M1: donor F:E nucleus-ratio weights; M2: equal F:E weights.

Restricted proxy, not measured Visium composition. Directions only; spatial and class-pseudobulk scales are distinct.
