## Supplementary Software 1 for "Layer-to-cell-class correspondence is gene-specific for pyroptosis-related transcription in human skin": render.py

```
import argparse,csv,json
from pathlib import Path
import numpy as np
import matplotlib
matplotlib.use('Agg')
import matplotlib.pyplot as plt
from matplotlib.backends.backend_pdf import PdfPages
from matplotlib.collections import PatchCollection
from matplotlib.patches import Circle
from matplotlib.colors import Normalize
from matplotlib.lines import Line2D
from scipy.spatial import cKDTree
import fitz

COLORS = {'epidermis': '#D55E00', 'dermis': '#0072B2',
          'endothelium': '#009E73', 'other': '#BDBDBD'}
CLASSES = ['keratinocyte', 'fibroblast', 'endothelial_strict']
MODELS = ['M1_DONOR_F_E_NUCLEUS_RATIO', 'M2_EQUAL_F_E']
DONORS = ['SSc' + str(i) for i in range(1, 11)]
NORM = Normalize(0, 5.5)
GENE = r'$\mathit{NLRP1}$'
plt.rcParams.update({'font.family': 'DejaVu Sans', 'font.size': 8,
                     'axes.labelsize': 8, 'axes.titlesize': 10,
                     'xtick.labelsize': 8, 'ytick.labelsize': 8,
                     'legend.fontsize': 8, 'pdf.fonttype': 42,
                     'ps.fonttype': 42, 'mathtext.fontset': 'dejavusans',
                     'axes.spines.top': False, 'axes.spines.right': False,
                     'savefig.facecolor': 'white'})

def axes_mm(fig, x, y, w, h):
    fw, fh = fig.get_size_inches() * 25.4
    return fig.add_axes([x / fw, 1 - (y + h) / fh, w / fw, h / fh])

def text_mm(fig, x, y, value, **kwargs):
    fw, fh = fig.get_size_inches() * 25.4
    return fig.text(x / fw, 1 - y / fh, value, va='top', **kwargs)

def spatial_map(ax, data, expression=False):
    circles = [Circle((x, y), data['radius']) for x, y in data['xy']]
    if expression:
        collection = PatchCollection(circles, cmap='viridis', norm=NORM, alpha=1, linewidth=0)
        collection.set_array(data['values'])
    else:
        collection = PatchCollection(circles, facecolors=[COLORS[x] for x in data['labels']], linewidth=0, alpha=1)
    ax.add_collection(collection)
    if expression:
        excluded = ~np.isin(data['labels'], ['epidermis', 'dermis'])
        outlines = [Circle(tuple(c), data['radius']) for c in data['xy'][excluded]]
        ax.add_collection(PatchCollection(outlines, facecolors='none', edgecolors='#828282', linewidth=0.22))
    x0, y0, x1, y1 = data['bounds']
    ax.set_xlim(x0, x1)
    ax.set_ylim(y1, y0)
    ax.set_aspect('equal')
    ax.axis('off')
    return collection

def label_legend(fig, x, y, columns=2):
    names = ['epidermis', 'dermis', 'endothelium-label', 'other-label']
    for i, (key, name) in enumerate(zip(COLORS, names)):
        xx, yy = x + (i % columns) * 36, y + (i // columns) * 4
        text_mm(fig, xx, yy, 'o', color=COLORS[key], fontweight='bold')
        text_mm(fig, xx + 4, yy, name)

def class_axis(ax):
    ax.set_xlim(-0.2, 2.2)
    ax.set_ylim(0, 8)
    ax.set_xticks([0, 1, 2], ['K', 'F', 'E'])
    ax.set_yticks([0, 2, 4, 6, 8])
    ax.tick_params(length=2, pad=2)
    ax.grid(axis='y', color='#E1E1E1', linewidth=0.4)
    ax.set_axisbelow(True)

def direction_matrix(ax, selected, row_labels=True):
    ax.set_xlim(-0.5, 2.5)
    ax.set_ylim(len(selected) - 0.5, -0.5)
    for i, row in enumerate(selected):
        for j, key in enumerate(('observed_sign', 'M1_predicted_sign', 'M2_predicted_sign')):
            sign = int(row[key])
            marker, color = {1: ('^', COLORS['epidermis']), -1: ('v', COLORS['dermis']), 0: ('o', '#707070')}[sign]
            ax.scatter(j, i, marker=marker, s=32, c=color, edgecolors='none')
        if len(selected) > 1:
            ax.axhline(i + 0.5, color='#E5E5E5', lw=0.4)
    ax.set_xticks([0, 1, 2], ['Observed', 'M1', 'M2'])
    ax.xaxis.tick_top()
    ax.set_yticks(range(len(selected)), [r['report_donor'] for r in selected] if row_labels else [''] * len(selected))
    ax.tick_params(length=0, pad=4)
    for spine in ax.spines.values():
        spine.set_visible(False)

def direction_key(fig, x, y):
    for i, (mark, color, label) in enumerate([('^', COLORS['epidermis'], 'Epidermal'), ('v', COLORS['dermis'], 'Dermal'), ('o', '#707070', 'Zero')]):
        xx = x + i * 27
        fw, fh = fig.get_size_inches() * 25.4
        fig.add_artist(Line2D([xx / fw], [1 - (y + 1.7) / fh], transform=fig.transFigure,
                              marker=mark, color=color, markersize=4, linestyle='none'))
        text_mm(fig, xx + 3, y, label)

def atlas_figure(display, maps, number):
    fig = plt.figure(figsize=(297 / 25.4, 210 / 25.4))
    text_mm(fig, 4, 3, GENE + ' spatial / cell-class atlas: all 10 donors', fontsize=11, fontweight='bold')
    text_mm(fig, 293, 4, f'{number}/2', ha='right')
    text_mm(fig, 78, 13, 'Stored expression-derived labels', ha='center')
    text_mm(fig, 138, 13, GENE + ' spatial signal', ha='center')
    text_mm(fig, 205, 13, 'Class-pseudobulk', ha='center')
    text_mm(fig, 271, 13, 'Frozen directions', ha='center')
    selected = display[(number - 1) * 5:number * 5]
    for i, row in enumerate(selected):
        y = 25 + i * 31
        donor = row['report_donor']
        text_mm(fig, 4, y + 1, donor, fontsize=10, fontweight='bold')
        text_mm(fig, 4, y + 6, f"Spots: epi {row['n_epidermis']}, derm {row['n_dermis']}")
        text_mm(fig, 4, y + 10, f"endo-label {row['n_endothelium']}, other {row['n_other']}")
        text_mm(fig, 4, y + 16, 'Nuclei K/F/E:')
        text_mm(fig, 4, y + 20, '/'.join(row[k + '_n_nuclei'] for k in ('K', 'F', 'E')))
        spatial_map(axes_mm(fig, 53, y - 2, 49, 28), maps[donor])
        image = spatial_map(axes_mm(fig, 113, y - 2, 49, 28), maps[donor], True)
        ax = axes_mm(fig, 185, y, 43, 22)
        class_axis(ax)
        vals = [float(row[k + '_log2_CPM_plus_1']) for k in ('K', 'F', 'E')]
        ax.plot([0, 1, 2], vals, marker='o', markersize=3.5, color='#454545', lw=0.8)
        ax.set_ylabel('log2(CPM+1)', labelpad=1)
        direction_matrix(axes_mm(fig, 249, y + 7, 43, 9), [row], row_labels=False)
        if i < 4:
            fig.add_artist(Line2D([0.013, 0.985], [1 - (y + 28) / 210] * 2,
                                  transform=fig.transFigure, lw=0.4, color='#DDDDDD'))
    label_legend(fig, 5, 181, columns=4)
    cb = fig.colorbar(image, cax=axes_mm(fig, 163, 182, 57, 2),
                      orientation='horizontal', ticks=[0, 2, 4, 5.5])
    cb.ax.tick_params(length=2, pad=1)
    text_mm(fig, 160, 192, GENE + ', stored log1p normalized expression')
    direction_key(fig, 223, 181)
    text_mm(fig, 5, 190, 'Outlined spots excluded from the E/D summary. All retained spots shown;')
    text_mm(fig, 5, 194, 'no histological image or physical scale.')
    text_mm(fig, 5, 199, 'K: keratinocyte; F: fibroblast; E: endothelial_strict. M1: donor F:E nucleus-ratio weights; M2: equal F:E weights.')
    text_mm(fig, 5, 204, 'Restricted proxy, not measured Visium composition. Directions only; spatial and class-pseudobulk scales are distinct.')
    return fig

def rows(path):
    with path.open(newline='') as f:return list(csv.DictReader(f,delimiter='\t'))

def load_display():
    data=Path(__file__).parent/'data';display=rows(data/'donor_display_values.tsv');spots=rows(data/'spot_values.tsv')
    assert len(spots)==10217 and [r['report_donor'] for r in display]==DONORS
    maps={}
    for r in display:
        selected=[s for s in spots if s['visium_sample_id']==r['visium_sample_id']]
        coords=np.array([[int(s['spatial_1']),int(s['spatial_0'])] for s in selected],dtype=float)
        labels=np.array([s['stored_layer'] for s in selected]);values=np.array([float(s['NLRP1_X_log1p']) for s in selected])
        nearest=cKDTree(coords).query(coords,k=2)[0][:,1];radius=.30*float(np.median(nearest))
        lo,hi=coords.min(axis=0),coords.max(axis=0);pad=(hi-lo)*.05
        maps[r['report_donor']]={'xy':coords,'labels':labels,'values':values,'radius':radius,'bounds':[*lo-pad,*hi+pad]}
    return display,maps

def overlay_means(native,output,font):
    doc=fitz.open(native);values=rows(Path(__file__).parent/'data/S14_LAYER_MEANS_20_VALUES.tsv')
    assert len(values)==20
    for i,donor in enumerate(DONORS):
        page=doc[i//5]
        headers=[s for b in page.get_text('dict')['blocks'] if 'lines' in b for line in b['lines'] for s in line['spans'] if s['text']==donor]
        assert len(headers)==1
        rr=[r for r in values if r['report_donor']==donor]
        label=rr[0]['label'];assert all(r['label']==label for r in rr)
        y=headers[0]['origin'][1]+66.5
        page.insert_text((11.338583,y),label,fontsize=8,fontname='LayerMeans',fontfile=str(font),color=(0,0,0))
    doc.save(output,deflate=True);doc.close()

if __name__=='__main__':
    p=argparse.ArgumentParser();p.add_argument('--output',type=Path,required=True);p.add_argument('--font',type=Path,required=True);a=p.parse_args();a.output.mkdir(parents=True,exist_ok=True)
    display,maps=load_display()
    with PdfPages(a.output/'atlas_native.pdf') as pdf:
        for number in (1,2):
            fig=atlas_figure(display,maps,number);fig.canvas.draw();pdf.savefig(fig);plt.close(fig)
    overlay_means(a.output/'atlas_native.pdf',a.output/'atlas_with_saved_means.pdf',a.font)
    ledger=[]
    for donor,data in maps.items():
        for xy,label,val in zip(data['xy'],data['labels'],data['values']):
            ledger.append({'donor':donor,'x':float(xy[0]),'y':float(xy[1]),'layer':str(label),'value':float(val),'radius':data['radius']})
    (a.output/'SPOT_DRAWING_LEDGER.json').write_text(json.dumps(ledger,separators=(',',':'))+'\n')
```
