## Supplementary figures and images for "Layer-to-cell-class correspondence is gene-specific for pyroptosis-related transcription in human skin"

### Figure_5.tif

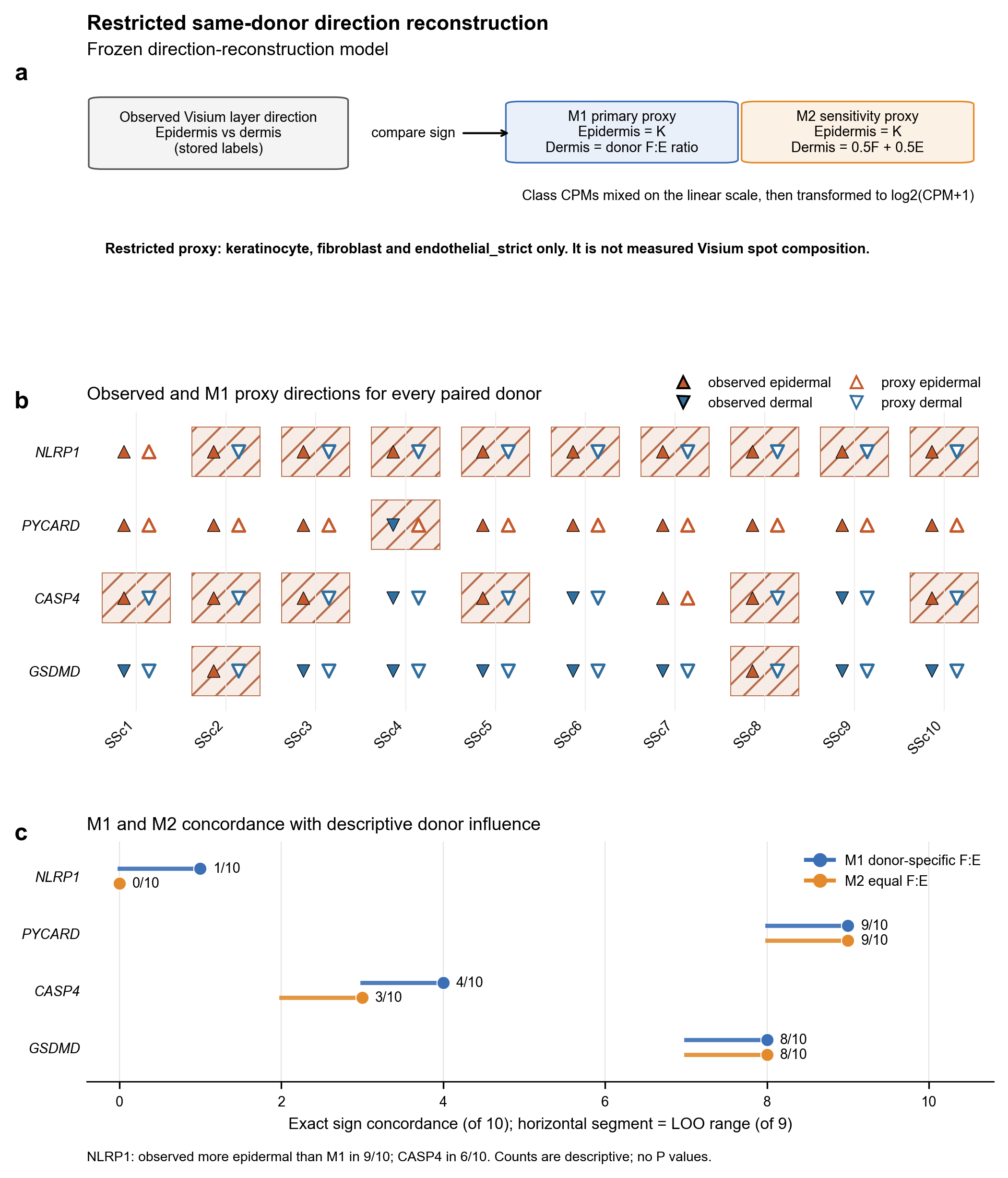

### S08_parent.png

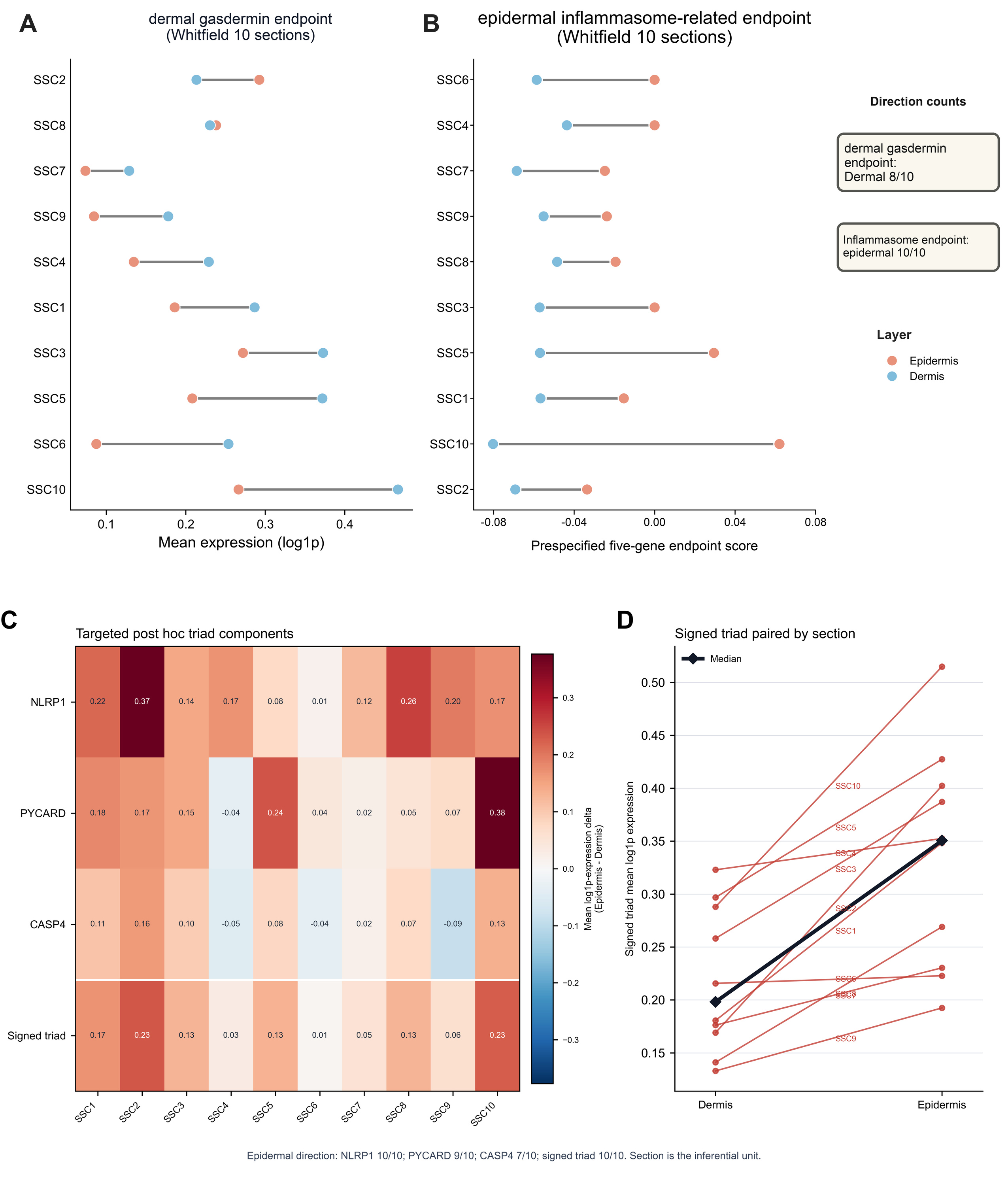

### S08_replacement.png

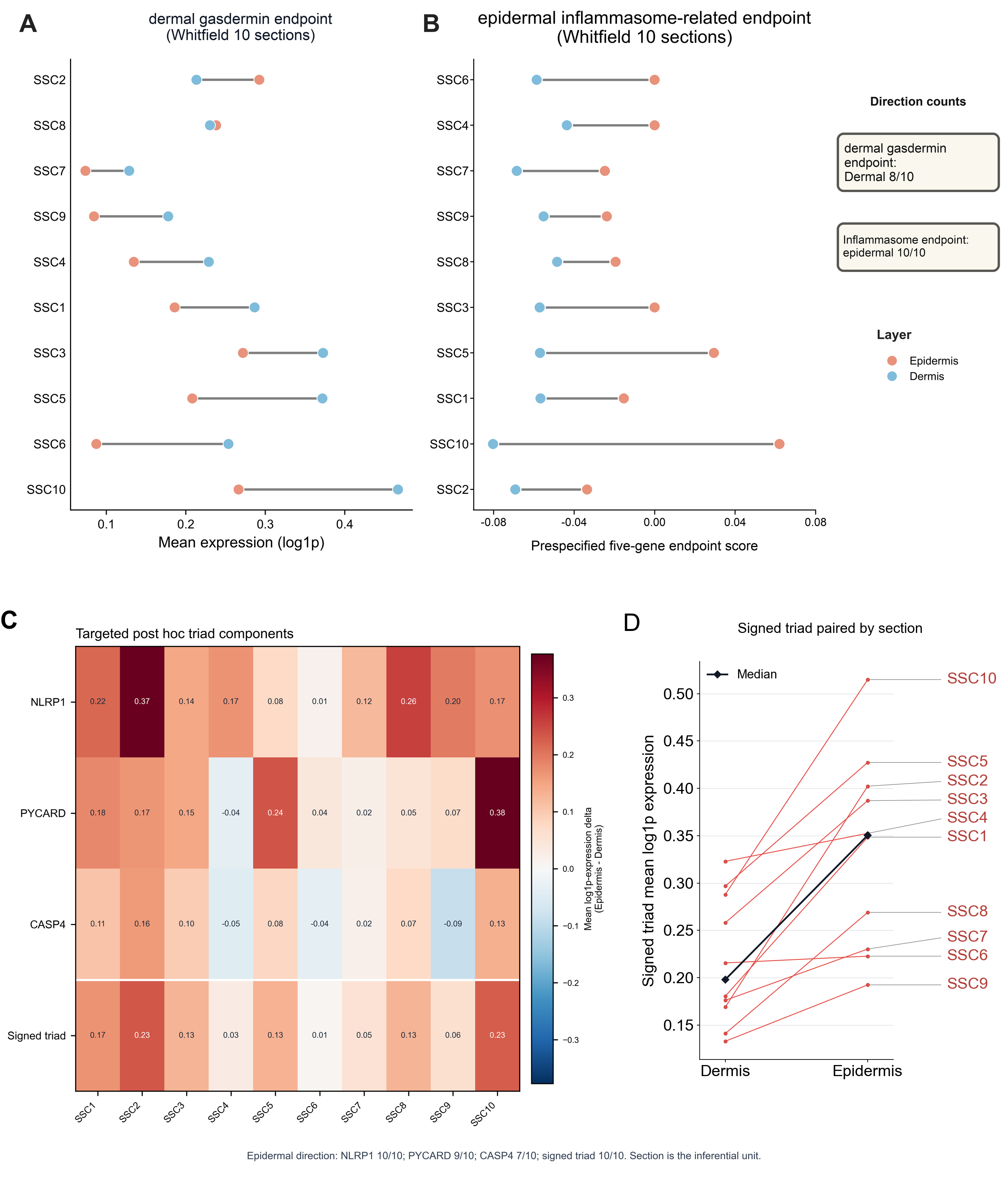

### S8D.png

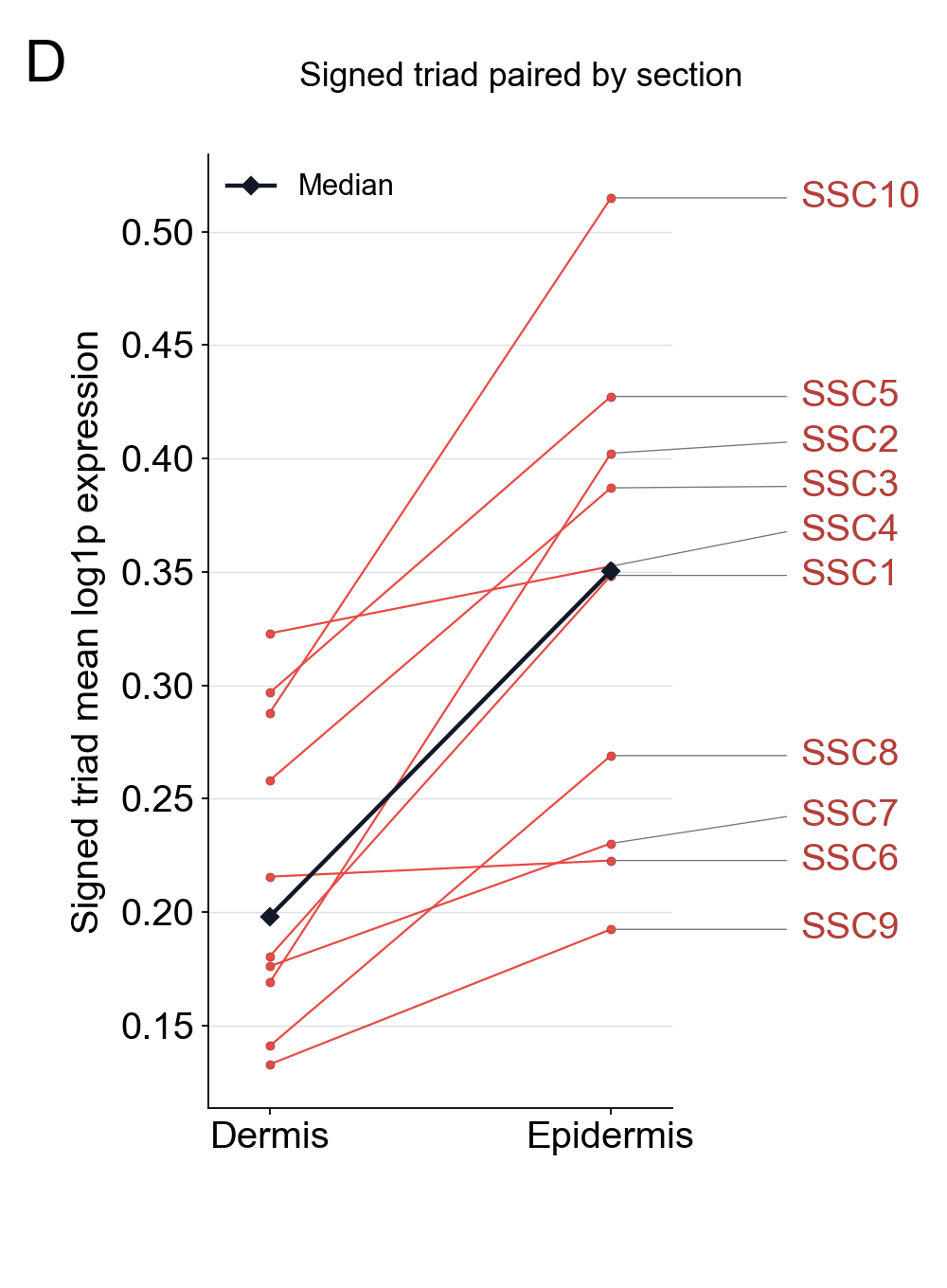

### S10_parent.png

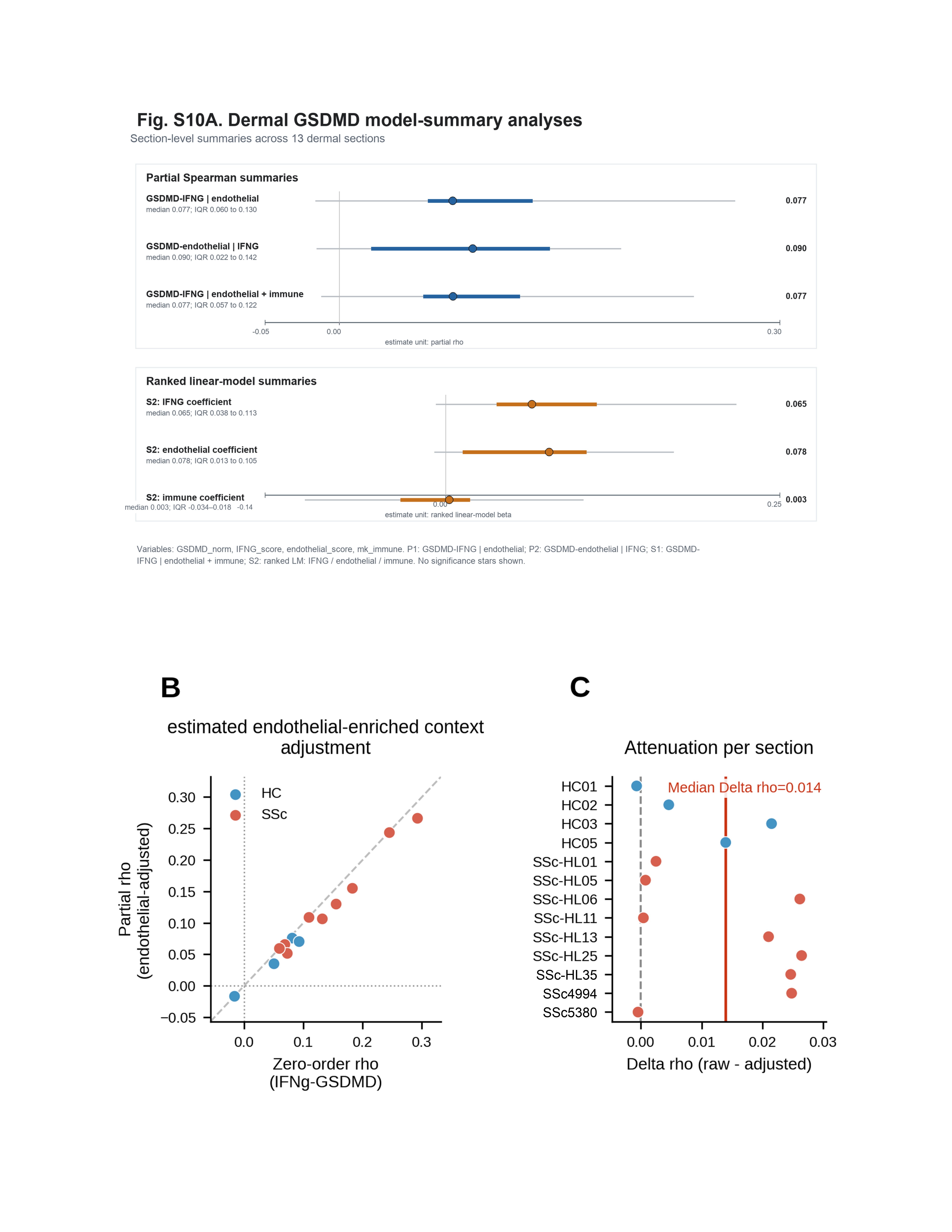

### S10_replacement.png

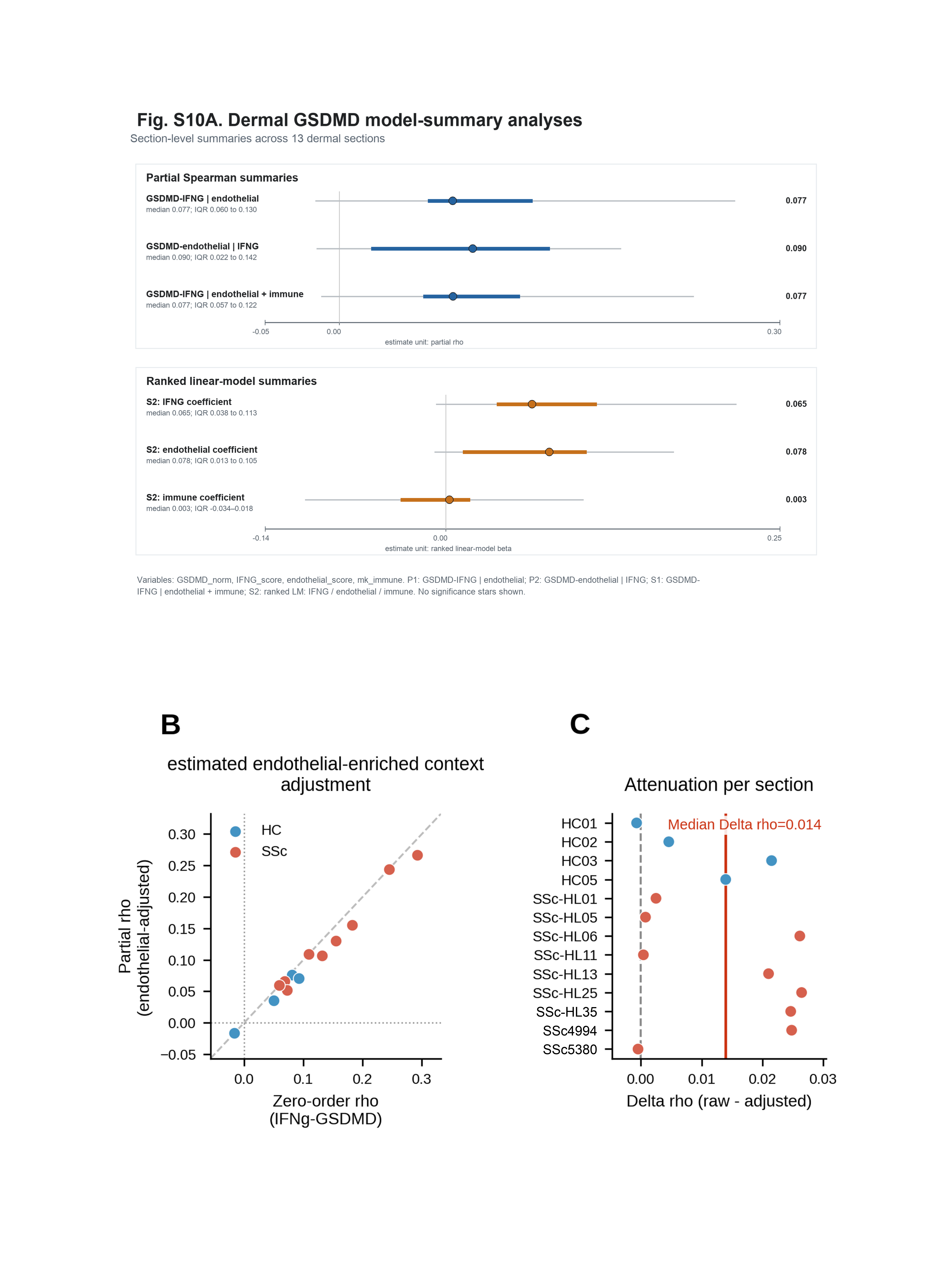

### S10A_replacement_native.png

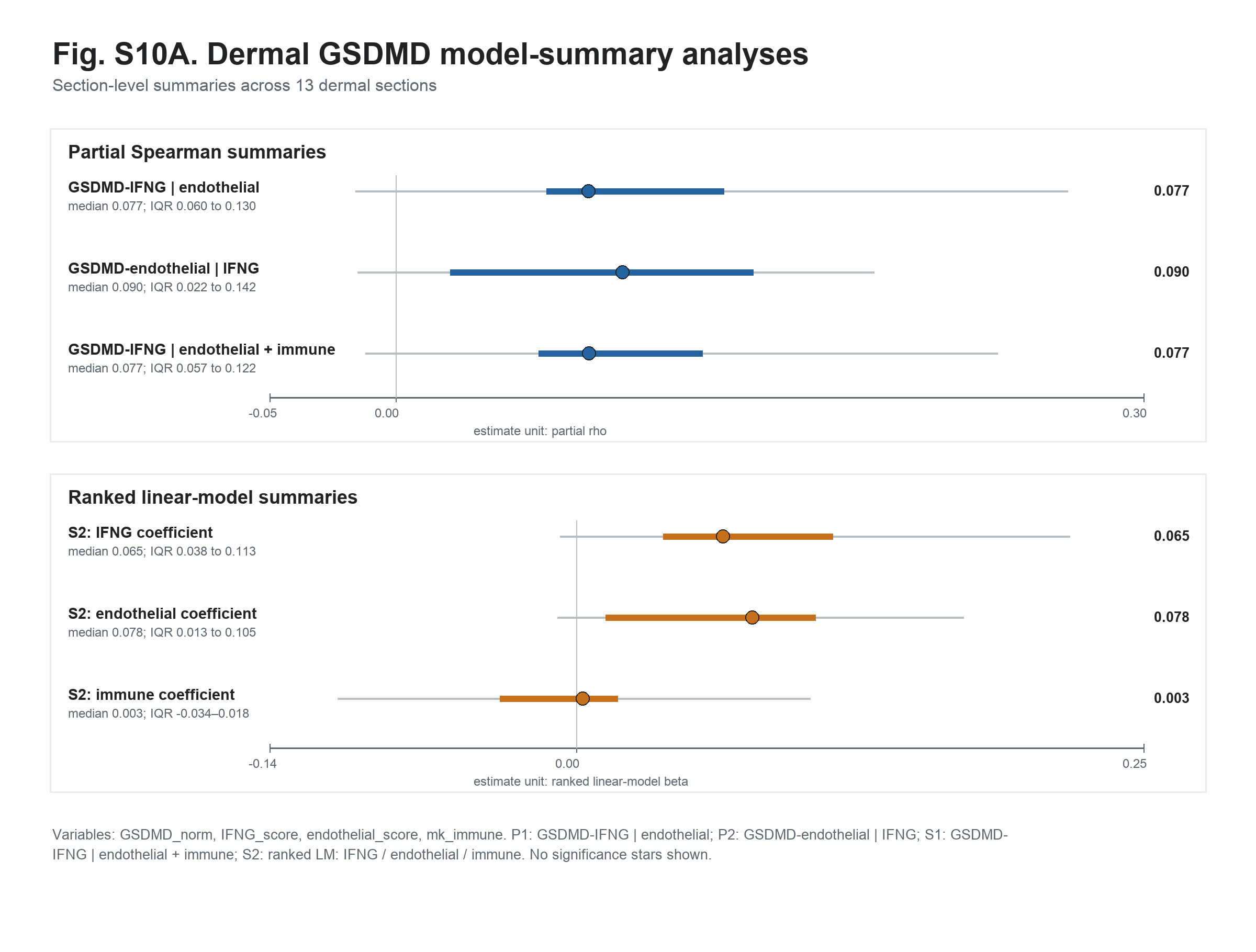
